# Open issues for protein function assignment in *Haloferax volcanii* and other halophilic archaea

**DOI:** 10.1101/2021.05.03.442417

**Authors:** Friedhelm Pfeiffer, Mike Dyall-Smith

## Abstract

**Background:** Annotation ambiguities and annotation errors are a general challenge in genomics. While a reliable protein function assignment can be obtained by experimental characterization, this is expensive and time-consuming, and the number of such Gold Standard Proteins (GSP) with experimental support remains very low compared to proteins annotated by sequence homology, usually through automated pipelines. Even a GSP may give a misleading assignment when used as a reference: the homolog may be close enough to support isofunctionality, but the substrate of the GSP is absent from the species being annotated. In such cases the enzymes cannot be isofunctional. Here, we examine a variety of such issues in halophilic archaea (class Halobacteria), with a strong focus on the model haloarchaeon *Haloferax volcanii*.

**Results:** Annotated proteins of *Hfx. volcanii* were identified for which public databases tend to assign a function that is probably incorrect. In some cases, an alternative, probably correct, function can be predicted or inferred from the available evidence but this has not been adopted by public databases because experimental validation is lacking. In other cases, a probably invalid specific function is predicted by homology, and while there is evidence that this assigned function is unlikely, the true function remains elusive. We list 50 of those cases, each with detailed background information so that a conclusion about the most likely biological function can be drawn. For reasons of brevity and comprehension, only key aspects are listed in the main text, with detailed information being provided in a corresponding section of the Supplementary Material.

**Conclusions:** Compiling, describing and summarizing these open annotation issues and functional predictions will benefit the scientific community in the general effort to improve the evaluation of protein function assignments and more thoroughly detail them. By highlighting the gaps and likely annotation errors currently in the databases, we hope this study will provide a framework for experimentalists to systematically confirm (or disprove) our function predictions or to uncover yet unexpected functions.

## 1 INTRODUCTION

*Haloferax volcanii* is a model organism for halophilic archaea (Hartman et al., 2010, Schulze et al., 2020, Leigh et al., 2011, Perez-Arnaiz et al., 2020, Soppa, 2011, Haque et al., 2020), for which an elaborate set of genetic tools has been developed (Allers et al., 2010, Allers and Mevarech, 2005, Kiljunen et al., 2014). Its genome has been sequenced and carefully annotated (Hartman et al., 2010, Pfeiffer et al., 2008a, Pfeiffer and Oesterhelt, 2015). A plethora of biological aspects have been successfully tackled in this species, with examples including DNA replication (Perez-Arnaiz et al., 2020); cell division and cell shape (Turkowyd et al., 2020, Walsh et al., 2019, de Silva et al., 2021, Duggin et al., 2015, Liao et al., 2021); metabolism (Brasen and Schonheit, 2001, Johnsen et al., 2009, Pickl et al., 2012, Sutter et al., 2016, Reinhardt et al., 2019, Kuprat et al., 2021, Kuprat et al., 2020, Sutter et al., 2020, Tästensen et al., 2020); protein secretion (Abdul-Halim et al., 2020, Abdul Halim et al., 2018, Abdul Halim et al., 2013, Storf et al., 2010); motility and biofilms (Schiller et al., 2020, Pohlschroder and Esquivel, 2015, Li et al., 2020, Collins et al., 2020, Quax et al., 2018, Nussbaum et al., 2020); mating (Shalev et al., 2017); signalling (Braun et al., 2019); virus defence (Maier et al., 2019); proteolysis (Reuter and Maupin-Furlow, 2004, Reuter et al., 2010, Prunetti et al., 2014, Cerletti et al., 2018, Cerletti et al., 2014, Costa et al., 2018); posttranslational modification (N-glycosylation; SAMPylation)(Cao et al., 2015, Kaminski and Eichler, 2014, Tripepi et al., 2012, Schulze et al., 2021, Shalev et al., 2018, Kandiba et al., 2016); gene regulation (Qi et al., 2016, Rawls et al., 2010, Hattori et al., 2016, Hwang et al., 2017, Johnsen et al., 2015, Tästensen et al., 2020, Reinhardt et al., 2019); microproteins (Zahn et al., 2021, Nagel et al., 2019, Kubatova et al., 2020) and small noncoding RNAs (sRNAs) (Straub et al., 2009, Heyer et al., 2012, Babski et al., 2014, Wyss et al., 2018).

Genome annotations are frequently compromised by annotation errors (Schnoes et al., 2009, Pfeiffer and Oesterhelt, 2015, Promponas et al., 2015, Danchin et al., 2018). Many of these errors are caused by invalid annotation transfer between presumed homologs, which, once introduced, are further spread by annotation robots. This problem can be partially overcome by using a Gold Standard Protein (GSP) based annotation strategy (Pfeiffer and Oesterhelt, 2015). Because the GSP has itself been subjected to experimental analysis, its annotation cannot be caused by an invalid annotation transfer process. The GSP strategy had already been applied to a detailed analysis of the metabolism of halophilic archaea (Falb et al., 2008). However, with decreasing level of sequence identity, the assumption of isofunctionality becomes increasingly insecure. Although this may be counterbalanced by additional evidence, e.g. gene clustering, experimental confirmation would be the best option for validation of the annotation.

There are additional and much more subtle genome annotation problems. In some cases, GSPs are true homologs and the annotated function in the database is correct. Nevertheless, the biological context in the query organism makes it unlikely that the homologs are isofunctional, e.g. when the substrate of the GSP is lacking in the query organism. Also, paralogs may have distinct but related functions, which cannot be assigned by sequence analysis but may be assigned based on phylogenetic considerations. Here, again, experimental confirmation is the preferred option for validation of the annotation. Lack of experimental confirmation may keep high-level databases like KEGG or the SwissProt section of UniProt from adopting assignments based on well-supported bioinformatic analyses, so that the database entries continue to provide information that is probably incorrect. We refer to annotation problems in these databases solely to underscore that the biological issues raised by us are far from trivial. There is no intention to question the exceedingly high quality of the SwissProt and KEGG databases (UniProt, 2021, Kanehisa et al., 2019) and their tremendous value for the scientific community. We have actively supported them by providing feedback and encourage others to do the same, e.g. with the recently implemented “Add a publication” functionality in UniProt entries, that allows users to connect a protein to a publication that describes its experimental characterization (https://community.uniprot.org/bbsub/bbsubinfo.html).

In this study, we describe a number of annotation issues for haloarchaea with a strong emphasis on *Hfx. volcanii*. We denote such cases as ‘open annotation issues’ with the hope of attracting members of the *Haloferax* community and other groups working with halophilic archaea to apply experimental analyses to elucidate the true function(s) of these proteins. This will increase the number of Gold Standard Proteins which originate from *Hfx. volcanii* or other haloarchaea, reduce genome annotation ambiguities, and perhaps uncover novel metabolic processes.

## 2 MATERIALS AND METHODS

### 2.1 Curation of genome annotation and Gold Standard Protein identification

The Gold Standard Protein based curation of haloarchaeal genomes has been described (Pfeiffer and Oesterhelt, 2015). Since then, a systematic comparison to KEGG data was performed for a subset of the curated genomes (Pfeiffer et al., 2020). The *Hfx. volcanii* genome annotation is continuously scrutinized, especially when a closely related genome is annotated (Tittes et al., 2021). The 16 haloarchaeal genomes that are currently under survey are listed in Suppl. Table S11.

### 2.2 Additional bioinformatics tools

Key databases were UniProtKB/Swiss-Prot (UniProt, 2021), InterPro (Hunter et al., 2009), KEGG (Kanehisa et al., 2019), and OrthoDB (Kriventseva et al., 2019). The SyntTax server was used for inspecting conservation of gene neighbourhood analysis (Oberto, 2013). As general tools the BLAST suite of programs (Johnson et al., 2008, Altschul et al., 1997) was used for genome comparisons.

## 3 RESULTS

Open issues are organised below under the subsections (3.1), the respiratory chain and oxidative decarboxylation; (3.2), amino acid metabolism; (3.3), heme and cobalamin biosynthesis; (3.4), coenzyme F420; (3.5), tetrahydrofolate as opposed to methanopterin; (3.6), NAD and riboflavin; (3.7), lipid metabolism; (3.8), genetic information processing, and (3.9), stand-alone (miscellaneous) cases.

### 3.1 The respiratory chain and oxidative decarboxylation

In the respiratory chain, coenzymes that have been reduced during catabolism (e.g. glycolysis) are reoxidized, with the energy being saved as an ion gradient. The textbook example of a respiratory chain are the five mitochondrial complexes (Rich and Marechal, 2010, Guo et al., 2018): complex I (NADH dehydrogenase), complex II (succinate dehydrogenase), complex III (cytochrome bc_1_ complex), complex IV (cytochrome-c oxidase as prototype for a terminal oxidase) and complex V (F-type ATP synthase). In mitochondria, a significant part of the NADH which feeds into the respiratory chain originates from oxidative decarboxylation: conversion of pyruvate to acetyl-CoA by the pyruvate dehydrogenase complex and conversion of alpha-ketoglutarate to succinyl-CoA by the homologous 2-oxoglutarate dehydrogenase complex. While complex I and II transfer reducing elements to a lipid-embedded two-electron carrier (ubiquinone), the bc_1_ complex transfers the electrons to the one-electron carrier cytochrome-c, a heme (and thus iron) protein, which then transfers electrons to the terminal oxidase.

Bacteria like *Escherichia coli* and *Paracoccus* have related complexes and enzymes: NADH dehydrogenase (encoded by the *nuo* operon), succinate dehydrogenase (encoded by *sdhABCD*) and the related fumarate reductase (encoded by *frdABCD*) (Crofts et al., 2013), several terminal oxidases (e.g. products of *cyoABCDE*, *cydABC*), and an F-type ATP synthase (encoded by *atp* genes). *E. coli* lacks a bc_1_ complex, which, however, occurs in *Paracoccus denitrificans* (Kaila and Wikstrom, 2021). *E. coli* contains the canonical complexes of oxidative decarboxylation (pyruvate dehydrogenase complex, encoded by *aceEF*+*lpdA*, and 2-oxoglutarate dehydrogenase complex, encoded by *sucAB*+*lpdA*).

The respiratory chain of *Hfx. volcanii* and other haloarchaea deviates considerably from those of mitochondria and bacteria such as *Paracoccus* and *E. coli* (reviewed by (Schafer et al., 1999)), and a number of questions remain unresolved. We focus on the equivalents of complex I, III and IV, because these have unresolved issues. We also cover some aspects relevant for NADH levels (oxidative decarboxylation enzymes and type II NADH dehydrogenase). We do not cover complexes that have already been studied in haloarchaea: complex II (succinate dehydrogenase) (Scharf et al., 1997, Sreeramulu et al., 1998, Gradin et al., 1985) and complex V (ATP synthase) (Steinert et al., 1997, Nanba and Mukohata, 1987).

(a) In haloarchaea, oxidative decarboxylation is not linked to reduction of NAD to NADH but to reduction of a ferredoxin (encoded by *fdx*, e.g. OE_4217R, HVO_2995) which has a redox potential similar to that of the NAD/NADH pair (Kerscher and Oesterhelt, 1977). The enzymes for oxidative decarboxylation are pyruvate--ferredoxin oxidoreductase (*porAB*, e.g. OE_2623R/2622R, HVO_1305/1304) and 2-oxoglutarate--ferredoxin oxidoreductase (*korAB*, e.g. OE_1711R/1710R, HVO_0888/0887), and these have been characterized from *Halobacterium salinarum* (Plaga et al., 1992, Kerscher and Oesterhelt, 1981b, Kerscher and Oesterhelt, 1981a).
(b) In *Hfx. volcanii*, ferredoxin Fdx (HVO_2995) plays an essential role in nitrate assimilation (Zafrilla et al., 2011). It may well be involved in additional metabolic processes and it is yet unresolved how ferredoxin Fdx is reoxidised, but this might be achieved by the Nuo complex.
(c) The *nuo* cluster of haloarchaea resembles that of *E. coli*, with genes and gene order highly conserved, and just a few domain fissions and fusions. However, haloarchaea lack NuoEFG (Falb et al., 2005), which is a subcomplex mediating interaction with NADH (Leif et al., 1995, Braun et al., 1998). Thus, the haloarchaeal *nuo* complex is unlikely to function as NADH dehydrogenase, despite its annotation as such in KEGG (as of April 2021).
(d) Other catabolic enzymes generate NADH, which also must be reoxidized. Based on inhibitor studies in *Hbt. salinarum*, NADH is not reoxidized by a type I but rather by a type II NADH dehydrogenase (Sreeramulu et al., 1998). A gene has been assigned for *Natronomonas pharaonis* (Falb et al., 2008). However, for reasons detailed in Suppl.Text.S1, this assignment is highly questionable, so that this issue calls for experimental analysis.
(e) About one-third of the haloarchaea, especially the *Natrialbales*, do not code for a complex III equivalent (cytochrome bc_1_ complex encoded by *petABC*) according to OrthoDB analysis. The bc_1_ complex is required to transfer electrons from the lipid-embedded two-electron carrier (menaquinone in haloarchaea) to the one-electron carrier associated with terminal oxidases (probably halocyanin). How electrons flow in the absence of a complex III equivalent is currently unresolved.

The haloarchaeal *petABC* genes resemble those of the chloroplast b6-f complex rather than those of the mitochondrial bc_1_ complex (see Suppl.Text S1 Section 1 for more details).

(f) A bc cytochrome has been purified from *Nmn. pharaonis*, but with an atypical 1:1 ratio between the b-type and c-type heme (Scharf et al., 1997). The complex is heterodimeric, with subunits of 18 kDa and 14 kDa. The 18 kDa subunit carries the covalently attached heme group (Scharf et al., 1997). An attempt was made to identify the genes coding for these subunits (Mattar, 1996) (for details see Suppl.Text S1 Section 1). Two approaches were used to obtain protein sequence data, one being N-terminal protein sequencing of the two subunits extracted from a SDS-polyacrylamide gel. In the other attempt, peptides from the purified complex were separated by HPLC, and a peptide absorbing at 280 nm (protein) as well as 400 nm (heme) was isolated. Absorption at 400 nm clearly indicates covalent attachment of the heme group to the peptide. The sequences from the two approaches overlapped and resulted in a contiguous sequence of 41 aa, with only the penultimate position remaining undefined (Mattar, 1996). Based on this information, a PCR probe was generated (designated “cyt-C Sonde”) that allowed the gene to be identified and sequenced, including its genomic neighbourhood. It turned out that the genes coding for the four subunits of succinate dehydrogenase (*sdhCDBA*) had been isolated. The obtained protein sequence corresponds to the N-terminal region of *sdhD* (with the initiator methionine cleaved off) and only 2 sequence discrepancies in addition to the unresolved penultimate residue.

In the PhD thesis (Mattar, 1996), this unambiguous result was rated to be a failure (and the data were never formally published). The reason is that SdhD is free of cysteine residues, while textbook knowledge states that a pair of cysteines is required for covalent heme attachment (Kletzin et al., 2015). The lack of the required cysteine pair was taken to indicate that the results were incorrect and that the identified genes did not encode the cytochrome bc that the study had been seeking (Mattar, 1996). In contrast, we speculate that the results were completely correct, despite being in conflict with the cysteine pair paradigm. In our view, a paradigm shift is required. The obtained results call for a yet unanticipated novel mode of covalent heme attachment, exemplified by the 18 kDa subunit of *Natronomonas* succinate dehydrogenase subunit SdhD. It should be noted that the 41 aa protein sequence, which had been obtained, turned out to contain three histidine residues upon translation of the gene, but none of these had been detected upon Edman degradation.

In *Halobacterium*, a small c-type cytochrome was purified (cytochrome c_552_, 14.1 kDa) (Sreeramulu, 2003). Heme staining after SDS-PAGE indicated a covalent heme attachment, but no sequence or composition data were reported, so that it is not possible to identify the protein based on the available information. We speculate that the *Halobacterium* cytochrome c_552_ also represents SdhD (as detailed in Suppl.Text S1 Section 1). In that case, the proposed novel type of covalent heme attachment would not be restricted to *Nmn. pharaonis* but might be a general property of haloarchaea. This would also solve the “*Halobacterium* paradox” (Kletzin et al., 2015).

(g) The haloarchaeal one-electron carrier is the copper protein halocyanin rather than the iron-containing heme protein cytochrome-c. A halocyanin from *Nmn. pharaonis* (NP_3954A) has been characterized, including its redox potential (Mattar et al., 1994, Scharf and Engelhard, 1993, Hildebrandt et al., 1994). A gene fusion supports the close connection of a halocyanin with a subunit of a terminal oxidase. For further details see Suppl.Text S1 Section 1.
(h) Terminal oxidases are highly diverse in haloarchaea and we restrict our analysis to three species (*Nmn. pharaonis*, *Hfx. volcanii*, and *Hbt. salinarum*) because in each of these at least one terminal oxidase has been experimentally studied (Table 1). Details are described in Suppl. Text S1 Section 1.
(i) NAD-dependent oxidative decarboxylation is a canonical reaction to convert pyruvate into acetyl-CoA, and alpha-ketoglutarate into succinyl-CoA. In haloarchaea, the conversion of pyruvate to acetyl-CoA and alpha-ketoglutarate to succinyl-CoA is dependent on ferredoxin, not on NAD (see above). Nevertheless, most haloarchaeal genomes also code for homologs of enzymes catalyzing NAD-dependent oxidative decarboxylation, such as the *E. coli* pyruvate dehydrogenase complex. In most cases, the substrates could not be identified, an exception being a paralog involved in isoleucine catabolism (Sisignano et al., 2010). In several cases the enzymes were found not to show catalytic activity with pyruvate or alpha-ketoglutarate (see Suppl.Text S1 Section 1 for details). Also, a conditional lethal *porAB* mutant was unable to grow on glucose or pyruvate, thus excluding that alternative enzymes for conversion of pyruvate to acetyl-CoA exist in *Hfx. volcanii* (Kuprat et al., 2021). Nonetheless, despite experimental results to the contrary, pyruvate is assigned as substrate for some of the homologs of the pyruvate dehydrogenase complex in KEGG (as of April 2021).

**Table 1:**
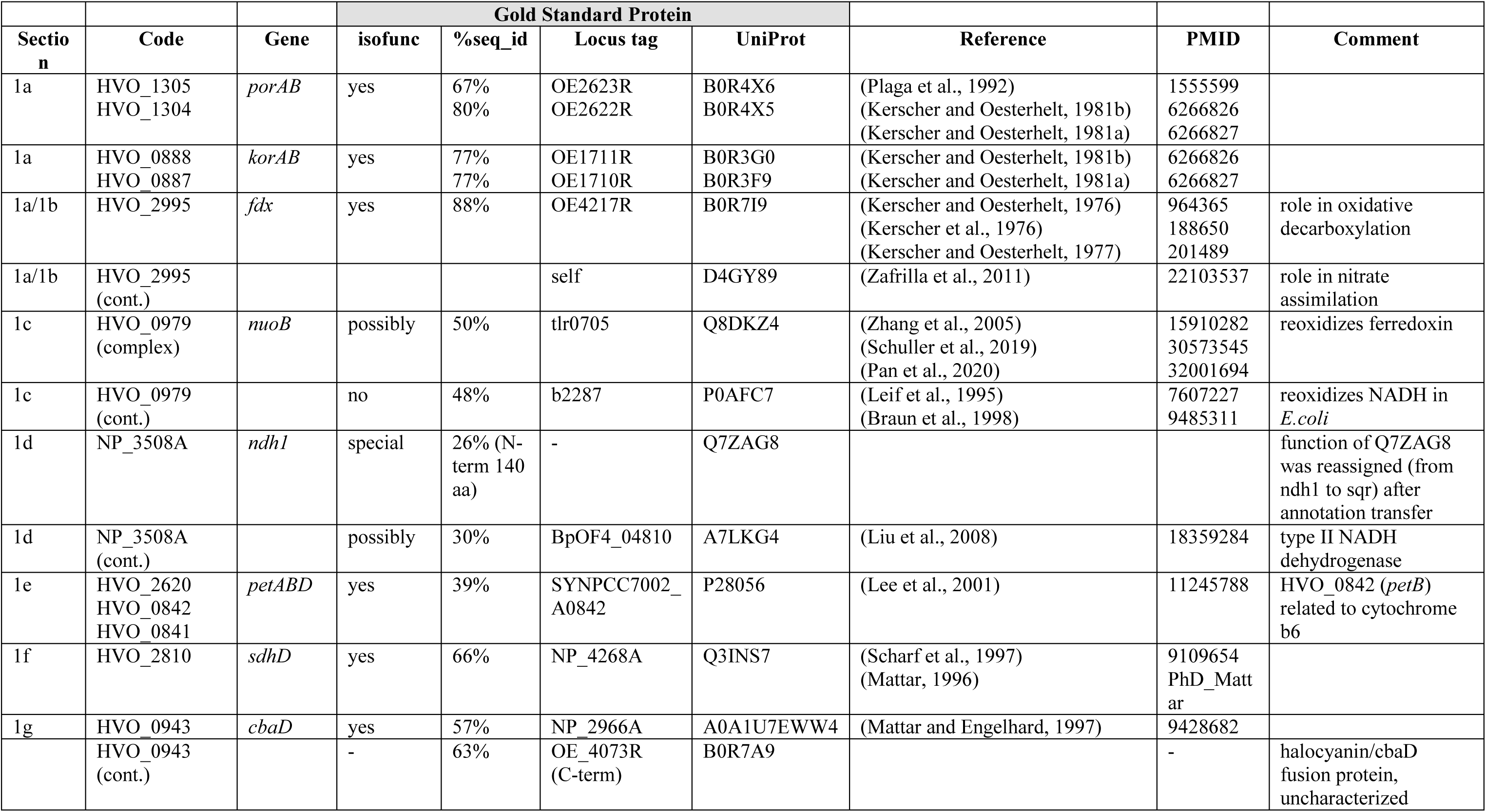
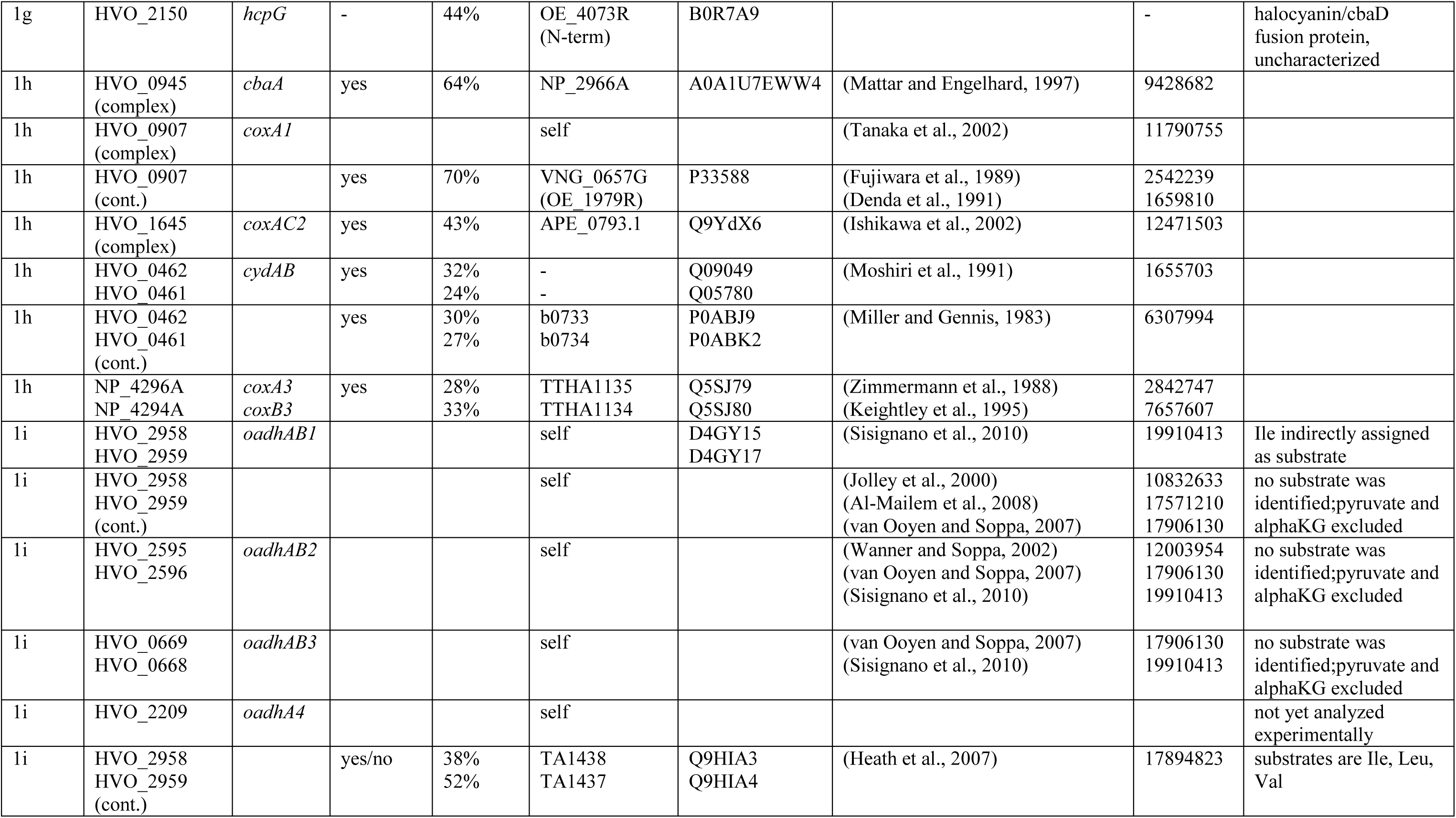
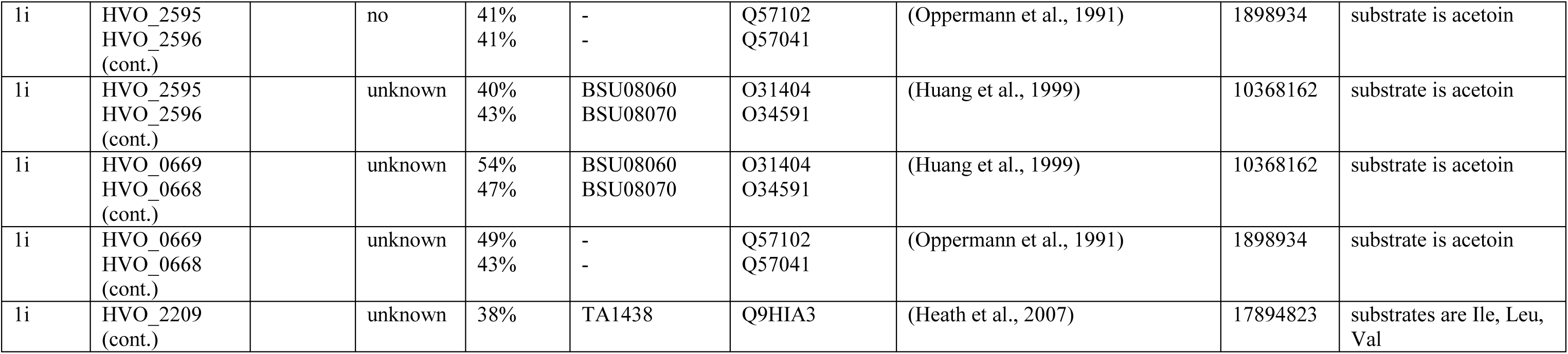
Proteins with open annotation issues and their Gold Standard Protein homologs (Section 1). The <u>column Section</u> refers to the Table listing the protein and to the section in the Results and in Suppl. Text S1. As an example, <u>2c</u> covers topic (<u>c</u>) from the decimal-numbered Results subsection 3.2 Amino Acid Biosynthesis. In Suppl. Text S1, this is covered under Section S<u>2</u> subsection S<u>2</u>.<u>c</u>. The corresponding proteins are listed in Table 2. For a few proteins, two sections are indicated (e.g. 1a/1b). The <u>column Code</u> refers to a haloarchaeal protein by its locus tag, which is mainly from *Haloferax volcanii* (HVO), but also from *Halobacterium salinarum* (OE), *Natronomonas pharaonis* (NP) and *Halohasta litchfieldiae* (halTADL). When the reconstruction of a complete pathway is presented, the unassigned genes are indicated as a “pathway gap”. In one case we indicate the absence of a haloarchaeal ortholog by a dash. In the case of a complex, we either list more than one code, or we list only one subunit together with the term (complex). All subunits of these complexes are listed groupwise in Table S10. A protein may be shown in more than one row. From the 2^nd^ row onwards, this is indicated by the term (cont.). The <u>column Gene</u> lists the assigned gene or a dash if no gene has been assigned. The assigned gene is only indicated in the first row of a protein. A set of four columns is used to relate a query protein to an experimentally characterized homolog, a GSP (Gold Standard Protein) (isofunc, %seq_id, Locus tag, UniProt). The <u>column isofunc</u> indicates if the query protein and its Gold Standard Protein homolog are <u>isofunc</u>tional. The meaning of the terms used in this column in Tables 1-9 (yes, no, yes/no, probably, possibly, unclear, unknown, prediction, special, “-“) is described at the end of this legend. The <u>column %seq_id</u> indicates the protein sequence identity between the query protein and the homologous GSP. The column <u>Locus tag</u> contains the locus tag, if assigned. The <u>column UniProt</u> contains the UniProt accession of the GSP. GSPs have been experimentally characterized as described in a publication. The <u>column Reference</u> links to the reference list of the manuscript. The column PMID lists the PubMed ID of the publication, if available. Otherwise, this is indicated as “not in PubMed”. Also, one PhD thesis is indicated (PhD_Matter). The <u>column Comment</u> provides various types of additional information. The terms used in the column isofunc in Tables 1-9 have the following meaning: The <u>term “yes”</u> indicates that we consider the two proteins as isofunctional and annotate the query protein accordingly. The <u>term “no”</u> is used when we conclude that the proteins differ in function. Additional terms are used for more difficult cases. The <u>term “yes/no”</u> is used for GSPs which are multifunctional, and we assign only one a subset of these functions to the query protein. The <u>term “probably”</u> is used when we consider isofunctionality likely and annotated the query protein accordingly (with the term probable added to the protein name). The <u>term “possibly”</u> is used when we see a good chance that the proteins are isofunctional, but consider it too speculative to annotate the protein accordingly. The <u>term “unclear”</u> is used when we consider it likely that the same overall reaction is catalyzed, but when reaction details, e.g. the energy-providing compound, is unresolved. The <u>term “unknown”</u> is used when it is not possible to predict the substrate of the query protein. The <u>term “prediction”</u> is used if a function assignment is based on bioinformatic analyses but not yet on an experimentally characterized homologous protein. The <u>term “special”</u> is used when multiple arguments have to be considered with full details provided in the corresponding section of Suppl. Text S1. Finally, <u>a dash (“-“)</u> is used when isofunctionality does not apply, e.g. when a homologous Gold Standard Protein could not be identified.

### 3.2 Amino acid metabolism

While most amino acid biosynthesis and degradation pathways can be reliably reconstructed, a few open issues remain, which are discussed below.

(a) The first and last steps of arginine biosynthesis deal with blocking and unblocking of the alpha-amino group of the substrate (glutamate) and a product intermediate (ornithine). As detailed in Suppl. Text S1 Section 2, it is highly likely that glutamate is attached to the gamma-carboxyl group of a carrier protein, and ornithine is released from that carrier protein. This is based on characterized proteins from *Thermus thermophilus* (Horie et al., 2009), *Thermococcus kodakarensis* (Yoshida et al., 2016) and *Sulfolobus acidocaldarius* (Ouchi et al., 2013). The assignment is strongly supported by clustering of the arginine biosynthesis genes. Some of the homologs are bifunctional, being involved in arginine biosynthesis but also in lysine biosynthesis via the prokaryotic variant of the alpha-aminoadipate pathway. This ambiguity is not assumed to occur in haloarchaea, which use the diaminopimelate pathway for Lys biosynthesis (Hochuli et al., 1999) (see Suppl. Text S1 Section 2 for further discussion of this issue).

Expanding the above, we provide full details underlying our reconstruction of arginine and lysine biosynthesis in *Hfx, volcanii* in Table 2.

(b) Archaea use a different precursor for aromatic amino acid biosynthesis than the classical pathway. This has been resolved for *Methanocaldococcus jannaschii* and for *Methanococcus maripaludis* (White, 2004, Porat et al., 2006). However, the initial steps may differ from those reported for *Methanocaldococcus* in that fructose-1,6-disphosphate rather than 6-deoxy-5-ketofructose might be a substrate (Gulko et al., 2014). Up to now, a clean deletion of the corresponding enzymes and confirmation with in vitro assays has not yet been achieved (for details see Suppl. Text S1 Section 2).
(c) The gene for tryptophanase (*tpa*) is stringently regulated in *Haloferax*, which is the basis for using its promoter in the toolbox for regulated gene expression (Large et al., 2007). The shutdown of this gene avoids tryptophan degradation when supplies are scarce. Tryptophanase cleaves tryptophan into indole, pyruvate and ammonia. The fate of indole is, however, yet unresolved.
(d) A probable histidine utilization cluster exists, based on characterized homologs from *Bacillus subtilis*, but has not yet been experimentally verified.
(e) Among 16 auxotrophic mutants observed in a *Hfx. volcanii* transposon insertion library (Kiljunen et al., 2014), some could grow only in the presence of one (or several) supplied amino acids. In many cases, the affected genes were known to be involved in the corresponding pathway, but the others may lead to novel function assignments. One affected gene resulted in histidine auxotrophy and the product of this gene (HVO_0431) is an interesting candidate. The InterPro domain assignment (HAD family hydrolase) fits to the only remaining pathway gap in histidine biosynthesis (histidinol-phosphatase). In this context it should be noted that the enzyme which catalyzes the preceding reaction (encoded by *hisC*) is part of a highly conserved three-gene operon involved in polar lipid biosynthesis (see below). For details see Suppl. Text S1 Section 2. One affected gene resulted in isoleucine auxotrophy. The product of this gene (HVO_0644) is currently annotated to catalyze two reactions, one being an early step in isoleucine biosynthesis (EC 2.3.1.182), the other being the first step after leucine biosynthesis branches off from valine biosynthesis (EC 2.3.3.13) (see below, f) (for details see Suppl. Text S1 Section 2).
(f) *Hfx. volcanii* codes for two paralogs with an attributed function as 2-isopropylmalate synthase (EC 2.3.3.13). This is the first reaction specific to leucine biosynthesis, when the pathway branches off valine biosynthesis. One paralog, HVO_0644, is annotated as bifunctional, also catalyzing a chemically similar reaction which is an early step in isoleucine biosynthesis (EC 2.3.1.182). When the gene encoding HVO_0644 is disrupted by transposon integration, cells cannot grow in the absence of isoleucine. It is unclear if the protein is really bifunctional and is really involved in leucine biosynthesis, catalyzing the reaction of EC 2.3.3.13. The other paralog, HVO_1510, belongs to an ortholog set with major problems concerning start codon assignment. The ortholog set from the 16 genomes listed in Suppl. Table S11 were analyzed. When only canonical start codons are considered (ATG, GTG, TTG), then the orthologs from *Haloferax mediterranei*, *Nmn. pharaonis*, *Natronomonas moolapensis* and *Halohasta litchfieldiae* either lack a long highly conserved N-terminal region, or they are disrupted (pseudogenes), being devoid of a potential start codon. The gene from *Hfx. volcanii* has a start codon (GTG) which is consistent to that of *Haloferax gibbonsii* strain LR2-5 (but a GTA in *Hfx. gibbonsii* strain ARA6). In this region, the gene from *Hfx. mediterranei* is closely related but has in-frame stop codons. HVO_1510 is considerably longer than the orthologs from *Haloquadratum walsbyi*, *Haloarcula hispanica*, and *Natrialba magadii*. The first alternative start codon for HVO_1510 codes for Met-93. This protein was proteomically identified in three ArcPP datasets (Schulze et al., 2020), and peptides upstream of Met-93 were identified. This gene might be translated from an atypical start codon, either an in-frame CTG, or an out-of-frame ATG, which would require ribosomal slippage (for details see Suppl. Text S1 Section 2). It is tempting to speculate that translation occurs only when leucine is not available.

**Table 2:** Proteins with open annotation issues and their Gold Standard Protein homologs (Section 2). For a description of this table see the legend to Table 1.

| Section | Code | Gene | Gold Standard Protein |  |  |  | Reference | PMID | Comment |
| --- | --- | --- | --- | --- | --- | --- | --- | --- | --- |
|  |  |  | isofunc | %seq_id | Locus tag | UniProt |  |  |  |
| 2a | HVO_0047 | *argW* | no | 54% | TT_C1544 | Q72HE5 | (Yoshida et al., 2015) | 25392000 | for Arg, not for Lys biosynthesis |
| 2a | HVO_0047 (cont.) |  | yes/no | 39% | Saci_0753 | Q4J AQ0 |  |  | only for Arg, not for Lys biosynthesis |
| 2a | HVO_0047 (cont.) |  | yes/no | 61% | TK0279 | Q5JFV9 | (Yoshida et al., 2016) | 27566549 | only for Arg, not for Lys biosynthesis |
| 2a | HVO_0046 | *argX* | no | 44% | TT_C1543 | Q72HE6 | (Horie et al., 2009) | 19620981 | for Arg, not for Lys biosynthesis |
| 2a | HVO_0046 (cont.) |  | yes | 30% | Saci_1621 | Q4J8E7 |  |  | only for Arg, not for Lys biosynthesis |
| 2a | HVO_0046 (cont.) |  | yes/no | 37% | TK0278 | Q5JFW0 | (Yoshida et al., 2016) | 27566549 | only for Arg, not for Lys biosynthesis |
| 2a | HVO_0044 | *argB* | no | 41% | TT_C1541 | O50147 | (Horie et al., 2009)<br>(Yoshida et al., 2015) | 19620981<br>25392000 | for Arg, not for Lys biosynthesis |
| 2a | HVO_0044 (cont.) |  | yes/no | 33% | Saci_0751 | Q4J AQ2 | (Ouchi et al., 2013) | 23434852 | only for Arg, not for Lys biosynthesis |
| 2a | HVO_0044 (cont.) |  | yes/no | 32% | TK0276 | Q5JFW2 | (Yoshida et al., 2016) | 27566549 | only for Arg, not for Lys biosynthesis |
| 2a | HVO_0045 | *argC* | no | 48% | TT_C1542 | O50146 | (Horie et al., 2009)<br>(Shimizu et al., 2016) | 19620981<br>26966182 | for Arg, not for Lys biosynthesis |
| 2a | HVO_0045 (cont.) |  | yes/no | 42% | Saci_0750 | Q4J AQ3 | (Ouchi et al., 2013) | 23434852 | only for Arg, not for Lys biosynthesis |
| 2a | HVO_0045 (cont.) |  | yes/no | 46% | TK0277 | Q5JFW1 | (Yoshida et al., 2016) | 27566549 | only for Arg, not for Lys biosynthesis |
| 2a | HVO_0043 | *argD* | no | 45% | TT_C1393 | Q93R93 | (Miyazaki et al., 2001) | 11489859 | for Arg, not for Lys biosynthesis |
| 2a | HVO_0043 (cont.) |  | yes/no | 40% | Saci_0755 | Q4J AP8 | (Ouchi et al., 2013) | 23434852 | only for Arg, not for Lys biosynthesis |
| 2a | HVO_0043 (cont.) |  | yes/no | 42% | TK0275 | Q5JFW3 | (Yoshida et al., 2016) | 27566549 | only for Arg, not for Lys biosynthesis |
| 2a | HVO_0042 | *argE* | no | 36% | TT_C1396 | Q8VUS5 | (Horie et al., 2009)<br>(Fujita et al., 2017) | 19620981<br>28720495 | for Arg, not for Lys biosynthesis |
| 2a | HVO_0042<br>(cont.) |  | yes/no | 29% | Saci_0756 | Q4JAP7 | (Ouchi et al., 2013) | 23434852 | only for Arg, not for Lys biosynthesis |
| 2a | HVO_0042<br>(cont.) |  | yes/no | 37% | TK0274 | Q5JFW4 | (Yoshida et al., 2016) | 27566549 | only for Arg, not for Lys biosynthesis |
| 2a | HVO_0041 | *argF* | yes | 50% | P18186 | BSU11250 | (Issaly and Issaly, 1974) | 4216455 |  |
| 2a | HVO_0041<br>(cont.) |  | yes | 47% | OE_5205R | B0R9X3 | (Ruepp et al., 1995) | 7868583 |  |
| 2a | HVO_0049 | *argG* | yes | 35% | - | P00966 | (Shaheen et al., 1994) | 8792870 | human |
| 2a | HVO_0049<br>(cont.) |  | yes | 23% | b3172 | P0A6E4 | (Lemke et al., 1999) | 10666579 | *E. coli* |
| 2a | HVO_0048 | *argH* | yes | 38% | MMP0013 | O74026 | (Cohen-Kupiec et al., 1999) | 10220900 |  |
| 2a | HVO_0008 | *lysC* | yes | 32% | BSU28470 | P08495 | (Kato et al., 2004) | 15033471 |  |
| 2a | HVO_2487 | *asd* | yes | 51% | MJ0205 | Q57658 | (Faehnle et al., 2005) | 16225889 |  |
| 2a/9e | HVO_1101 | *dapA* | yes | 45% | PA1010 | Q9I4W3 | (Kaur et al., 2011) | 21396954 |  |
| 2a | HVO_1100 | *dapB* | yes | 33% | b0031 | P04036 | (Reddy et al., 1995) | 7893644 |  |
| 2a | HVO_1099 | *dapD* | yes | 32% | b0166 | P0A9D8 | (Simms et al., 1984) | 6365916 |  |
| 2a | HVO_1096 | *dapE* | yes | 29% | b2472 | P0AED7 | (Lin et al., 1988) | 3276674 | function supported by gene clustering |
| 2a | HVO_1097 | *dapF* | yes | 35% | b3809 | P0A6K1 | (Wiseman and Nichols, 1984) | 6378903 |  |
| 2a | HVO_1098 | *lysA* | yes | 38% | b2838 | P00861 | (White and Kelly, 1965) | 14343156 |  |
| 2a | HVO_A0634 | - | unknown | 25% | b2472 | P0AED7 | (Lin et al., 1988) | 3276674 | function assigned to HVO_1096 in *dap* cluster |
| 2b | HVO_0790 | *fba2* | special | 67% | OE_1472F | B0R334 | (Gulko et al., 2014) | 25216252 | EC 2.2.1.10 activity of OE_1472F not yet confirmed in vitro |
| 2b | HVO_0790<br>(cont.) |  | special | 45% | MJ0400 | Q57843 | (White, 2004) | 15182204 | substrate uncertain |
| 2b | HVO_0792 | *aroB* | yes | 69% | OE_1475F | B0R336 | (Gulko et al., 2014) | 25216252 | OE_1475F only partially characterized |
| 2b | HVO_0792<br>(cont.) |  | yes | 44% | MJ1249 | Q58646 | (White, 2004) | 15182204 |  |
| 2b | HVO_0602 | *aroD1* | yes | 44% | OE_1477R | B0R338 | (Gulko et al., 2014) | 25216252 |  |
| 2b | HVO_0602<br>(cont.) |  | yes | 31% | MMP1394 | Q6LXF7 | (Porat et al., 2004) | 15262931 |  |
| 2c | HVO_0009 | *tnaA* | yes | 41% | b3708 | P0A853 | (Phillips and Gollnick, 1989)<br>(Newton et al., 1965) | 2659590<br>14284727 |  |
| 2d | HVO_A0559 | *hutH* | yes | 42% | BSU39350 | P10944 | (Oda et al., 1988)<br>(Hartwell and Magasanik, 1963) | 2454913<br>14066617 |  |
| 2d | HVO_A0562 | *hutU* | yes | 62% | BSU39360 | P25503 | (Kaminskas et al., 1970) | 4990470 |  |
| 2d | HVO_A0560 | *hutI* | yes | 42% | BSU39370 | P42084 | (Yu et al., 2006) | 16990261 |  |
| 2d | HVO_A0561 | *hutG* | yes | 33% | BSU39380 | P42068 | (Kaminskas et al., 1970) | 4990470 |  |
| 2e | HVO_0431 | - | - |  |  |  |  |  | no GSP available |
| 2e | HVO_0644 | *leuA1* | yes/no | 47% | MJ1392 | Q58787 | (Howell et al., 1999) | 9864346 | HVO_0644 monofunc<br>(CimA) or bifunc<br>(CimA+LeuA);<br>MJ1392 CimA |
| 2e | HVO_0644<br>(cont.) |  | unclear | 44% | MJ1195 | Q58595 | (Howell et al., 1998) | 9665716 | HVO_0644 monofunc<br>(CimA) or bifunc<br>(CimA+LeuA);<br>MJ1195 LeuA |
| 2e/2f | HVO_1510 | *leuA2* | yes | 47% | MJ1195 | Q58595 | (Howell et al., 1998) | 9665716 | HVO_1510 LeuA;<br>MJ1195 LeuA |
| 2e/2f | HVO_1510<br>(cont.) |  | no | 41% | MJ1392 | Q58787 | (Howell et al., 1999) | 9864346 | HVO_1510 LeuA<br>MJ1392 CimA |
| 2e | HVO_A0489 | - | no | 31% | MJ1392 | Q58787 | (Howell et al., 1999) | 9864346 | HVO_A0489 general<br>function only;<br>MJ1392 CimA |
| 2e | HVO_A0489<br>(cont.) |  | no | 30% | MJ1195 | Q58595 | (Howell et al., 1998) | 9665716 | HVO_A0489 general<br>function only;<br>MJ1195 LeuA |
| 2e | HVO_1153 | - | - |  |  |  |  |  | function unassigned;<br>no GSP |

### 3.3 Coenzymes I: cobalamin and heme

The classical heme biosynthesis pathway branches off cobalamin biosynthesis at the level of uroporphyrinogen III. The alternative heme biosynthesis pathway (Bali et al., 2011), which is used by haloarchaea, has an additional common step, the conversion of uroporphyrinogen III to precorrin-2. For heme biosynthesis, precorrin-2 is converted to siroheme. This pathway has been reconstructed (Siddaramappa et al., 2012), except for the iron insertion step. For de novo cobalamin biosynthesis, haloarchaea use the cobalt-early pathway with a cobalt-dependent key reaction being catalyzed by CbiG (Moore et al., 2013). Several aspects of heme and cobalamin biosynthesis in haloarchaea are yet unresolved. This is illustrated in Figure 1.

(a) *Hfx. volcanii* contains two annotated *cbiX* genes. For reasons detailed in Suppl.Text S1 Section 3, we predict that one is a cobaltochelatase, involved in cobalamin biosynthesis, while the other is a ferrochelatase, responsible for conversion of precorrin-2 to siroheme in the alternative heme biosynthesis pathway.
(b) De novo cobalamin biosynthesis has been extensively reconstructed upon curation of the genome annotation (Pfeiffer and Oesterhelt, 2015). All enzymes of the pathway and their associated GSPs are listed in Table 3. Only two pathway gaps remained, and because these are consecutive, it may be possible that the haloarchaeal pathway is non-canonical and proceeds via a novel biosynthetic intermediate. There are only four genes with yet unassigned function in the *Hfx. volcanii* cobalamin gene cluster, and their synteny is well conserved in the majority of haloarchaeal genomes. Thus, these genes are obvious candidates for filling the pathway gaps (for details see Suppl.Text S1 Section 3).
(c) The cobalamin biosynthesis and salvage reactions (those beyond ligand cobyrinate a,c diamide) involve “adenosylation of the corrin ring, attachment of the aminopropanol arm, and assembly of the nucleotide loop that bridges the lower ligand dimethylbenzimidazole and the corrin ring” (Rodionov et al., 2003). The enzymes of these branches of cobalamin biosynthesis and their associated GSPs are listed in Table Only two pathway gaps remain open. For one of these, a candidate was proposed upon detailed bioinformatic analysis (Rodionov et al., 2003) (for further details see Suppl.Text S1 Section 3).
(d) Haloarchaea may code for a late cobaltochelatase of the heterotrimeric type. Distantly related GSPs are either cobalt or magnesium chelatases. A late cobaltochelatase is not required for de novo cobalamin biosynthesis via the cobalt-early pathway. We speculate that it may be involved in cobalamin salvage. The chelatase has a mosaic subunit structure as also reported previously (Rodionov et al., 2003) (see Suppl.Text S1 Section 3 for details).
(e) In the alternative heme biosynthesis pathway, siroheme is decarboxylated to 12,18-didecarboxysiroheme, which is attributed to the proteins encoded by *ahbA* and *ahbB*. These are homologous to each other and are organized as two two-domain proteins. It is unclear if AhbA and AhbB function independently or if they form a complex.

**Figure 1.**
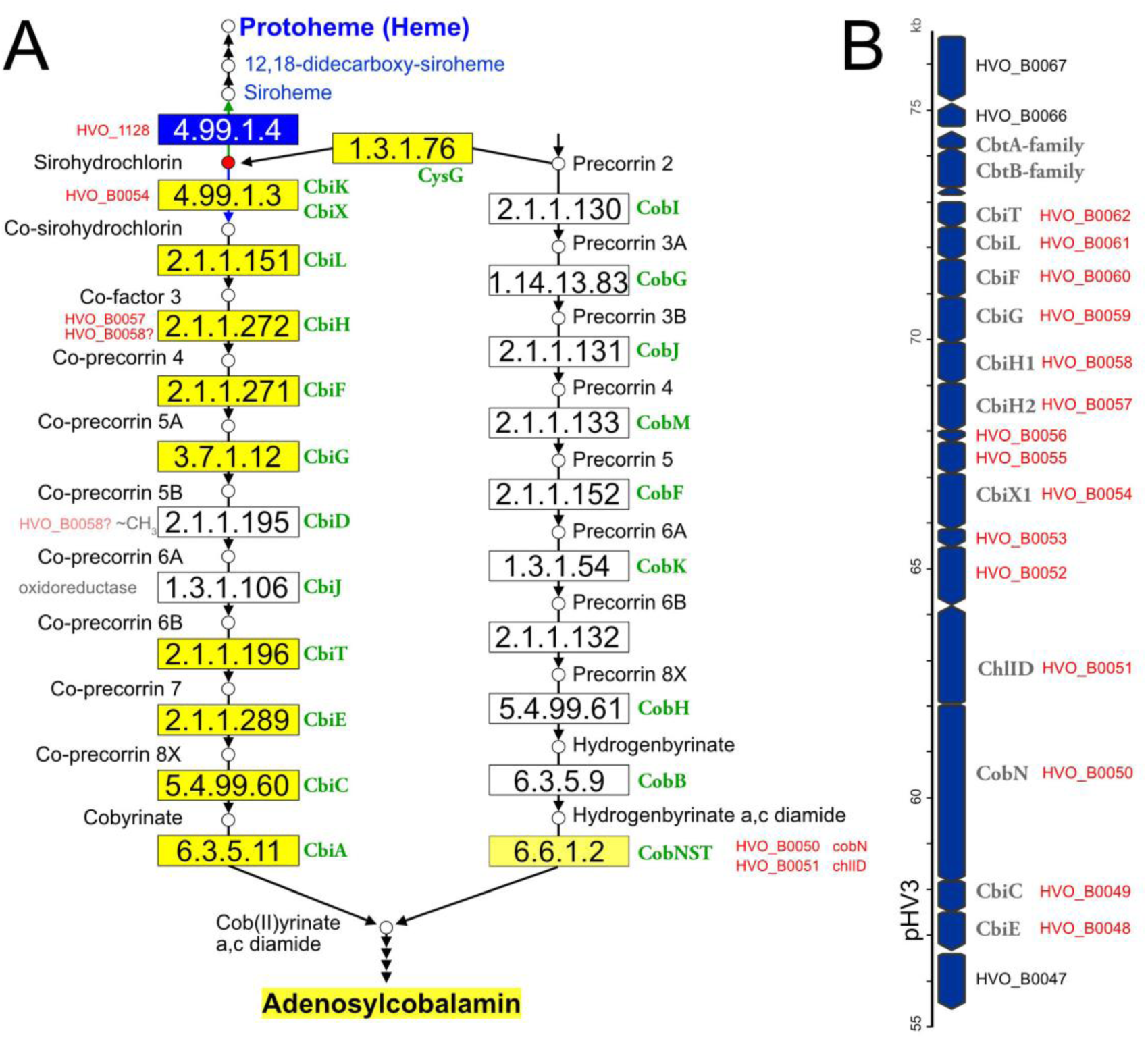
Illustration of the haloarchaeal cobalamin and heme biosynthesis pathways and of the major cobalamin biosynthesis gene cluster. (A) Biosynthesis pathways. This illustration is based on the corresponding KEGG map 00860. Small circles represent pathway intermediates and have their names assigned. Pathway intermediates upstream of Precorrin 2 are not displayed. The circle for sirohydrochlorin is highlighted in red as this is the branchpoint for heme and cobalamin biosynthesis in haloarchaea. Enzymatic reactions are shown by arrows, EC numbers being provided in rectangular boxes. Rectangles are colored when the enzyme has been reconstructed for haloarchaea (blue: heme biosynthesis; dark yellow: de novo cobalamin biosynthesis; light yellow: late cobaltochelatase which may be a salvage reaction). Gene names in green are adopted from KEGG and represent those from bacterial model pathways. Consecutive arrowheads indicate reaction series which are not shown in detail for space reasons. For enzymatic reactions which are considered to be open issues, the *Hfx. volcanii* locus tags are provided. For two pathway gaps (white boxes in the cobalt-early pathway), the type of reaction is indicated (oxidoreductase and ∼CH3, indicating a methylation reaction). The question mark after HVO_B0058 indicates that this protein, currently co-attributed to EC 2.1.1.272, is a candidate for the yet unassigned EC 2.1.1.195 reaction. We note that haloarchaea might use a deviating biosynthesis pathway, e.g. by swapping the methylation and oxidoreductase reactions (not illustrated). (B) The major cobalamin cluster, encoded on megaplasmid pHV3. Arrows are used to indicate the coding strand and are roughly drawn to scale. If assigned, the gene name is provided in addition to the *Hfx. volcanii* locus tag. Locus tags in red indicate genes that are part of the cobalamin cluster.

**Table 3:**
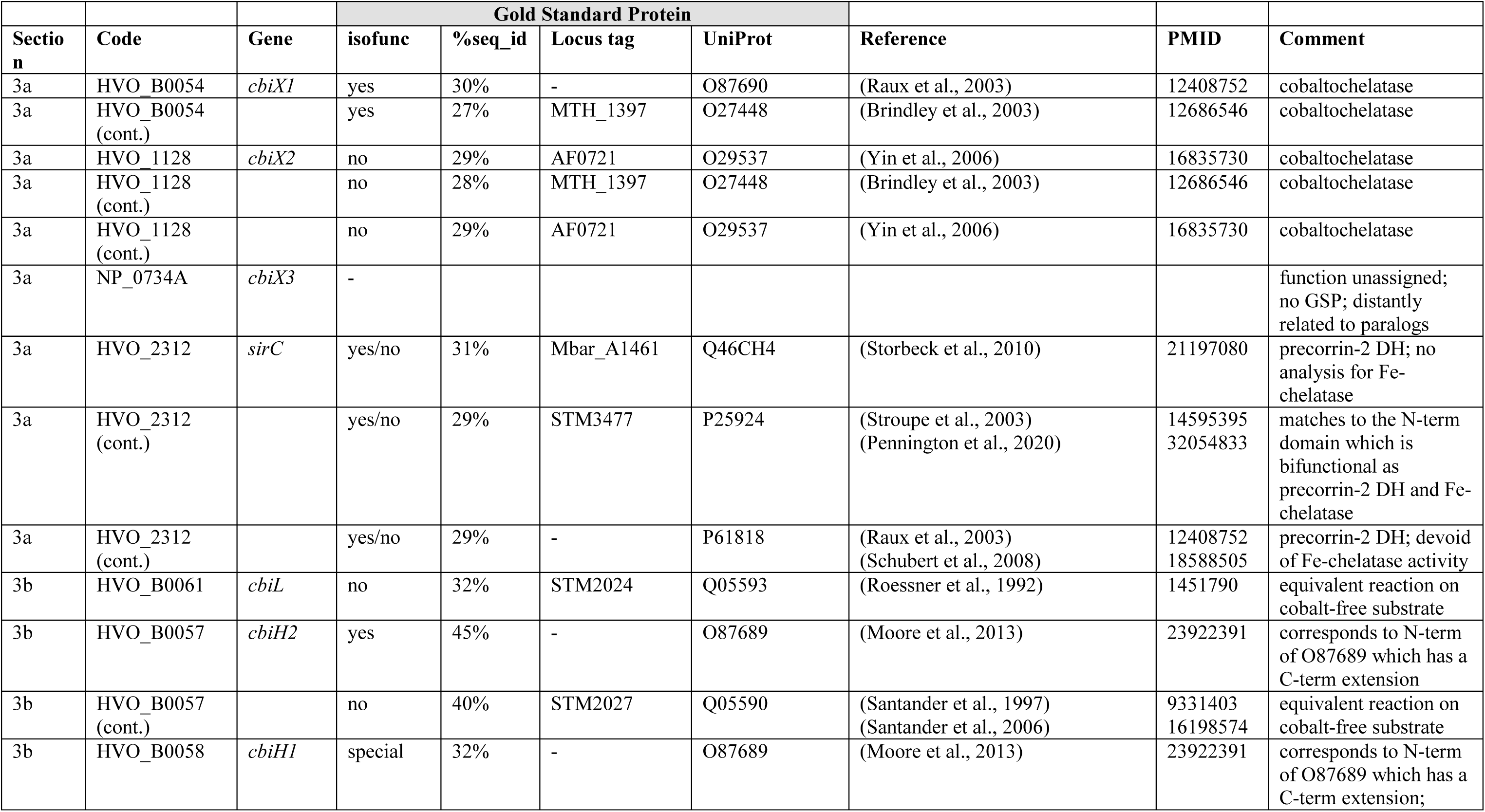
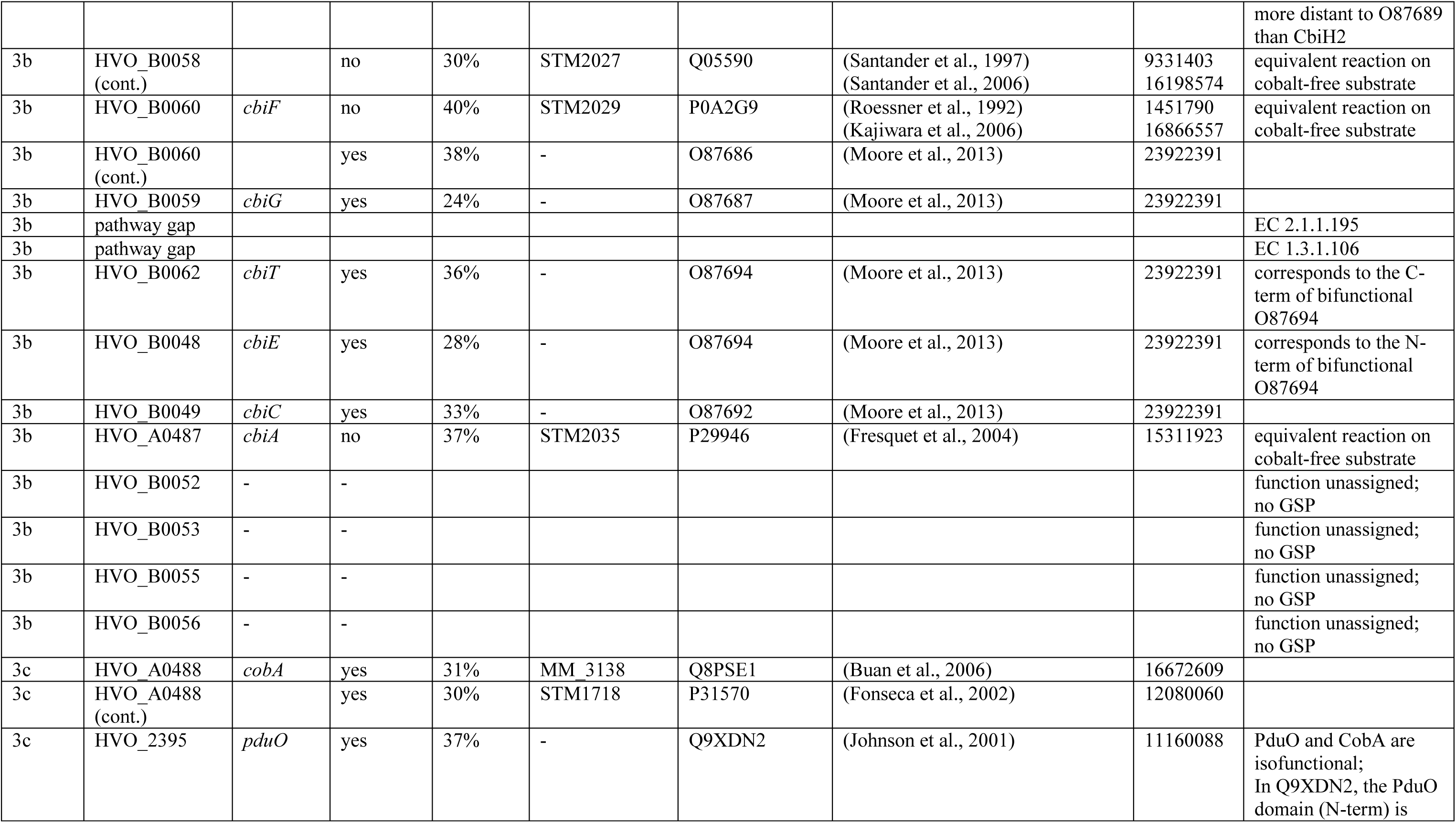
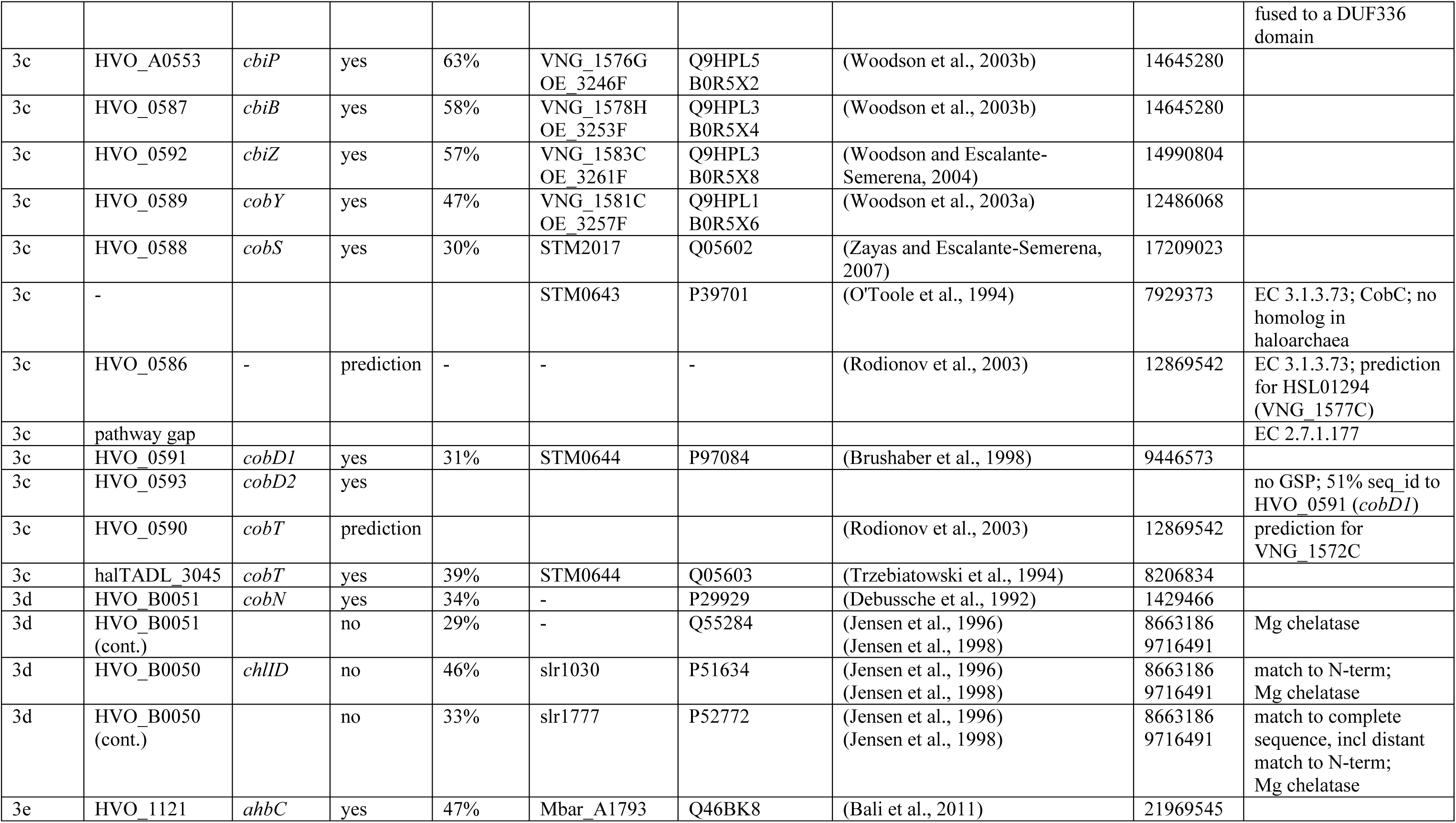
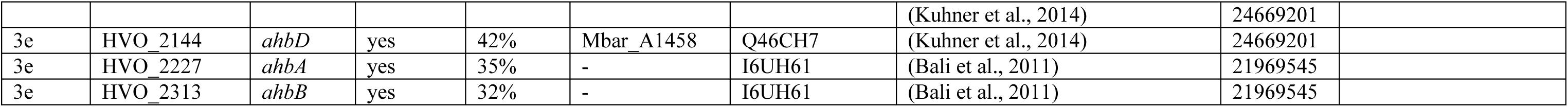
Proteins with open annotation issues and their Gold Standard Protein homologs (Section 3). For a description of this table see the legend to Table 1.

| Section | Code | Gene | Gold Standard Protein |  |  |  | Reference | PMID | Comment |
| --- | --- | --- | --- | --- | --- | --- | --- | --- | --- |
|  |  |  | isofunc | %seq_id | Locus tag | UniProt |  |  |  |
| 3a | HVO_B0054 | *cbiX1* | yes | 30% | - | O87690 | (Raux et al., 2003) | 12408752 | cobaltochelatase |
| 3a | HVO_B0054 (cont.) |  | yes | 27% | MTH_1397 | O27448 | (Brindley et al., 2003) | 12686546 | cobaltochelatase |
| 3a | HVO_1128 | *cbiX2* | no | 29% | AF0721 | O29537 | (Yin et al., 2006) | 16835730 | cobaltochelatase |
| 3a | HVO_1128 (cont.) |  | no | 28% | MTH_1397 | O27448 | (Brindley et al., 2003) | 12686546 | cobaltochelatase |
| 3a | HVO_1128 (cont.) |  | no | 29% | AF0721 | O29537 | (Yin et al., 2006) | 16835730 | cobaltochelatase |
| 3a | NP_0734A | *cbiX3* | - |  |  |  |  |  | function unassigned; no GSP; distantly related to paralogs |
| 3a | HVO_2312 | *sirC* | yes/no | 31% | Mbar_A1461 | Q46CH4 | (Storbeck et al., 2010) | 21197080 | precorrin-2 DH; no analysis for Fe-chelatase |
| 3a | HVO_2312 (cont.) |  | yes/no | 29% | STM3477 | P25924 | (Stroupe et al., 2003)<br>(Pennington et al., 2020) | 14595395<br>32054833 | matches to the N-term domain which is bifunctional as precorrin-2 DH and Fe-chelatase |
| 3a | HVO_2312 (cont.) |  | yes/no | 29% | - | P61818 | (Raux et al., 2003)<br>(Schubert et al., 2008) | 12408752<br>18588505 | precorrin-2 DH; devoid of Fe-chelatase activity |
| 3b | HVO_B0061 | *cbiL* | no | 32% | STM2024 | Q05593 | (Roessner et al., 1992) | 1451790 | equivalent reaction on cobalt-free substrate |
| 3b | HVO_B0057 | *cbiH2* | yes | 45% | - | O87689 | (Moore et al., 2013) | 23922391 | corresponds to N-term of O87689 which has a C-term extension |
| 3b | HVO_B0057 (cont.) |  | no | 40% | STM2027 | Q05590 | (Santander et al., 1997)<br>(Santander et al., 2006) | 9331403<br>16198574 | equivalent reaction on cobalt-free substrate |
| 3b | HVO_B0058 | *cbiH1* | special | 32% | - | O87689 | (Moore et al., 2013) | 23922391 | corresponds to N-term of O87689 which has a C-term extension; |
|  |  |  |  |  |  |  |  |  | more distant to O87689 than CbiH2 |
| 3b | HVO_B0058 (cont.) |  | no | 30% | STM2027 | Q05590 | (Santander et al., 1997)<br>(Santander et al., 2006) | 9331403<br>16198574 | equivalent reaction on cobalt-free substrate |
| 3b | HVO_B0060 | *cbiF* | no | 40% | STM2029 | P0A2G9 | (Roessner et al., 1992)<br>(Kajiwara et al., 2006) | 1451790<br>16866557 | equivalent reaction on cobalt-free substrate |
| 3b | HVO_B0060 (cont.) |  | yes | 38% | - | O87686 | (Moore et al., 2013) | 23922391 |  |
| 3b | HVO_B0059 | *cbiG* | yes | 24% | - | O87687 | (Moore et al., 2013) | 23922391 |  |
| 3b | pathway gap |  |  |  |  |  |  |  | EC 2.1.1.195 |
| 3b | pathway gap |  |  |  |  |  |  |  | EC 1.3.1.106 |
| 3b | HVO_B0062 | *cbiT* | yes | 36% | - | O87694 | (Moore et al., 2013) | 23922391 | corresponds to the C-term of bifunctional O87694 |
| 3b | HVO_B0048 | *cbiE* | yes | 28% | - | O87694 | (Moore et al., 2013) | 23922391 | corresponds to the N-term of bifunctional O87694 |
| 3b | HVO_B0049 | *cbiC* | yes | 33% | - | O87692 | (Moore et al., 2013) | 23922391 |  |
| 3b | HVO_A0487 | *cbiA* | no | 37% | STM2035 | P29946 | (Fresquet et al., 2004) | 15311923 | equivalent reaction on cobalt-free substrate |
| 3b | HVO_B0052 | - | - |  |  |  |  |  | function unassigned; no GSP |
| 3b | HVO_B0053 | - | - |  |  |  |  |  | function unassigned; no GSP |
| 3b | HVO_B0055 | - | - |  |  |  |  |  | function unassigned; no GSP |
| 3b | HVO_B0056 | - | - |  |  |  |  |  | function unassigned; no GSP |
| 3c | HVO_A0488 | *cobA* | yes | 31% | MM_3138 | Q8PSE1 | (Buan et al., 2006) | 16672609 |  |
| 3c | HVO_A0488 (cont.) |  | yes | 30% | STM1718 | P31570 | (Fonseca et al., 2002) | 12080060 |  |
| 3c | HVO_2395 | *pduO* | yes | 37% | - | Q9XDN2 | (Johnson et al., 2001) | 11160088 | PduO and CobA are isofunctional; In Q9XDN2, the PduO domain (N-term) is fused to a DUF336 domain |
| 3c | HVO_A0553 | *cbiP* | yes | 63% | VNG_1576G OE_3246F | Q9HPL5 B0R5X2 | (Woodson et al., 2003b) | 14645280 |  |
| 3c | HVO_0587 | *cbiB* | yes | 58% | VNG_1578H OE_3253F | Q9HPL3 B0R5X4 | (Woodson et al., 2003b) | 14645280 |  |
| 3c | HVO_0592 | *cbiZ* | yes | 57% | VNG_1583C OE_3261F | Q9HPL3 B0R5X8 | (Woodson and Escalante-Semerena, 2004) | 14990804 |  |
| 3c | HVO_0589 | *cobY* | yes | 47% | VNG_1581C OE_3257F | Q9HPL1 B0R5X6 | (Woodson et al., 2003a) | 12486068 |  |
| 3c | HVO_0588 | *cobS* | yes | 30% | STM2017 | Q05602 | (Zayas and Escalante-Semerena, 2007) | 17209023 |  |
| 3c | - |  |  |  | STM0643 | P39701 | (O'Toole et al., 1994) | 7929373 | EC 3.1.3.73; CobC; no homolog in haloarchaea |
| 3c | HVO_0586 | - | prediction | - | - | - | (Rodionov et al., 2003) | 12869542 | EC 3.1.3.73; prediction for HSL01294 (VNG_1577C) |
| 3c | pathway gap |  |  |  |  |  |  |  | EC 2.7.1.177 |
| 3c | HVO_0591 | *cobD1* | yes | 31% | STM0644 | P97084 | (Brushaber et al., 1998) | 9446573 |  |
| 3c | HVO_0593 | *cobD2* | yes |  |  |  |  |  | no GSP; 51% seq_id to HVO_0591 (*cobD1*) |
| 3c | HVO_0590 | *cobT* | prediction |  |  |  | (Rodionov et al., 2003) | 12869542 | prediction for VNG_1572C |
| 3c | halTADL_3045 | *cobT* | yes | 39% | STM0644 | Q05603 | (Trzebiatowski et al., 1994) | 8206834 |  |
| 3d | HVO_B0051 | *cobN* | yes | 34% | - | P29929 | (Debussche et al., 1992) | 1429466 |  |
| 3d | HVO_B0051 (cont.) |  | no | 29% | - | Q55284 | (Jensen et al., 1996)<br>(Jensen et al., 1998) | 8663186<br>9716491 | Mg chelatase |
| 3d | HVO_B0050 | *chlID* | no | 46% | slr1030 | P51634 | (Jensen et al., 1996)<br>(Jensen et al., 1998) | 8663186<br>9716491 | match to N-term;<br>Mg chelatase |
| 3d | HVO_B0050 (cont.) |  | no | 33% | slr1777 | P52772 | (Jensen et al., 1996)<br>(Jensen et al., 1998) | 8663186<br>9716491 | match to complete<br>sequence, incl distant<br>match to N-term;<br>Mg chelatase |
| 3e | HVO_1121 | *ahbC* | yes | 47% | Mbar_A1793 | Q46BK8 | (Bali et al., 2011) | 21969545 |  |
|  |  |  |  |  |  |  | (Kuhner et al., 2014) | 24669201 |  |
| 3e | HVO_2144 | *ahbD* | yes | 42% | Mbar_A1458 | Q46CH7 | (Kuhner et al., 2014) | 24669201 |  |
| 3e | HVO_2227 | *ahbA* | yes | 35% | - | I6UH61 | (Bali et al., 2011) | 21969545 |  |
| 3e | HVO_2313 | *ahbB* | yes | 32% | - | I6UH61 | (Bali et al., 2011) | 21969545 |  |

### 3.4 Coenzymes II: coenzyme F420

Even though coenzyme F420 is predominantly associated with methanogenic archaea (Eirich et al., 1979, Jaenchen et al., 1984), it occurs also in bacteria and a small amount of this coenzyme has been detected in non-methanogenic archaea, including halophiles (Lin and White, 1986). The genes required for the biosynthesis of this coenzyme are encoded in haloarchaeal genomes, but the origin and attachment of the phospholactate moiety are not completely resolved (see below). To the best of our knowledge, only a single coenzyme F420 dependent enzymatic reaction has yet been reported for halophilic archaea (de Wit and Eker, 1987). Thus, the importance of this coenzyme in haloarchaeal biology is currently enigmatic and awaits experimental analysis.

(a) The pathway that creates the carbon backbone of this coenzyme has been reconstructed. We list the enzymes with their associated GSPs in Table 4. Coenzyme F420 contains a phospholactate moiety, which was reported to originate from 2-phospho-lactate (Grochowski et al., 2008), but this compound is not well connected to the remainder of metabolism. As summarized in Suppl.Text S1 Section 4, there are various new insights regarding this pathway from recent studies in other prokaryotes (Bashiri et al., 2019, Braga et al., 2019). To the best of our knowledge, the haloarchaeal coenzyme F420 biosynthesis pathway has never been experimentally analyzed.
(b) The prediction of coenzyme F420-specific oxidoreductases in *Mycobacterium* and actinobacteria has been reported (Selengut and Haft, 2010), leading to patterns and domains that are also found in haloarchaea. Several such enzymes are described in Suppl.Text S1 Section 4.
(c) HVO_1937 might be a coenzyme F420-dependent 5,10-methylenetetrahydrofolate reductase (see also below, C1 metabolism, and Suppl.Text S1 Section 4).
(d) The precursor for coenzyme F420 may be used by a photo-lyase involved in DNA repair.

**Table 4:**
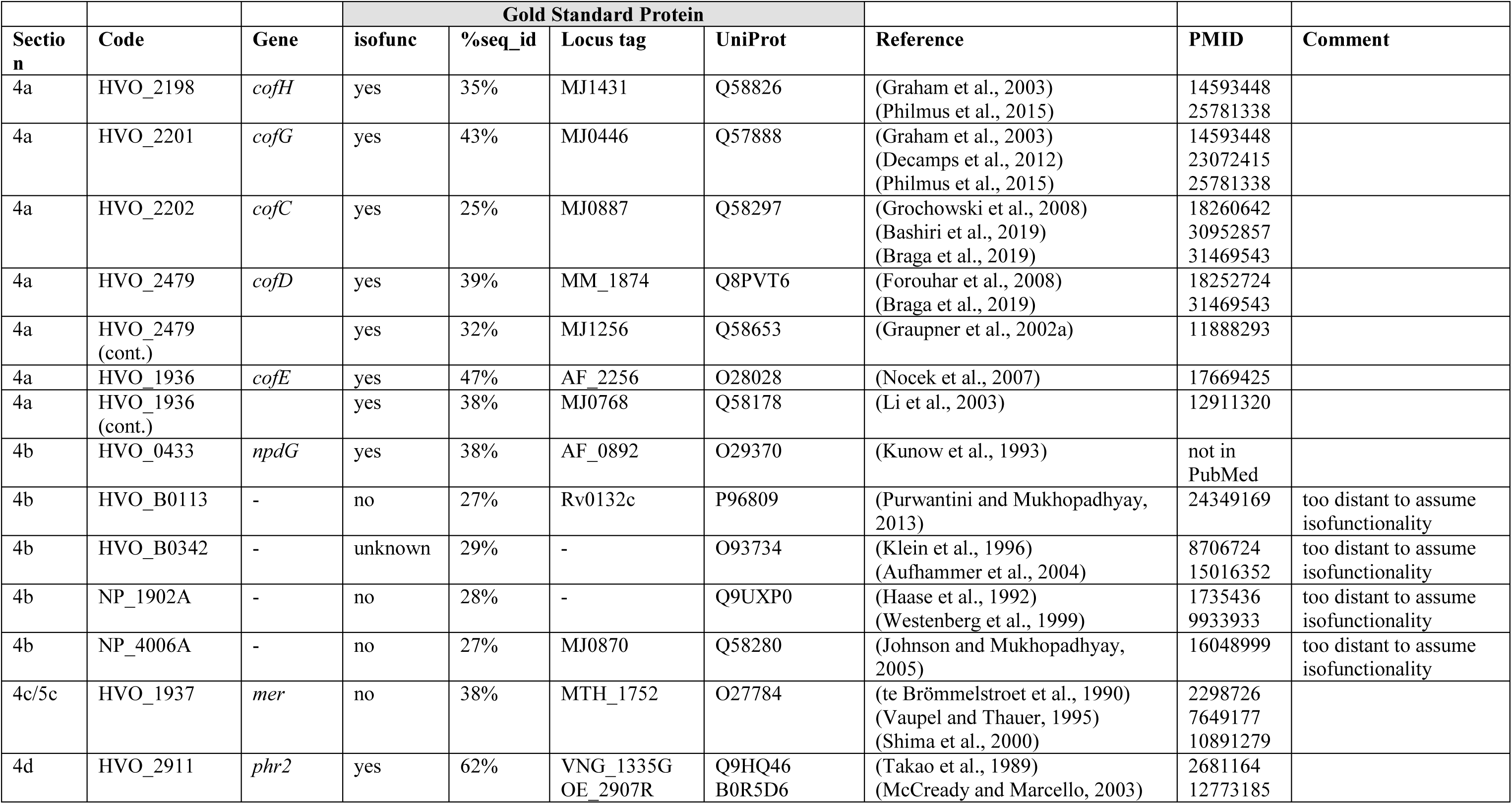
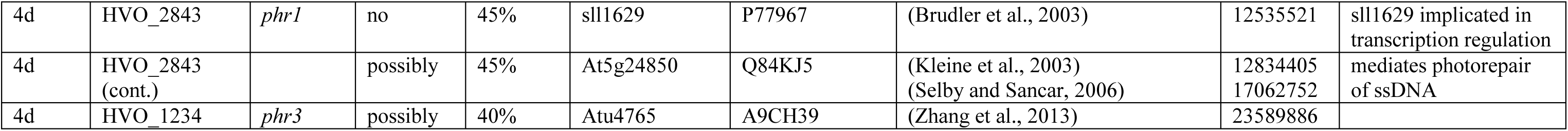
Proteins with open annotation issues and their Gold Standard Protein homologs (Section 4). For a description of this table see the legend to Table 1.

### 3.5 Coenzymes III: coenzymes of C1 metabolism: tetrahydrofolate in haloarchaea, methanopterin in methanogens

Halophilic and methanogenic archaea use distinct coenzymes as one-carbon carrier (C1 metabolism): tetrahydrofolate in haloarchaea and methanopterin in methanogens (White, 1988, Maden, 2000). Several characterized methanogenic proteins that act on or with methanopterin have comparably close homologs in haloarchaea (Table 5), which results in misannotation of haloarchaeal proteins (e.g. in SwissProt) as being involved in methanopterin biology. We assume that the haloarchaeal proteins function with the haloarchaeal one-carbon carrier tetrahydrofolate and that this shift in coenzyme specificity is possible due to the structural similarity between methanopterin and tetrahydrofolate (a near-identical core structure consists of a pterin heterocyclic ring linked via a methylene bridge to a phenyl ring; both also have a polyglutamate tail). A detailed review on the many variants of the tetrahydrofolate biosynthetic pathway is available (de Crecy-Lagard, 2014).

(a) Folate biosynthesis requires aminobenzoate. We had proposed candidates for a pathway from chorismate to para-aminobenzoate (Falb et al., 2008, Pfeiffer et al., 2008b) (for details see Suppl.Text S1 Section 5). However, these predictions have not been adopted by KEGG (accessed April 2021) and without experimental confirmation this is unlikely to ever happen.
(b) GTP cyclohydrolase MptA (HVO_2348) catalyzes a reaction in the common part of tetrahydrofolate and methanopterin biosynthesis. The enzymes specific for methanopterin biosynthesis are absent from haloarchaea and thus the assignment of HVO_2348 to the methanopterin biosynthesis pathway in UniProt is invalid (accessed March 2021).

**Table 5:**
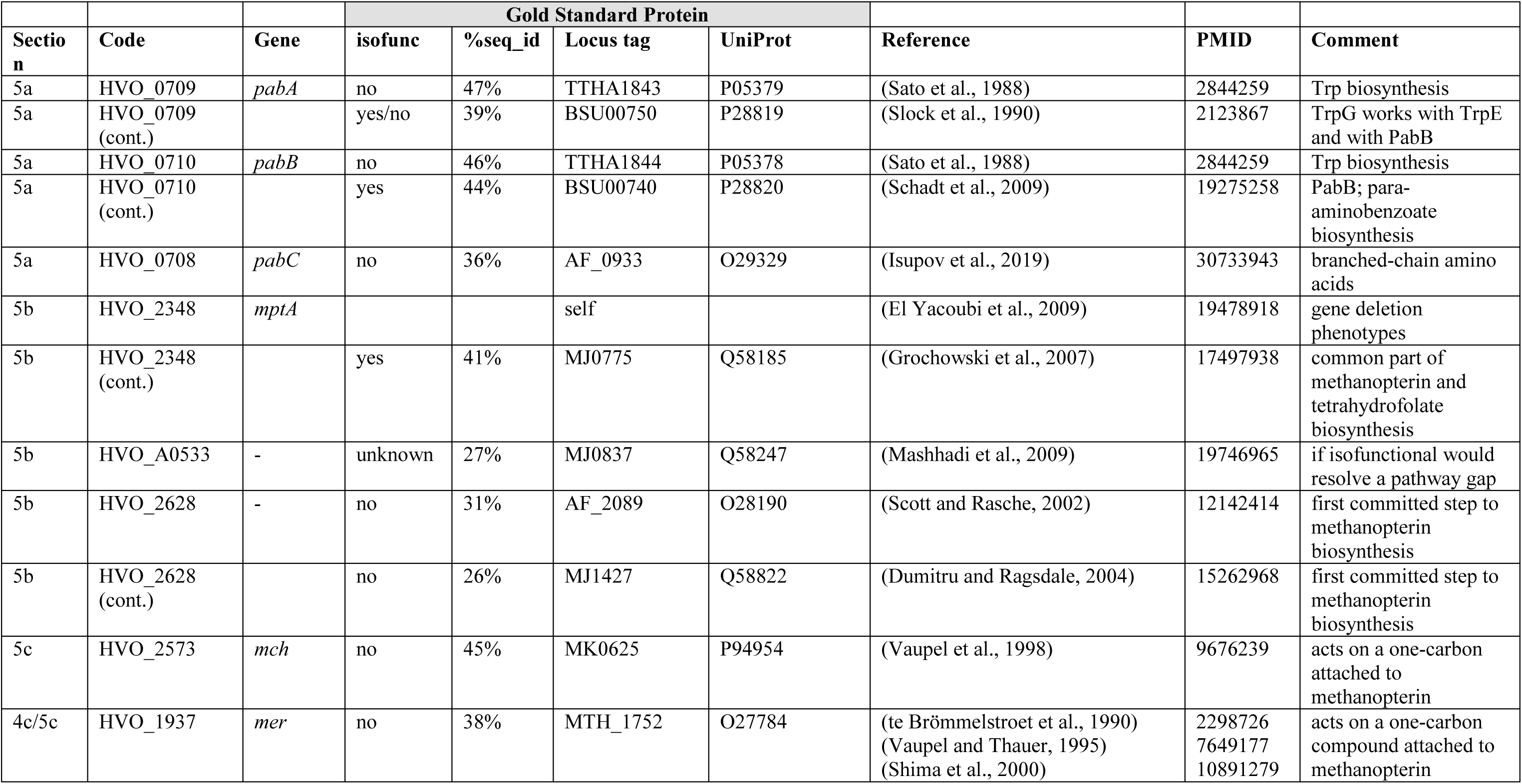
Proteins with open annotation issues and their Gold Standard Protein homologs (Section 5). For a description of this table see the legend to Table 1.

The next common pathway step (EC 3.1.4.56) has been resolved in *M. jannaschii* (MJ0837) but still is a pathway gap in halophilic archaea. MJ0837 is very distantly related to HVO_A0533, which thus is a promising candidate for experimental analysis.

HVO_2628 shows 30% protein sequence identity to the enzyme catalyzing the first committed step to methanopterin biosynthesis. As detailed in Suppl.Text S1 Section 5, we consider it likely that it does not catalyze that reaction.

(c) Two enzymes that alter the oxidation level of the coenzyme-attached one-carbon compound probably function with tetrahydrofolate, even though their methanogenic homologs function with methanopterin. In contrast to their assignments in KEGG and UniProt (as of March 2021), their probable functions are thus methenyltetrahydrofolate cyclohydrolase (HVO_2573) and 5,10-methylenetetrahydrofolate reductase (HVO_1937) (see Suppl.Text S1 Section 5)

### 3.6 Coenzymes IV: NAD and FAD (riboflavin)

(a) The energy source for NAD kinase may be ATP or polyphosphate. This is unresolved for the two paralogs of probable NAD kinase (HVO_2363, *nadK1*, HVO_0837, *nadK2*). These show only 25% protein sequence identity to each other (see Suppl. Text S1 Section 6). Polyphosphate was not found in exponentially growing *Hfx. volcanii* cells (Zerulla et al., 2014), so that ATP is the more likely energy source.
(b) HVO_0782 is an enzyme involved in NAD biosynthesis, which is encoded in most haloarchaeal and archaeal genomes. The adjacent gene, HVO_0781, is encoded in nearly all haloarchaeal genomes according to OrthoDB, and with very strong syntenic coupling revealed by SyntTax analysis. Thus, HVO_0781 is a candidate to also be involved in NAD biosynthesis. Characterized homologs to HVO_0781 decompose S-adenosyl-methionine into methionine and adenosine, a reaction that seems wasteful and might not be expected to be highly conserved with respect to existence and gene clustering (see Suppl. Text S1 Section 6).
(c) We describe the reconstruction of riboflavin biosynthesis based on a detailed bioinformatic reconstruction (Rodionova et al., 2017). The enzymes and their associated GSPs are listed in Table 6.Three pathway gaps remain, with candidate genes predicted for two of these (Rodionova et al., 2017) (for details see Suppl. Text S1 Section 6).

**Table 6:** Proteins with open annotation issues and their Gold Standard Protein homologs (Section 6). For a description of this table see the legend to Table 1.

| Section | Code | Gene | Gold Standard Protein |  |  |  | Reference | PMID | Comment |
| --- | --- | --- | --- | --- | --- | --- | --- | --- | --- |
|  |  |  | isofunc | %seq_id | Locus tag | UniProt |  |  |  |
| 6a | HVO_2363 | *nadK1* | unclear | 37% | Rv1695 | P9WHV7 | (Kawai et al., 2000) | 11006082 | can use ATP and PP |
| 6a | HVO_2363 (cont.) |  | unclear | 31% | AF_2373 | O30297 |  |  | ATP or PP usage unresolved |
| 6a | HVO_0837 | *nadK2* | unclear | 28% | Rv1695 | P9WHV7 |  |  | can use ATP and PP |
| 6a | HVO_0837 (cont.) |  | unclear | partial | AF_2373 | O30297 |  |  | ATP or PP usage unresolved |
| 6b | HVO_0782 | *nadM* | yes | 53% | MJ0541 | Q57961 | (Raffaelli et al., 1997)<br>(Raffaelli et al., 1999) | 9401030<br>10331644 |  |
| 6b | HVO_0781 | - | unknown | 42% | Sare_1364 | A8M783 | (Eustaquio et al., 2008) | 18720493 |  |
| 6b | HVO_0781 (cont.) |  | unknown | 35% | PH0463 | O58212 | (Deng et al., 2008) | 18551689 |  |
| 6b | HVO_0327 | *ribB* | yes | 43% | MJ0055 | Q60364 | (Fischer et al., 2002) | 12200440 |  |
| 6b | HVO_0974 | *ribH* | yes | 45% | MJ0303 | Q57751 | (Haase et al., 2003) | 12603336 |  |
| 6b | HVO_1284 | *arfA* |  | self |  |  | (Phillips et al., 2012) | 21999246 | gene deletion leads to riboflavin auxotrophy |
| 6b | HVO_1284 (cont.) |  | yes | 44% | MJ0145 | Q57609 | (Graham et al., 2002) | 12475257 |  |
| 6b | HVO_1235 | - | prediction |  |  |  | (Rodionova et al., 2017) | 28073944 | *arfB* candidate |
| 6b | HVO_1341 | *arfC* | yes | 36% | MJ0671 | Q58085 | (Graupner et al., 2002b)<br>(Romisch-Margl et al., 2008) | 11889103<br>18671734 |  |
| 6b | HVO_2483 | - | prediction | 34% | MJ0699 | Q58110 | (Rodionova et al., 2017) | 28073944 | predicted also for MJ0699 |
| 6b | pathway gap |  |  |  |  |  |  |  | EC 3.1.3.104 |
| 6b | HVO_0326 | *rbkR* | yes | 37% | TA1064 | Q9HJA6 | (Rodionova et al., 2017) | 28073944 | bifunctional as gene regulator and enzyme |
| 6b | HVO_0326 (cont.) |  | yes/no | 32% | MJ0056 | Q60365 | (Ammelburg et al., 2007) | 18073108 | enzyme only; lacks an N-terminal HTH domain |
| 6b | HVO_1015 | *ribL* | yes | 50% | MJ1179 | Q58579 | (Mashhadi et al., 2010) | 20822113 |  |

### 3.7 Biosynthesis of membrane lipids, bacterioruberin and menaquinone

Archaeal membrane lipids contain ether-linked isoprenoid side chains (see (Caforio and Driessen, 2017) and references cited therein). The isoprenoid precursor isopentenyl diphosphate is synthesized in haloarchaea by a modified version of the mevalonate pathway (Vannice et al., 2014). Isoprenoid units are then linearly condensed to the C20 compound geranylgeranyl diphosphate. The haloarchaeal core lipid, archaeol, consists of 2,3-sn-glycerol with two C20 isoprenoid side chains attached by ether linkages. In some archaea, especially alkaliphiles, C25 isoprenoids are also found (see e.g. (De and Gambacorta, 1988, Dawson et al., 2012)). Also, a number of distinct headgroups are found in polar lipids (phospholipids) (reviewed in (Caforio and Driessen, 2017)). Even though polar lipids are used as important taxonomic markers (Oren et al., 1997) their biosynthetic pathways are not completely resolved.

Haloarchaea typically have a red color, which is due to carotenoids, mainly the C50 carotenoid bacterioruberin (Oren, 2002, Kushwaha et al., 1975, Yang et al., 2015). For carotenoid biosynthesis, two molecules of geranylgeranyl diphosphate, a C20 compound, are linked head to head to generate phytoene, which is desaturated to lycopene (Falb et al., 2008, Giani et al., 2020). The pathway from lycopene to the C50 compound bacterioruberin has been experimentally characterized (Dummer et al., 2011, Yang et al., 2015).

(a) We assigned HVO_2725 (*idsA1*, paralog of NP_3696A) and HVO_0303 (*idsA2*, paralog of NP_0604A) for the linear isoprenoid condensation reactions resulting in a C20 isoprenoid (EC 2.5.1.10, 2.5.29, short chain isoprenyl diphosphate synthase) (see also Suppl.Text S1 Section 7). Some archaea, mainly haloalkaliphiles, also contain C25 isoprenoid side chains. Geranylfarnesyl diphosphate synthase, the enzyme which generates the C25 isoprenoids, has been purified and enzymatically characterized from *Nmn. pharaonis* (Tachibana, 1994), but data that allow the assignment to a specific gene have not been collected. Three paralogous genes from *Nmn. pharaonis* are candidates for this function (NP_0604A, NP_3696A, and NP_4556A). Because NP_0604A and NP_3696A have orthologs in *Hfx. volcanii*, a species devoid of C25 lipids, we assign the synthesis of C25 isoprenoids (geranylfarnesyl diphosphate synthase activity) to the third paralog, NP_4556A. UniProt assigns C25 biosynthesis activity to NP_3696A for undescribed reasons (as of April 2021) and KEGG does not make this assignment for any of the three paralogs (as of April 2021). Our assignments are supported by analysis of key residues which determine the length of the isoprenoid chain (Bale et al., 2019). These authors label the cluster containing NP_3696A (WP011323557.1) as “C15/C20” and the cluster containing NP_4556A (WP011323984.1) as “C20->C25->C30?”.
(b) Typical polar lipids in haloarchaea are phosphatidylglycerophosphate methyl ester (PGP-Me) and phosphatidylglycerol (PG), but also phosphatidylglycerosulfate (PGS) (Kates, 1993, Kates et al., 1993, Bale et al., 2019). Other polar lipids are archaetidylserine and its decarboxylation product archaetidylethanolamine, both of which are found in rather low quantities in *Haloferax* (Kellermann et al., 2016). A third group of polar lipids has a headgroup derived from myo-inositol. The biosynthetic pathway of the head groups is only partially resolved. One CDP-archaeol 1-archaetidyltransferase that belongs to a highly conserved three-gene operon may attach either glycerol phosphate or myo-inositol phosphate. In Suppl. Text S1 Section 7 we summarize arguments in favor of each of these candidates, but the true function can only be decided by experimental analysis.
(c) Carotenoid biosynthesis involves the head-to-head condensation of the C20 isoprenoid geranylgeranyl diphosphate to phytoene, which is desaturated to lycopene (Falb et al., 2008, Giani et al., 2020). The *crtB* gene product (e.g. HVO_2524) catalyzes the head-to-head condensation. It is yet uncertain which gene product is responsible for the desaturation of phytoene to lycopene. The further pathway from lycopene to bacterioruberin has been experimentally characterized in *Haloarcula japonica* (Yang et al., 2015). A three gene cluster (*crtD*-*lyeJ*-*cruF*) codes for the three enzymes of this pathway. The synteny of this three gene cluster is strongly conserved according to SyntTax analysis. Several genes which are certainly or possibly involved in carotenoid biosynthesis are encoded in the vicinity of this cluster (for details see Suppl. Text S1 Section 7).
(d) Halophilic archaea contain menaquinone as a lipid based two electron carrier of the respiratory chain (Elling et al., 2016, Kellermann et al., 2016). We describe the reconstruction of the menaquinone biosynthesis pathway (Table 7), with two pathway gaps remaining open (see Suppl.Text S1 Section 7 for details).

**Table 7:**
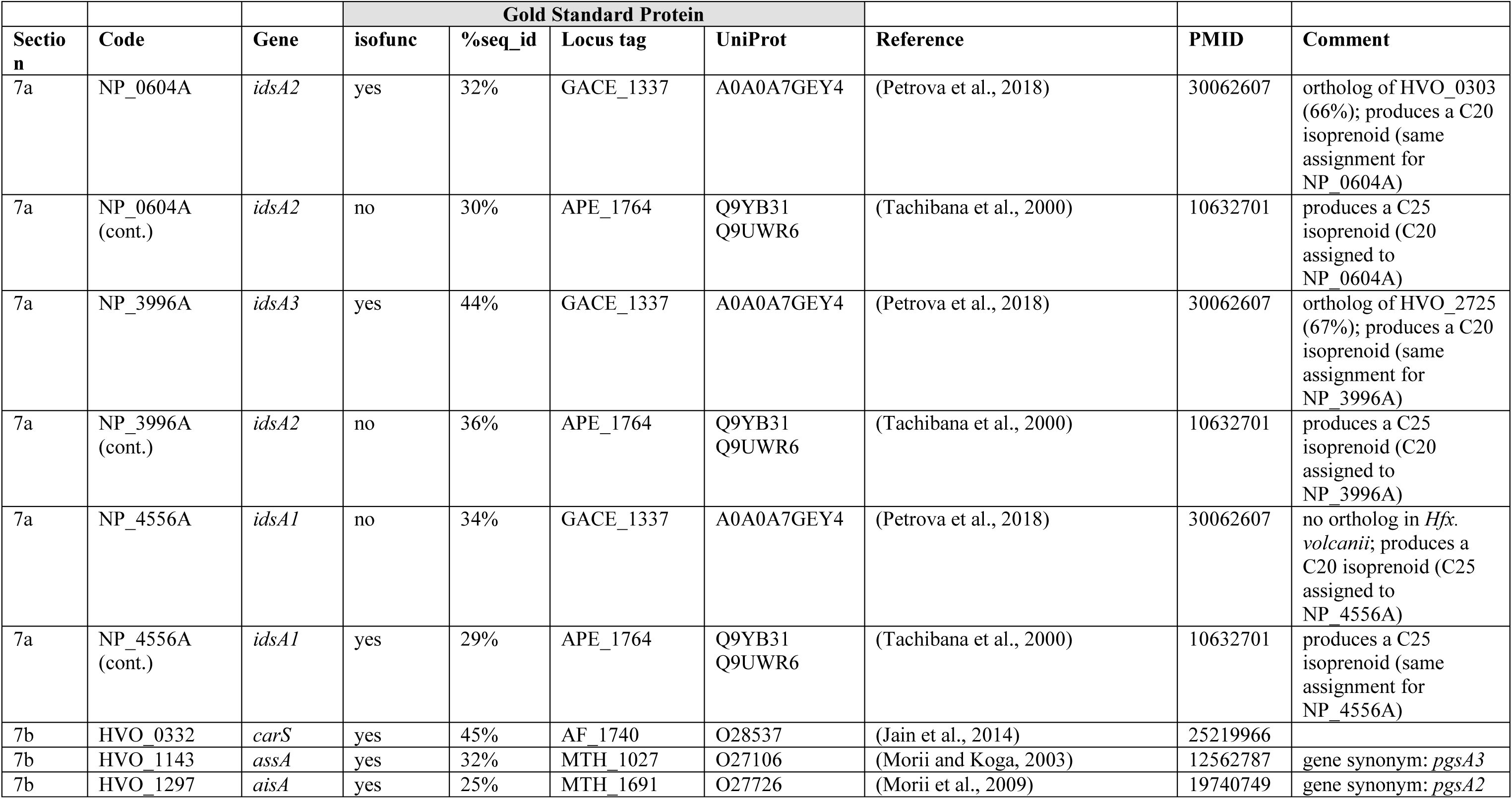
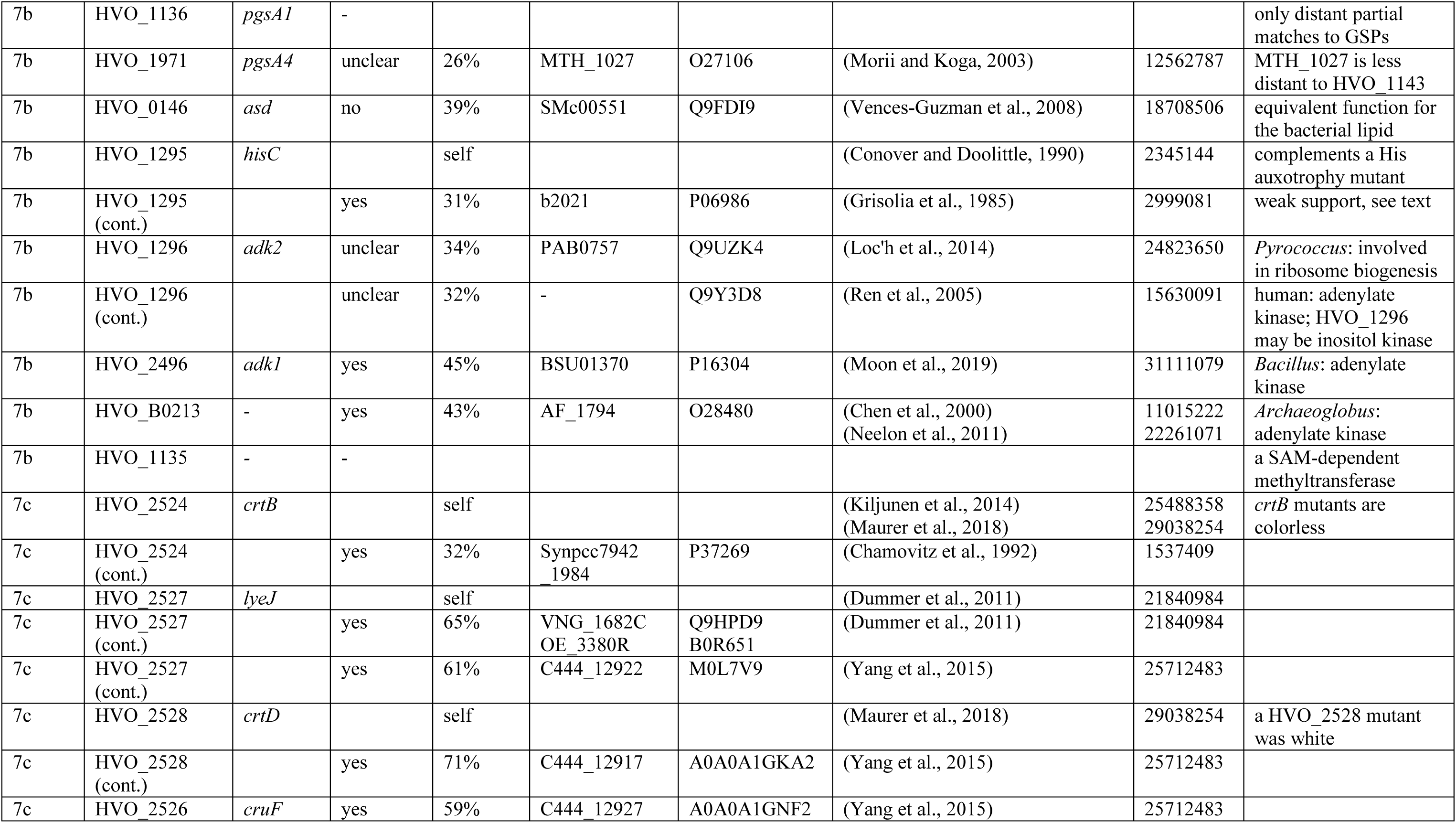
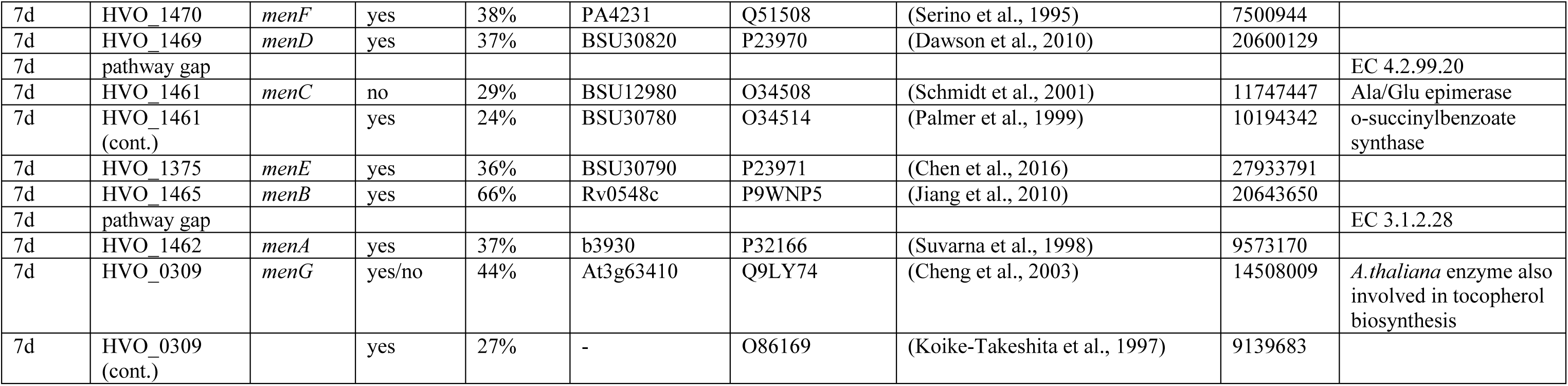
Proteins with open annotation issues and their Gold Standard Protein homologs (Section 7). For a description of this table see the legend to Table 1.

### 3.8 Issues concerning RNA polymerase, protein translation components and signal peptide degradation

(a) Haloarchaeal RNA polymerase consists of a set of canonical subunits (encoded by *rpoA1A2B1B2DEFHKLNP*). *Hbt. salinarum* and a subset of other haloarchaea contain an additional subunit called epsilon (Leffers et al., 1989, Madon and Zillig, 1983). Purified RNA polymerase containing the epsilon subunit transcribes native templates efficiently, in contrast to RNA polymerase devoid of this subunit (Madon and Zillig, 1983). The biological relevance of this subunit is enigmatic (see Suppl.Text S1 Section 8).
(b) Two distant paralogs are found for haloarchaeal ribosomal protein S10 (uS10) in nearly all haloarchaeal genomes. It is uncertain if both occur in the ribosome, whether they occur together or are mutually exclusive. The latter distribution would result in heterogeneity of the ribosomes. Alternatively, one of the paralogs may exclusively have a non-ribosomal function.

In a subset of archaea, two distant paralogs are found for haloarchaeal ribosomal protein S14 (uS14) (ca 20% of the genomes, e.g. in *Nmn. pharaonis*). For more details see Suppl.Text S1 Section 8.

(c) The ribosomal protein L43e (eL43) shows heterogeneity with respect to the present of C2-C2 type zinc finger motif. This zinc finger is found in L43e from all *Halobacteriales* and all euryarchaeal proteins outside the order *Halobacteria*, but is not found in *Haloferacales* and very rare in Natrialbales,. Eukaryotic orthologs (e.g. from rat and yeast) contain this zinc finger and its biological importance has been experimentally shown for the yeast protein (Rivlin et al., 1999) (for details see Suppl.Text S1 Section 8).
(d) Diphthamide is a complex covalent modification of a histidine residue of translation elongation factor a-EF2. The pathway has been reconstructed (Table 8), based on distant homologs (enzymes encoded by *dph2*, *dph5*) and by detailed bioinformatic analysis (enzyme encoded by *dph6*) (de Crecy-Lagard et al., 2012) (for details see Suppl.Text S1 Section 8). These uncertain function assignments await experimental confirmation.
(e) N-terminal signal sequences target proteins to the secretion machinery. Subsequent to membrane insertion or transmembrane transfer, the signal sequence is cleaved off by signal peptidase. After cleavage, the signal peptide must be degraded to avoid clogging of the membrane. Degradation is catalyzed by signal peptide peptidase. Candidates for this activity have been predicted from two protein families (Ng et al., 2007, Raut et al., 2021) (for details see Suppl.Text S1 Section 8).

**Table 8:** Proteins with open annotation issues and their Gold Standard Protein homologs (Section 8). For a description of this table see the legend to Table 1.

| Section | Code | Gene | Gold Standard Protein |  |  |  | Reference | PMID | Comment |
| --- | --- | --- | --- | --- | --- | --- | --- | --- | --- |
|  |  |  | isofunc | %seq_id | Locus tag | UniProt |  |  |  |
| 8a | OE_1279R | *rpoeps* |  | self |  |  | (Leffers et al., 1989)<br>(Madon and Zillig, 1983) | 2495365<br>6852054 |  |
| 8b | HVO_0360 | *rps10a* | yes | 94% | rrnAC2405 | P23357 | (Arndt et al., 1991) | 1764513 |  |
| 8b | HVO_1392 | *rps10b* | - |  |  |  |  |  | no GSP; 24% seq_id to HVO_0360 (*rps10a*) |
| 8b | NP_4882A | *rps14a* | yes | 72% | rrnAC1597.1 | P26816 | (Scholzen and Arndt, 1991) | 1832208 | full-length similarity; *Haloarcula* protein was not isolated or characterized |
| 8b | NP_4882A (cont.) |  | yes | 57% | YDL061C | P41058 | (Otaka et al., 1984) | 18782943 | yeast YS29B; N-term 20 aa divergent |
| 8b | NP_1768A | *rps14b* | unclear | 80% | rrnAC1597.1 | P26816 | (Scholzen and Arndt, 1991) | 1832208 | N-term 20 aa divergent |
| 8c | OE_1373R | *rpl43e* | yes | 69% | rrnAC1669 | P60619 | (Ban et al., 2000) | 10937989 |  |
| 8c | OE_1373R (cont.) |  | yes | 39% | YPR043W | P0CX25 | (Rivlin et al., 1999)<br>(Dresios et al., 2002) | 10588896<br>11866512 |  |
| 8c | HVO_0654 | *rpl43e* | yes | 54% | rrnAC1669 | P60619 | (Ban et al., 2000) | 10937989 | *Haloarcula*: has zinc finger;<br>*Haloferax*: lacks zinc finger |
| 8d | HVO_1631 | *dph2* | yes | 35% | PH1105 | O58832 | (Zhu et al., 2011) | 20931132 |  |
| 8d | HVO_0916 | *dph5* | yes | 39% | PH0725 | O58456 | (Zhu et al., 2010) | 20873788 |  |
| 8d | HVO_1077 | *dph6* | yes | 31% | YLR143W | Q12429 | (Su et al., 2012)<br>(Uthman et al., 2013) | 23169644<br>23468660 |  |
| 8e | HVO_0881 | *sppA1* | yes | 33% | BSU19530 | O34525 | (Bolhuis et al., 1999)<br>(Nam et al., 2012) | 10455123<br>22472423 |  |
| 8e | HVO_1987 | *sppA2* | probably | 23% | BSU19530 | O34525 | (Bolhuis et al., 1999)<br>(Nam et al., 2012) | 10455123<br>22472423 |  |
| 8e | HVO_1107 | - | prediction |  |  |  |  |  | no GSP |

### 3.9 Miscellaneous metabolic enzymes and proteins with other functions

Here, we list a few other enzymatic or non-enzymatic functions for which candidate genes have been assigned, but without experimental validation.

(a) Ketohexokinase from *Haloarcula vallismortis* has been experimentally characterized (Rangaswamy and Altekar, 1994). However, the activity was not assigned to a gene. Detailed bioinformatic analyses have been made (Anderson et al., 2011, Williams et al., 2019) and point to a small set of orthologs represented by Hmuk_2662, the ortholog of HVO_1812 (for further details see Suppl. Text S1 Section 9).
(b) The assignment of fructokinase activity to the *Hht. litchfieldiae* candidate gene halTADL_1913 (UniProt:A0A1H6QYL4) is based on differential proteomic analysis (Williams et al., 2019) (see Suppl. Text S1 Section 9 for details). Very close homologs are rare in haloarchaea. For this protein family (carbohydrate kinase) it is unclear if more distant homologs (with about 50% protein sequence identity) are isofunctional.
(c) A candidate gene for glucoamylase is HVO_1711 for reasons described in Suppl. Text S1 Section 9. The enyzme from *Halorubrum sodomense* has been characterized (Chaga et al., 1993) but the activity was not yet assigned to a gene.
(d) A strong candidate for having glucose-6-phosphate isomerase activity is *Hfx. volcanii* (HVO_1967, pgi), based on 36% protein sequence identity to the characterized enzyme from *M. jannaschii* (MJ1605) (Rudolph et al., 2004) (Table 9).
(e) A candidate gene for specifying an enzyme with 2-dehydro-3-deoxy-(phospho)gluconate aldolase activity is *Hbt. salinarum kdgA* (OE_1665R). It is rather closely related (36% protein sequence) to *Hfx, volcanii* HVO_1101 (encoded by *dapA*), which is involved in lysine biosynthesis, a biosynthetic pathway that is absent from *Hbt. salinarum*. The function assignment is based on distant homologs from *Saccharolobus (Sulfolobus) solfataricus* and *Thermoproteus tenax* which have been characterized (Ahmed et al., 2005) (for details see Suppl. Text S1 Section 9).
(f) Haloarchaea may contain an NAD-independent L-lactate dehydrogenase, LudBC (HVO_1692 and HVO_1693). Deletion of this gene pair impairs growth on rhamnose, which is catabolized to pyruvate and lactate (Reinhardt et al., 2019). There is a very distant relationship (for details see Suppl.Text S1 Section 9) to the LldABC subunits of the characterized L-lactate dehydrogenase from *Pseudomonas stutzeri* A1501 (Gao et al., 2015) and to the LutABC proteins from *B. subtilis*, which have been shown to be involved in lactose utilization (Chai et al., 2009).
(g) *Hfx. volcanii* may be able to convert urate to allantoin, using the gene cluster HVO_B0299-HVO_B0302. This could be part of a complete degradation pathway for purines, which, however, has to be considered highly speculative (see Suppl.Text S1 Section 9).
(h) *Hfx. volcanii* may contain an enzyme having a “nickel-pincer cofactor”. The biogenesis of this cofactor may be catalyzed by *larBCE* (as detailed in Suppl. Text S1 Section 9).
(i) cyclic diguanylate hydrolysis Cyclic di-AMP (c-di-AMP) is an important nucleotide signalling molecule in bacteria and archaea. It is generated from two molecules of ATP by diadenylate cyclase (encoded by *dacZ*) and is degraded to pApA by phosphodiesterases (Corrigan and Grundling, 2013). The level of this signalling molecule is strictly controlled (Gundlach et al., 2015, Commichau et al., 2019), thus requiring a sophisticated interplay of cyclase and phosphodiesterase. DacZ from *Hfx. volcanii* has been characterized and it was shown that c-di-AMP levels must be tightly regulated (Braun et al., 2019). The degrading enzyme, however, has not yet been identified in *Haloferax* but candidates have been proposed (Corrigan and Grundling, 2013, He et al., 2020, Yin et al., 2020) (see Suppl. Text S1 Section 9).
(j) homolog to RNaseZ HVO_2763 is distantly related to RNase Z (HVO_0144, *rnz*). The experimental characterization of HVO_2763 (Fischer et al., 2012) excluded activity as exonuclease but did not reveal its physiological function. Upon transcriptome analysis, the downregulation of several genes was detected. Several of these were uncharacterized at the time of the experiment but have since been shown to be involved in the minor N-glycosylation pathway that was initially detected under low salt conditions (see Suppl. Text S1 Section 9 for further details).
(k) A pair of genes (*dabAB*) is predicted to function as CO_2_ transporter. HVO_2410 and HVO_2411 probably function as carbon dioxide transporter, based on the identification of such transporters in *Halothiobacillus neapolitanus* (Desmarais et al., 2019). Being a member of the proton-conducting membrane transporter family, this protein may be misannotated as a subunit of the *nuo* or *mrp* complex (see Suppl. Text S1 Section 9 for further details).

**Table 9:** Proteins with open annotation issues and their Gold Standard Protein homologs (Section 9). For a description of this table see the legend to Table 1.

| Section | Code | Gene | Gold Standard Protein |  |  |  | Reference | PMID | Comment |
| --- | --- | --- | --- | --- | --- | --- | --- | --- | --- |
|  |  |  | isofunc | %seq_id | Locus tag | UniProt |  |  |  |
| 9a | HVO_1812 | - | prediction |  |  |  |  |  | no GSP |
| 9b | halTADL_1913 | - | yes | 37% | - | P26984 | (Aulkemeyer et al., 1991) | 1809835 |  |
| 9b | halTADL_1913 (cont.) | - | yes | 31% | OCC_03567 | Q7LYW8<br>H3ZP68 | (Qu et al., 2004) | 15138858 |  |
| 9c | HVO_1711 | - | probably | 33% | - | P29761 | (Ohnishi et al., 1992) | 1633799 | P29761 matches to C-term half of HVO_1711 |
| 9c | HVO_1711 (cont.) | - | probably | 51% | SAMN04487937_2677 | A0A1I6HD35 | (Chaga et al., 1993) | 8305855 | correlation between PMID:8305855 and A0A1I6HD35 likely (see text) |
| 9d | HVO_1967 | *pgi* | yes | 36% | MJ1605 | Q59000 | (Rudolph et al., 2004) | 14655001 |  |
| 9e | OE_1665R | *kdgA* | no | 31% | PA1010 | Q9I4W3 | (Kaur et al., 2011) | 21396954 | GSP for *dapA* (see under 2a) |
| 9e | OE_1665R (cont.) |  | probably | 30% | TTX_1156.1<br>TTX_1156a | G4RJQ2 | (Ahmed et al., 2005) | 15869466 |  |
| 9e | OE_1665R (cont.) |  | probably | 25% | SSO3197 | Q97U28 | (Ahmed et al., 2005) | 15869466 |  |
| 9f | HVO_1692 | *ludB* |  | self |  |  | (Reinhardt et al., 2019) | 30707467 |  |
| 9f | HVO_1692 (cont.) |  | probably | 35% | BSU34040 | O07021 | (Chai et al., 2009) | 19201793 | matches up to HVO_1692 pos 490 of 733 |
| 9f | HVO_1692 (cont.) |  | probably | 35% | PST_3338 | O4VPR6 | (Gao et al., 2015) | 25917905 | matches up to HVO_1692 pos 400 of 733 |
| 9f | HVO_1693 | *ludC* |  | self |  |  | (Reinhardt et al., 2019) | 30707467 |  |
| 9f | HVO_1693 (cont.) |  | probably | 30% | BSU34030 | O32259 | (Chai et al., 2009) | 19201793 |  |
| 9f | HVO_1693 (cont.) |  | probably | 33% | PST_3339 | O4VPR7 | (Gao et al., 2015) | 25917905 | partial match |
| 9f | HVO_1697 | - | unclear | 24% | PST_3340 | O4VPR8 | (Gao et al., 2015) | 25917905 |  |
| 9f | HVO_1696 | *lctP* | probably | 44% | PST_3336 | O4VPR4 | (Gao et al., 2015) | 25917905 |  |
| 9g | HVO_B0300 | *pucL1* | yes | 49% | BSU32450 | O32141 | (Pfrimer et al., 2010) | 20168977 | *Bacillus*: bifunctional, matches to C-term |
| 9g | HVO_B0299 | *pucM* | yes | 43% | BSU32460 | O32142 | (Lee et al., 2005) | 16098976 |  |
| 9g | HVO_B0301 | *pucL2* | yes | 43% | BSU32450 | O32141 | (Kim et al., 2007) | 17567580 | *Bacillus*: bifunctional, matches to N-term |
| 9g | HVO_B0302 | *pucH1* | no | 33% | - | Q8VTT5 | (Xu et al., 2002) | 12148274 | paper in Chinese, abstract in English; pyrimidine degradation |
| 9g | HVO_B0302 (cont.) |  | yes | 30% | STM0523 | Q7CR08 | (Ho et al., 2013) | 23287969 | purine degradation |
| 9g | HVO_B0302 (cont.) |  | yes | 29% | BSU32410 | O32137 | (Schultz et al., 2001) | 11344136 | purine degradation |
| 9g | HVO_B0306 | *amaB4* | no | 39% | - | Q53389 | (Martinez-Rodriguez et al., 2012) | 22904279 | carbamoyl-AA hydrolysis |
| 9g | HVO_B0306 (cont.) |  | yes | 34% | At5g43600 | Q8VXY9 | (Werner et al., 2010)<br>(Werner et al., 2013) | 19935661<br>23940254 | purine degradation |
| 9g | HVO_B0308 | *coxS* | no | 46% | Saci_2270 | Q4J6M5 | (Kardinahl et al., 1999) | 10095793 | GAPDH |
| 9g | HVO_B0308 (cont.) |  | no | 41% | - | P19915 | (Kang and Kim, 1999) | 10482497 | CO-DH |
| 9g | HVO_B0308 (cont.) |  | yes | 39% | b2868 | Q46801 | (Xi et al., 2000) | 10986234 | xanthine DH |
| 9g | HVO_B0309 | *coxL* | yes | 33% | b2866 | Q46799 | (Xi et al., 2000) | 10986234 | xanthine DH |
| 9g | HVO_B0309 (cont.) |  | no | 28% | - | P19913 | (Kang and Kim, 1999) | 10482497 | CO-DH |
| 9g | HVO_B0309 (cont.) |  | no | 26% | Saci_2271 | Q4J6M3 | (Kardinahl et al., 1999) | 10095793 | GAPDH |
| 9g | HVO_B0310 | *coxM* | no | 31% | Saci_2269 | Q4J6M6 | (Kardinahl et al., 1999) | 10095793 | GAPDH |
| 9g | HVO_B0310 (cont.) |  | no | 31% | - | P19914 | (Kang and Kim, 1999) | 10482497 | CO-DH |
| 9g | HVO_B0310 (cont.) |  | yes | 25% | b2867 | Q46800 | (Xi et al., 2000) | 10986234 | xanthine DH |
| 9g | HVO_B0303 | *uraA4* | yes | 38% | b3654 | P0AGM9 | (Karatza and Frillingos, 2005) | 16096267 |  |
| 9h | HVO_0197 | - | possibly | 39% | lp_0105 | F9UST0 | (Desguin et al., 2016) | 27114550 | LarB family protein |
| 9h | HVO_2381 | - | possibly | 31% | lp_0106 / | F9UST1 | (Desguin et al., 2016) | 27114550 | LarC family protein |
|  |  |  |  |  | lp_0107 |  |  |  |  |
| 9h | HVO_0190 | - | possibly | 34% | lp_0109 | F9UST4 | (Desguin et al., 2016) | 27114550 | LarE family protein |
| 9i | HVO_1660 | *dacZ* |  | self |  |  | (Braun et al., 2019) | 30884174 |  |
| 9i | HVO_0756 | - | prediction |  |  |  | (He et al., 2020) | 32095817 |  |
| 9i | HVO_0990 | - | prediction |  |  |  | (He et al., 2020) | 32095817 |  |
| 9i | HVO_1690 | - | prediction |  |  |  | (He et al., 2020) | 32095817 |  |
| 9j | HVO_2763 | - |  | self |  |  |  | 22350204 | no function could be assigned |
| 9j | HVO_2763<br>(cont.) |  | no | 27% | HVO_0144 | D4GZ88 | (Spath et al., 2008) | 18437358 | Rnase Z |
| 9k | HVO_2410 | *dabA* | yes | 33% | Hneap_0211 | D0KWS7 | (Desmarais et al., 2019) | 31406332 |  |
| 9k | HVO_2411 | *dabB* | yes | 31% | Hneap_0212 | D0KWS8 | (Desmarais et al., 2019) | 31406332 |  |

## 4. Conclusion

We describe a large number of cases where protein function cannot be correctly predicted when restricting considerations to computational analyses without taking the biological context into account. An example is the switch from methanopterin to tetrahydrofolate as C1 carrier in haloarchaea. Homologous enzymes, inherited from the common ancestor, have adapted to the new C1 carrier, rather being replaced by non-homologous proteins. Function prediction tools may misannotate haloarchaeal proteins to work with methanopterin. Another example is the *nuo* complex and its misannotation as a type I NADH dehydrogenase. In other cases, even distant sequence similarity may allow a valid function prediction if additional evidence (e.g. from gene neighbourhood analysis or from detailed evaluation of metabolic pathway gaps) is taken into account. Examples include cobalamin cluster proteins, which probably close the two residual pathway gaps, and the predicted degradation pathway for purines. In all these cases, we have presented reasonable hypotheses based on current knowledge, and in many cases these are so well supported as to be compelling, but to be certain, experimental data are required. With this overview, we attempt to arouse the curiosity of colleagues, hoping that they will confirm or disprove our speculations and thus advance the knowledge about haloarchaeal biology. *Hfx. volcanii* is a model species for halophilic archaea, and the more complete and correctly its genome is annotated, the higher will be its value for systems biology analyses (modelling) and for synthetic biology (metabolic engineering) and biotechnology.

## Supporting information

Supplemental Text S1 provides detailed information

## Acknowledgments

We thank all members of the *Haloferax* community for more than a decade of fruitful cooperation and for sharing their deep knowledge about the biology of this model species. We thank our colleagues for kindly reading and providing thoughtful comments on the manuscript before submission: Sonja-Verena Albers, Thorsten Allers, Maria Jose Bonete, Rosana de Castro, Sebastien Ferreira-Cerca, Anita Marchfelder, Mechthild Pohlschroder, Joerg Soppa. We thank Birgit Scharf for insightful discussions on the *Natronomonas* respiratory chain.

## Data Availability Statement

None.

## Funding

This research received no specific grant from any funding agency in the public, commercial, or non-for-profit sectors.

## Authors contribution

**Conceptualization**, F.P.

**Data curation**, F.P.

**Project administration**, F.P.

**Formal analysis**, F.P., M.D-S.

**Investigation** F.P., M.D-S.

**Validation**, F.P., M.D-S.

**Visualization**, F.P., M.D-S.

**Writing – original draft**, F.P., M.D-S.;

**Writing – review & editing**, F.P., M.D-S.

## Conflict of interest

None declared.

## Ethics statement

Not applicable

## Notes

### Competing Interest Statement

The authors have declared no competing interest.

