## Supplemental Text S1 provides detailed information for "Open issues for protein function assignment in *Haloferax volcanii* and other halophilic archaea"

### SUPPLEMENTARY TEXT S1: Detailed background information for all open annotation issues

This text is organized so as to correspond with the decimal-numbered subsections (.1, .2, .3 etc.) of the Results segment (3) of the main text. For example, details for 3.2 Amino Acid Biosynthesis are in Section S2, and S2.c covers topic (c) in the results.

#### Section S1: The respiratory chain and oxidative decarboxylation

##### S1.a Ferredoxin-dependent oxidative decarboxylation

In haloarchaea, oxidative decarboxylation is not linked to reduction of NAD to NADH but to reduction of a 2Fe-2S ferredoxin (encoded by *fdx*) (e.g. OE_4217R, HVO_2995) which has a redox potential similar to that of the NAD/NADH pair (Kerscher and Oesterhelt, 1977). The enzymes for oxidative decarboxylation are pyruvate--ferredoxin oxidoreductase (*porAB*) and 2-oxoglutarate--ferredoxin oxidoreductase (*korAB*), and these have been characterized from *Halobacterium salinarum* (Plaga et al., 1992, Kerscher and Oesterhelt, 1981b, Kerscher and Oesterhelt, 1981a). In *Haloferax*, a conditionally lethal *porAB* mutant was unable to grow on glucose or pyruvate, but could grow on acetate. This demonstrates that alternative enzymes for conversion of pyruvate to acetyl-CoA do not exist in this species (Kuprat et al., 2021).

##### S1.b Re-oxidation or reduced ferredoxin

In *Haloferax*, the same ferredoxin (HVO_2995) plays an essential role in nitrate assimilation (Zafrilla et al., 2011), but additional biological roles might be uncovered by further experimental analyses. It is yet unclear how the reduced ferredoxin Fdx (HVO_2995) is reoxidized. We speculate that the haloarchaeal Nuo complex may be responsible for ferredoxin reoxidation.

##### S1.c The haloarchaeal Nuo complex

Mitochondrial complex I functions as NADH dehydrogenase, transferring two electrons to the lipid electron carrier ubiquinone (Rich and Marechal, 2010). The equivalent in *Escherichia coli* is the Nuo complex which functions as NADH dehydrogenase (Sousa et al., 2012, Kaila and Wikstrom, 2021). Haloarchaeal genomes contain a *nuo* cluster with genes and gene order largely conserved to the gene set in *E. coli*. Apart from a few gene fissions and fusions, the haloarchaeal *nuo* cluster is characterized by the absence of the three genes *nuoEFG*. The NuoEFG gene products function as a NADH adaptor and make this complex a NADH dehydrogenase (Leif et al., 1995, Braun et al., 1998). Thus, the haloarchaeal Nuo complex is unlikely to reoxidize NADH, even though databases, e.g. KEGG (as of April 2021), assign this function. Our rejection of the haloarchaeal Nuo complex being annotated as NADH dehydrogenase is supported by experimental data which show that the haloarchaeal complex I is not involved in NADH oxidation (Sreeramulu et al., 1998). Up to now, the substrate of the Nuo complex in haloarchaea has remained enigmatic but reduced ferredoxin Fdx is a reasonable candidate. The thermophilic cyanobacterium *Thermosynechococcus elongatus* has an equivalent complex which also lacks the *nuoEFG* genes and was shown to directly mediate oxidation of ferredoxin, which is reduced by photosystem I in this species (Zhang et al., 2005, Schuller et al., 2019, Pan et al., 2020).

**S1.d Type II NADH dehydrogenase in halophilic archaea**

Halophilic archaea do not use a type I enzyme for NADH oxidation but instead oxidize NADH via a type II NADH dehydrogenase (Sreeramulu et al., 1998). For *Natronomonas pharaonis*, we have predicted that NP_3508A functions as type II NADH dehydrogenase (Falb et al., 2005). The *Hfx. volcanii* ortholog is HVO_1578 (59% protein sequence identity). However, this assignment is highly questionable.

The assignment of NP_3508A as the *Nmn. pharaonis* type II NADH dehydrogenase was based on the enzyme from *Acidianus ambivalens* which at that time was considered to be a characterized type II NADH dehydrogenase (Gomes et al., 2001, Bandeiras et al., 2002). There is, however, only 26% protein sequence identity, and this is even restricted to the N-terminal 140 of ca 400 aa. Even more problematic is the fact that the function assignment to the very distant homolog has subsequently been corrected and the enzyme is now known to be a sulfide:quinone oxidoreductase devoid of NADH dehydrogenase activity (Brito et al., 2009).

HVO_1578 and NP_3508A are distantly related to *E. coli ndh* but with only 24% protein sequence identity. *E. coli ndh* has been characterized as a type II NADH dehydrogenase (Jaworowski et al., 1981). NP_3508A is somewhat more closely related (30% protein sequence identity) to *Alkalihalobacillus (Bacillus) pseudofirmus* NDH-2A which has been functionally characterized (Liu et al., 2008). The identification of the type II NADH dehydrogenase in halophilic archaea has to be considered questionable and can only be clarified by experimental characterization. The assignment of type II NADH dehydrogenase activity to HVO_1578/NP_3508A and to HVO_1413) was also made in an extensive analysis of the NADH:quinone oxidoreductase family (Marreiros et al., 2016).

**S1.e A canonical complex III is absent in many haloarchaea**

While the overwhelming majority of haloarchaea code for the equivalent of complexes I and II (Nuo complex and succinate dehydrogenase), about one-third do not code for a complex III equivalent (cytochrome $bc_1$ complex encoded by *petABC*) according to OrthoDB analysis. While the $bc_1$ complex is found in most *Haloferacales* and *Halobacteriales*, it is absent in nearly all *Natrialbales*. However, it was also not found in the genus *Natronomonas,* which belongs to the *Halobacteriales*. In *Hbt. salinarum,* the genes for the three subunits (*petABD*) are clustered in the genome (OE_1876R/1874R/1872R) while in *Hfx volcanii* the *petBD* genes (HVO_0842/0841) are clustered and *petA* (HVO_2620) is solitary and separated by 1.1Mb.

The $bc_1$ complex is required to transfer electrons from the lipid-embedded two-electron carrier (menaquinone in haloarchaea) to the one-electron carrier associated with terminal oxidases (probably halocyanin). Most likely, electrons flow via an alternative pathway, which, however, is yet unresolved.

The haloarchaeal genes share some characteristics with those of the chloroplast b6-f complex rather than those of the mitochondrial $bc_1$ complex. In chloroplasts and in haloarchaea, the cytochrome b gene is split (chloroplast genes code for cytochrome b6, PetB, and the 17K subunit, PetD). As an example, *Haloferax* PetB shows 39% protein sequence identity to cytochrome b6 from *Synechococcus sp.* PCC7002 (Lee et al., 2001). Also, the associated one electron carrier in chloroplasts is a copper protein (plastocyanin) as in haloarchaea (probably halocyanin, see below) and not a heme (and thus iron) protein (mitochondrial cytochrome-c).

**S1.f An atypical "bc complex" from *Natronomonas pharaonis* and the claim that a novel way of covalent heme attachment may exist in haloarchaea**

An atypical cytochrome bc has been isolated from *Nmn. pharaonis* (Scharf et al., 1997). While canonical $bc_1$ complexes have a 2:1 ratio of b-type to c-type cytochrome, the *Natronomonas* complex had an unusual 1:1 stoichiometry (Scharf et al., 1997). The enzyme was characterized but data allowing its assignment to a gene were not integrated into that publication. However, the authors mention unpublished amino acid composition data: “No Cys could be determined during the analysis of the amino acid composition after performic acid oxidation (data not shown), but since heme C is covalently bound via two cysteine residues to the protein, the hydrolysis product would escape the detection by amino acid analysis”.

Sequence data are available from a PhD thesis (Mattar, 1996). Analyses attributed to two diploma theses resulted in a long nearly contiguous protein sequence (41 amino acids). In one experiment, the band from the SDS-PAGE was subjected to N-terminal protein sequencing. In the other attempt, the protein complex was digested with proteases (Glu-C, chymotrypsin) and the peptides were separated by HPLC. A peak absorbing at 400 nm (in addition to 280 nm) was considered to contain covalently attached heme and was subjected to protein sequencing. Two overlapping peptides were obtained in this experiment, and these also overlapped with the sequence obtained by N-terminal sequencing. In total, a region of 41 amino acids was sequenced with only the penultimate residue remaining unassigned. From this partial protein sequence, degenerated primers were designed and could be used successfully for PCR amplification (Mattar, 1996). The PCR product was used as probe to identify and clone a genomic restriction fragment coding for the corresponding gene. The translation of the gene was near-identical to the protein sequencing result, except for two mismatches (Mattar, 1996). The gene identified was *sdhD*, and was part of a four-gene operon coding for the four subunits of succinate dehydrogenase (respiratory complex II). The product of *sdhD* is one of the membrane subunits. Succinate dehydrogenase itself was characterized in the same paper that also described the cytochrome bc, and was found to contain b-type heme (Scharf et al., 1997). We list all subunits of this enzyme from *Hfx. volcanii* and *Nmn. pharaonis* in Suppl. Table S10.

Expecting the identified gene to code for a sequence with two Cys residues (see above), the authors were surprised to find that SdhD lacks cysteine residues, and concluded that their research strategy had failed and they had inadvertently isolated an unrelated gene. This conclusion is expressed in the PhD thesis as “Looking at the remainder of the translated sequence of ORF 3, there is neither the described heme C binding motif nor any cysteine residue” and “nevertheless, the lack of a heme c binding motif and any cysteine in the translated sequence indicates that the isolated gene cannot represent a cytochrome-c”.

A similar attitude is conveyed in a recent review on haloarchaeal heme proteins (Kletzin et al., 2015) under the heading "The Haloarchaea Paradox". After summarizing descriptions of small cytochrome-c proteins in *Natronomonas*, *Halobacterium*, and *Haloferax*, the authors state, "This leads to the conclusion that they might not be found using similarity and/or pattern searches and that they use non-standard amino acid patterns and heme c linkage".

We speculate that all the described experiments in the Mattar thesis had been successful. In our view, it is well possible that haloarchaeal c-type cytochromes exemplify a novel type of heme attachment which is independent of Cys residues. In this context, it should be noted that the translation of the SdhD gene over the 41 residues that are covered by the protein sequencing results contains three His residues, none of which was identified upon Edman degradation. Two of the His residues correspond to the two mismatches, while the third corresponds to the residue which could not be called.

It is textbook knowledge and a paradigm that heme must be covalently attached via a pair of Cys residues. However, a paradigm shift may be necessary if haloarchaea have developed a cysteine-independent heme attachment mechanism. We hope that our detailed analysis helps to pave the way for an experimental challenge to the existing paradigm.

Next, we consider if the described small cytochrome-c from *Halobacterium* could also correspond to SdhD. This cytochrome $c_{552}$ is reported to be 14.1 kDa. It was purified and covalent heme attachment was shown by heme staining after SDS-PAGE (Sreeramulu, 2003). The original authors considered their cytochrome-c to be the functional equivalent of mitochondrial cytochrome-c (Sreeramulu et al., 1998, Sreeramulu, 2003). The *Halobacterium* $c_{552}$ is likely to correspond to the *Natronomonas* cytochrome bc for a number of reasons: (a) the *Halobacterium* cytochrome $c_{552}$ was purified from membranes which had been solubilized with Triton X-100. The mitochondrial and bacterial cytochrome-c which functions as one-electron carrier to cytochrome-c oxidase is a soluble and monomeric protein. (b) The *Halobacterium* cytochrome $c_{552}$ contained considerable amounts of a b-type heme. The authors state that hydroxyapatite chromatography "resolved the preparation into fractions with different ratios of cytochromes" (Sreeramulu, 2003), which was interpreted as partially separating distinct proteins but an equally consistent interpretation is that a single protein complex might have undergone some degree of disaggregation. (c) The size of the c-type heme containing protein is in the same range (14.1 kDa vs 18 kDa), and the difference might be attributed to a comparably low size resolution for small proteins and the performance of the experiments in different laboratories. (d) A gene coding for a protein homologous to cytochrome-c has never been identified in any haloarchaeal genome. Taking all these points into consideration, there is good evidence to believe that *Halobacterium* $c_{552}$ and *Natronomonas* cytochrome bc are corresponding proteins, and are both SdhD.

**S1.g Halocyanin as candidate one electron carrier for cba-type terminal oxidase**

The last step of the respiratory electron transport chain is catalyzed by terminal oxidase. There are two classes of terminal oxidases. Some terminal oxidases act directly on the lipid based two electron carrier, which is the product of complexes I and II. The majority of terminal oxidases, however, function with a one electron carrier. The classical (mitochondrial) one electron carrier is the heme protein cytochrome-c. In halophilic archaea, the one electron carrier is most likely halocyanin, a copper containing protein related to plastocyanin from chloroplasts.

A halocyanin from *Nmn. pharaonis* (NP_3954A) has been characterized, including its redox potential (Mattar et al., 1994, Scharf and Engelhard, 1993, Hildebrandt et al., 1994). It is a protein which is TATLipo positive (Storf et al., 2010) and thus is secreted by the TAT pathway, followed by signal peptidase II cleavage and N-terminal lipid attachment to the conserved cysteine of the lipobox motif. Protein-chemical characterization identified the attached lipid and provided circumstantial evidence of the N-terminus being additionally acetylated (Mattar et al.,

1994, Scharf and Engelhard, 1993). The redox potential would be consistent with it being an electron donor to the cba-type terminal oxidase, but it is unknown if this halocyanin is the cognate electron donor.

The *cbaD* subunit of the cba terminal oxidase is a very short protein in *Natronomonas* (NP_2966A, 54 aa) (Mattar and Engelhard, 1997). In *Halobacterium*, this domain is fused with a pair of blue copper domains (OE_4073R, 429 aa). In strain NRC-1, one of these domains has been cut out with surgical precision, while both domains are found in strains R1 and the type strain 91-R6 of *Halobacterium* (Pfeiffer et al., 2008, Pfeiffer et al., 2020). In *Haloferax*, the Cba type terminal oxidase has a short CbaD subunit (HVO_0943, 97 aa) and no halocyanin gene is found in the genomic vicinity of the *cba* gene cluster. Experimental confirmation of the involvement of halocyanin in terminal oxidation by cba-type terminal oxidases is, to our knowledge, yet unavailable. The *Haloferax* halocyanin HVO_2150 is the most closely related to the N-terminal part of OE_4073R, and is thus a candidate to be the electron donor to the cba-type terminal oxidase.

**S1.h Terminal oxidases in haloarchaea**

Mitochondrial complex IV is the terminal oxidase which transfers electrons from the one electron carrier cytochrome-c to oxygen, finally resulting in the generation of water (Rich and Marechal, 2010, Guo et al., 2018). Halophilic archaea code for a diverse set of terminal oxidases (Schaefer et al., 1996, Falb et al., 2005, Refojo et al., 2019). Some probably function with one electron carriers, similar to those of mitochondria. However, the one electron carrier is not a heme protein (cytochrome-c) but a copper protein (halocyanin). It cannot be excluded that some halophilic archaea code for terminal oxidases which are able to directly reoxidize the two electron carrier menaquinone.

In three haloarchaea (*Nmn. pharaonis*, *Hfx. volcanii*, *Hbt. salinarum*) at least one terminal oxidase has been experimentally studied and we restrict our analysis to terminal oxidases in these species. All three species code for additional, yet uncharacterized, terminal oxidases, which are also covered.

*Nmn. pharaonis* codes for three terminal oxidases, of which one belongs to the $ba_3$-type (encoded by *cbaABDE*) and has been experimentally characterized (Mattar and Engelhard, 1997). Each subunit has an ortholog in *Hfx. volcanii*, which includes a short ortholog of CbaD and an ortholog of CbaE. The long subunits have an ortholog in *Hbt. salinarum*. This species has CbaD fused to a halocyanin with a pair of blue copper domains (OE_4073R) (see above, halocyanin). An ortholog of CbaE is missing in *Hbt. salinarum*. The subunit structure in *Natrialba magadii*, which has two copies of this terminal oxidase, corresponds to that of *Nmn. pharaonis*, while the subunit structure in *Haloarcula marismortui* corresponds to that of *Hbt. salinarum*.

Another terminal oxidase of *Nmn. pharaonis* belongs to the $aa_3$-type and consists of three subunits (according to InterPro domain assignments) which are located nearby on the genome (CoxA1, NP_2456A, CoxB1, NP_2448A, CoxC1, NP_2450A). Its long chain (subunit I, CoxA1) is orthologous to the long chain of the characterized $aa_3$-type terminal oxidases of *Hfx. volcanii* (CoxA1, HVO_0907, 68% protein sequence identity) and to CoxA1 of *Hbt. salinarum* (OE_1979R, 65% protein sequence identity). The characterized enzymes from these species were probably partially disintegrated upon solubilization and purification as only two subunits were found for the *Haloferax* enzyme (only CoxA1 sequenced and thus assigned to a gene) (Tanaka et al., 2002) and only a single subunit was found for the *Halobacterium* enzyme (Fujiwara et al., 1989, Denda et al., 1991). However, there are orthologs to *Natronomonas* CoxB1 and CoxC1 in both species. In *Halobacterium*, the genes for the three subunits are located nearby on the genome (CoxA1, OE_1979R, CoxB1, OE1988R, CoxC1, OE1984F). In *Haloferax*, the three subunits are separated from each other (CoxA1, HVO_0907, CoxB1, HVO_1014, CoxC1, HVO_1138). In all three species, the *coxB1* gene is directly adjacent to that of heme o synthase (encoded by *ctaB*).

*Hfx. volcanii* and *Hbt. salinarum* also contain a cytochrome bd ubiquinol oxidase (encoded by *cydAB*) which is absent from *Nmn. pharaonis*. CydA is distantly related to the characterized enzyme from *Azotobacter vinelandii* and slightly more distant to that of *E. coli*. CydAB directly oxidizes the intramembrane two-electron carrier to the corresponding quinone (ubiquinone in *E. coli* and *A. vinelandii*, menaquinone in haloarchaea).

*Hfx. volcanii* codes for a fourth terminal oxidase, which does not have an ortholog in the other two species, exemplifying the variability of respiratory chains in haloarchaea. It resembles the characterized $aa_3$-type terminal oxidase of *Aeropyrum pernix* (Ishikawa et al., 2002). This terminal oxidase is characterized by the fusion of the genes for subunits 1 and 3 (encoded by *coxAC2*, HVO_1645), a subunit 2 (encoded by *coxB2*, HVO_1646) and an additional small subunit (encoded by *coxD2*, HVO_1644).

*Nmn. pharaonis* also codes for another terminal oxidase (*coxA3*, NP_4296A, *coxB3*, NP_4294A) which does not have an ortholog in *Hbt. salinarum* or *Hfx. volcanii*. This is very distantly related to CbaAB (NP_2962A/2964A). The closest SwissProt homolog (from *Thermus thermophilus*) is more closely related to the cognate CbaAB. Among the 16 haloarchaeal genomes which are under survey (Pfeiffer and Oesterhelt, 2015) (Suppl. Table S11), only *Halobacterium hubeiense* has orthologs with >55% protein sequence identity.

**S1.j NAD-dependent oxoacid dehydrogenase complexes**

In *E. coli*, pyruvate is oxidized to acetyl-CoA by the classical pyruvate dehydrogenase complex, which results in NADH that is subsequently reoxidized by complex I (NADH dehydrogenase).

In halophilic archaea, pyruvate is oxidized to acetyl-CoA exclusively via a ferredoxin-dependent mechanism, while NAD(P)-dependent pyruvate oxidation could not be detected in cell homogenates (Kerscher and Oesterhelt, 1977). The enzyme (pyruvate--ferredoxin oxidoreductase; EC 1.2.7.1; OE2622R+OE2623R) has been characterized (Plaga et al., 1992). An equivalent reaction oxidizes alpha-ketoglutarate to succinyl-CoA (oxoglutarate--ferredoxin oxidoreductase; EC 1.2.7.3; OE1711R+OE1710R) (Kerscher and Oesterhelt, 1981b, Kerscher and Oesterhelt, 1981a). Further confirmation that NAD-dependent enzymes are not involved in conversion of pyruvate to acetyl-CoA was obtained by deletion of the *Haloferax porAB* genes.

The deletion strain loses the ability to grow on glucose or pyruvate as a single carbon source, but still can grow on acetate (Kuprat et al., 2021).

Haloarchaeal genomes code for distant homologs of the classical pyruvate dehydrogenase complex. Because NAD was experimentally shown not to be involved in oxidative decarboxylation of pyruvate or alpha-ketoglutarate, these enzymes must act as 2-oxoacid dehydrogenases using substrates other than pyruvate and alpha-ketoglutarate. In *Haloferax*, several studies have attempted to unravel the substrate, but only with partial success. For example, in one study (Jolley et al., 2000), the authors wrote: "Enzyme activities: In the context of our discovering genes for a putative ODHC" (Oxoacid DeHydrogenase Complex), "it is worth reiterating that we and others have never detected enzymic activities of PDHC" (Pyruvate DeHydrogenase Complex), "OGDHC" (OxoGlutarate (which is alpha-ketoglutarate) DeHydrogenase Complex) "or the BCODHCs" (Branched Chain Oxoacid DeHydrogenase Complex) in *H. volcanii* or any other halophilic archaeon. However, DHLipDH" (dihydrolipoate dehydrogenase) "activity is present in all strains tested. Assays for the ODHCs have been repeated with the strain of *H. volcanii* used in this work and the absence of enzymatic activity has been reconfirmed. Positive controls for these activities have been performed with cell extracts of *E. coli*." Several other publications have attempted to unravel the substrate of the oxoacid dehydrogenase complexes by analyzing three of the four E1 components without any success (van Ooyen and Soppa, 2007, Al-Mailem et al., 2008, Wanner and Soppa, 2002). Not only were enzyme assays attempted, but also the growth of *Haloferax* with pyruvate and/or alpha-ketoglutarate as sole carbon source was analyzed. Only one publication successfully identified the substrate of one of the oxoacid dehydrogenase complexes (Sisignano et al., 2010). For three of the four genes known at that time to encode an E1 alpha subunit, a number of gene deletions were generated: single gene deletions, all pairs of double deletions, and a triple deletion. Growth on isoleucine was impaired in all deletion strains lacking the *oadh1* gene (HVO_2958). Metabolomic analyses showed that these strains failed to metabolize isoleucine as well as its deamination product while the parent strain and those deletion strains that retain the *oadh1* gene were unaffected. However, even though the combination of experimental results makes the substrate assignment unambiguous, it was not possible to establish an enzyme assay for *Haloferax*, even though the activity of the positive control used in their assays (*Pseudomonas* *putida* branched-chain OADHC) could be readily measured. The *Haloferax* oxoacid dehydrogenase complexes seem notoriously difficult to study.

The identification of closely related experimentally characterized homologs (GSPs) is not straight-forward due to database incompleteness. HVO_2958 (encoded by *oadA1*) and HVO_2209 (encoded by *oadA4*) are equidistant to a characterized homolog from *Thermoplasma acidophilum* (38% protein sequence identity). That enzyme is involved in the degradation of all three branched chain amino acids (Heath et al., 2007), while HVO_2958 was proposed to be specific for Ile and HVO_2209 has not yet been characterized as its existence could not be anticipated prior to sequencing of the complete genome. The *Thermoplasma* proteins are still unreviewed in UniProt (as of April 2021), but a bibliography submission has been provided using the newly introduced community feedback system of UniProt. HVO_0669 (encoded by *oadA3*) is related to the E1 component of a TPP-dependent acetoin dehydrogenase from *Bacillus subtilis* (54% protein sequence identity) and from *Pelobacter carbinolicus* (49% protein sequence identity), both being characterized (Huang et al., 1999, Oppermann et al., 1991). These are also top matches for HVO_2595 (encoded by *oadA2*) (40-41% protein sequence identity), but only the *B. subtilis* homolog is incorporated into the SwissProt section of UniProt (accessed April 2021). Given database incompleteness, it cannot be excluded that more closely related characterized homologs exist.

Despite strong evidence which excludes pyruvate and alpha-ketoglutarate as substrates, KEGG insists (despite corresponding feedback) to maintain the annotation of these proteins as pyruvate dehydrogenase (EC 1.2.4.1) (accessed April 2021). According to our correspondence with KEGG, this will only be discontinued when the haloarchaea community is able to provide positive identification of the true biological substrate.

To provide further information, we list in Suppl. Table S10 all components of oxoacid dehydrogenase complexes, the alpha and beta chain of component E1, component E2 including the short (86 aa) lipoyl domain protein, and the single component E3. We thus highlight gene clusters and indicate sequence similarity between paralogs.

#### Section S2: Amino acid metabolism

##### S2.a Reconstruction of arginine and lysine biosynthesis

Arginine biosynthesis starts from glutamate with phosphorylation of the gamma-carboxylic group. To avoid unfavorable intramolecular circularization, the alpha-amino group must be protected, with the key intermediate metabolite ornithine being released by deprotection. The protecting group is part of the enzyme name for the pathway reactions between glutamate and ornithine, and thus, a reliable annotation requires a correct assignment of the protection mode. We are confident that haloarchaea use a carrier-mediated approach for arginine biosynthesis, but there is yet no experimental proof. The assignment is based on experimentally characterized homologs, which, however, are additionally or even exclusively involved in lysine biosynthesis.

*Thermococcus kodakarensis*, *Sulfolobus acidocaldarius* and *T. thermophilus* all synthesize lysine via the alpha-aminoadipate pathway using a carrier protein for protection of the alpha-amino group. Thus, they use a carrier protein (encoded by *lysW*) and a ligase (encoded by *lysX*). In *Thermococcus*, all enzymes of the arginine and lysine biosynthesis pathway are bifunctional (promiscuous) and are involved in both pathways (Yoshida et al., 2016). In *Sulfolobus*, the same carrier protein (LysW) is used for both, lysine and arginine biosynthesis (Ouchi et al., 2013), but the ligases which attach the substrate are distinct (being encoded by *argX* and *lysX*, respectively) (Ouchi et al., 2013). In *Thermus*, the LysX/LysW pair has been characterized (Horie et al., 2009). In UniProt, this reference has been mis-associated with the proteins from strain HB8 instead of HB27 (accessed March 2021). The LysW protein from strain HB27 (TT_C1544, Q72HE5) has the codon for Val- 23 incorrectly assigned to be the start codon. After sequence extension, there is 57% protein sequence identity to HVO_0047.

Halophilic archaea do not use the alpha-aminoadipate pathway for lysine biosynthesis but instead use the diaminopimelate pathway (Hochuli et al., 1999), which does not require protection of the alpha-amino group. The genes for carrier protein and ligase are encoded in the arginine biosynthesis cluster. We thus assign gene names *argW* and *argX* rather than *lysW* and *lysX* (as in *Thermococcus*, *Sulfolobus*, and *Thermus*). However, experimental confirmation of these predictions is not yet available. Also, no genes have been assigned for conversion of alpha-ketoglutarate to alpha-aminoadipate in haloarchaea, which would be a prerequisite for involvement of the *arg* cluster genes in lysine biosynthesis via the alpha-aminoadipate pathway. It should be noted that KEGG also does not assign two of the four enzymes for *T. kodakarensis* (accessed April 2021), even though *Thermococcus* is known to use that pathway (Yoshida et al., 2016).

Because most of the enzymes for arginine biosynthesis are related to bifunctional enzymes, we provide the reconstruction of the complete pathway in Table 2, where genes are arranged along the biosynthesis pathway according to the enzymes they encode. It has been observed experimentally that the level of arginine biosynthesis enzymes decreases with increased temperature (Jevtic et al., 2019). Additionally, we provide an extensive reconstruction of the lysine biosynthesis pathway in haloarchaea in Table 2. There is one gap in the pathway (EC 2.6.1.17, *dapC*, *argD*). One of the generally annotated pyridoxal phosphate-dependent aminotransferases may code for this reaction, but there is none in the genomic vicinity. Several haloarchaea lack homologs to *dapF* and/or to *dapE*. It should also be noted that KEGG assigns DapE function also to HVO_A0634 (accessed March 2021). We consider this unlikely because it's genomic position is not adjacent to other genes of the pathway, and because it shows low similarity to a GSP.

**S2.b Aromatic amino acid biosynthesis**

*Methanocaldococcus jannaschii* enzymes MJ0400 and MJ1249 have been functionally characterized and shown to be involved in the archaea specific part of the shikimate pathway for aromatic amino acid biosynthesis (2-amino-3,7-dideoxy-D-threo-hept-6-ulosonate synthase, EC 2.2.1.10 and 3-dehydroquinate synthase, EC 1.4.1.24) (White, 2004). Orthologs in halophilic archaea (encoded by *fba2*) are OE_1472F from *Hbt. salinarum* and HVO_0790 from *Hfx. volcanii* for MJ0400 and OE_1475F and HVO_0792 for MJ1249. OE_1472F and OE_1475F have been partially characterized (Gulko et al., 2014). Gene deletion was not possible but transcription could be strongly diminished (Gulko et al., 2014). This resulted in poor growth in the absence of aromatic amino acids, supporting the involvement in aromatic amino acid biosynthesis. According to the pathway in *M. jannaschii*, MJ0400 should condense aspartate semialdehyde with 6-deoxy-5-ketofructose (DKFP). Based on circumstantial evidence (see below), an alternative reaction is proposed where aspartate semialdehyde is condensed with fructose-1,6-bisphosphate (F-1,6-P) rather than DKFP. This would result in an alternative product that, however, could be converted to an identical compound, 3-dehydroquinate, in the subsequent step, because the part where the molecules differ would not be incorporated into the product. Reasons to propose a different substrate are: (a) difficulties in reconstructing a pathway leading to DKFP in *Halobacterium* and (b) the ability to detect binding of fructose-1,6-bisphosphate to MJ0400 by X-ray structural studies which involve soaking of the crystal with fructose-1,6-bisphosphate (Morar et al., 2007) and (c) the observation by White (White, 2004) that DKFP, incubated with labelled glucose, “only slightly decreased the extent of label incorporation into the products”. White concludes “This result would indicate that that DKFP may not be the preferred precursor to Compound 1”. F-1,6-P has been tested only in combination with labelled pyruvate and with L-homoserine (no product formation with either), but not in combination with aspartate semialdehyde.

Unfortunately, the establishment of an in vitro assay for the condensation of F-1,6-P with aspartate semialdehyde for the *Halobacterium* protein was not successful. Thus, the proposed alternative reaction for haloarchaeal biosynthesis of aromatic amino acids remains unconfirmed. A clean deletion of the *Haloferax* ortholog HVO_0790 has been generated (Maupin-Furlow, personal communication), but this strain has not been further characterized. Thus, the initial reactions in aromatic amino acid biosynthesis are a persisting open issue which awaits experimental confirmation.

**S2.c Tryptophan degradation**

The gene for tryptophanase (*tnaA*) is stringently regulated in *Haloferax*, which is the basis for using its promoter in the genetic toolbox for regulated gene expression (Large et al., 2007). Clearly, tryptophan should only be degraded when available in the medium. Tryptophanase can then cleaves Trp to $NH_3$, pyruvate, and indole. However, the fate of the resulting indole remains unknown. None of the indole conversions represented in KEGG are assigned to a protein from *Hfx. volcanii* (as of April 2021).

**S2.d Histidine utilization**

A probable histidine utilization cluster exists, based on distant homologs from *B. subtilis* (Kaminskas et al., 1970, Yu et al., 2006, Oda et al., 1988, Hartwell and Magasanik, 1963). The set of four clustered genes (HVO_A0562 – HVO_A0559) would code for a complete pathway that converts histidine to glutamate. To our knowledge, growth of *Hfx. volcanii* on histidine has not yet been characterized and the enzymes have not yet been experimentally verified. Previously we had observed an enhanced level of enzymes involved in histidine utilization under high-salt conditions (Jevtic et al., 2019).

**S2.e Auxotrophic mutants from a transposon insertion library with yet unassigned function**

Among 16 auxotrophic mutants observed in a *Hfx. volcanii* transposon insertion mutant library (Kiljunen et al., 2014), some could grow only in the presence of one (or several) supplied amino acids. Besides disruption of genes already known to be involved in the biosynthesis of the required amino acid (e.g. *argB*, *argC, carA*), some of the targeted genes do not yet have an assigned function. These are listed here.

Targeting of HVO_0431 (HAD superfamily hydrolase) makes *Haloferax* auxotrophic for histidine. This fits well to what currently is considered the only remaining pathway gap of histidine biosynthesis (histidinol-phosphatase). However, the enzyme catalyzing the preceding step (HVO_1295) had its function assigned because it can complement a histidine auxotrophic mutation and shows distant similarity to *E. coli* histidinol-phosphate aminotransferase (EC 2.6.1.9) encoded by *hisC* (31% protein sequence identity). This level of sequence identity does not allow substrate assignments for this protein family (pyridoxal-phosphate dependent aminotransferase). To the best of our knowledge, it has not been experimentally confirmed if HVO_1295 catalyzes the reaction of EC 2.6.1.9. The gene coding for HVO_1295 is part of a highly conserved three-gene operon, with at least one of the other members being involved in polar lipid biosynthesis (see below, Section 7). It cannot be excluded that the complementation of histidine auxotrophy is due to a side activity of an enzyme with reduced substrate specificity which primarily functions in polar lipid biosynthesis. In that case, even, it might be possible that haloarchaea do not use the classical histidine biosynthesis pathway where the product of EC 4.2.1.19 (imidazole-acetol phosphate) is first transaminated (by EC 2.6.1.9, histidinol-phosphate aminotransferase) and then dephosphorylated (by EC 3.1.3.15, the pathway gap described above). These reactions might be swapped in haloarchaea. Only experimental analysis can clarify whether HVO_1295 functions as histidinol-phosphate aminotransferase and HVO_0431 functions as histidinol-phosphatase.

HVO_0644 is annotated as 2-isopropylmalate synthase / (R)-citramalate synthase (EC 2.3.3.13 and EC 2.3.1.182, respectively). The two reactions are chemically similar and thus could be catalyzed by an enzyme with relaxed substrate specificity. (R)-citramalate synthase is an early step of isoleucine biosynthesis and transposon insertion into HVO_0644 makes *Haloferax* auxotrophic for isoleucine (Kiljunen et al., 2014). It is yet unresolved if HVO_0644 (encoded by *leuA1*) is really bifunctional, having 2-isopropylmalate synthase activity and thus being involved in leucine biosynthesis. This activity is assigned to the distant paralog HVO_1510 (encoded by *leuA2*, 39% protein sequence identity). HVO_1510 belongs to an ortholog set for which the start codon assignment is very difficult (see below, S2.f).

Targeting of HVO_1153 makes *Haloferax* auxotrophic for proline. The characteristics of that protein provide no link to proline biosynthesis. The protein has an N-terminal transmembrane domain but tools to detect secretion signals (Sec, Tat, Fla) are negative. The protein may be O-glycosylated (positive with NetOGlyC) and thus may belong to the secretome. Thus, this protein is currently annotated as "probable secreted glycoprotein", the same annotation being used for the adjacently encoded HVO_1154 and HVO_1155. In this case, a standard gene deletion should be generated to confirm the phenotype of the transposon insertion library. It should be noted that transposon insertion mutants are relatively frequently associated with secondary genome alterations (Collins et al., 2020), and the observed phenotypes might be caused by these rather than the transposon insertion event.

**S2.f Difficulty in finding a phylogenetically consistent start codon for a conserved leucine biosynthesis gene may indicate that the translation start is atypical**

HVO_1510 is annotated as 2-isopropylmalate synthase (EC 2.3.3.13) (also in KEGG as of April-2021), which is the first reaction specific for leucine biosynthesis, just after that pathway branches off from valine biosynthesis. This activity is also assigned as one of two activities to

HVO_0644, the other being a chemically similar reaction (EC 2.3.1.182) of isoleucine biosynthesis.

HVO_1510 belongs to an ortholog set with a major difficulty in start codon assignment. Each of the genomes under survey (Suppl. Table S11) has an ortholog (with more than 68% protein sequence identity), sequence similarity starting between pos 27 and 45. The most N-terminal peptide of HVO_1510 which was identified in the ArcPP project (Schulze et al., 2020) is a tryptic peptide starting at pos 62, the next in-frame start codon (when accepting ATG, GTG, or TTG) is the codon for Met-93. Given the high coverage obtained for that protein, it is remarkable that the 18 aa peptide from pos 43-61 has not been identified, which might indicate that this peptide does not exist in the protein. Other peptides in this region cannot be considered as they are too small for proteomic detection, a consequence of the large number of arginine residues. The genomes from *Haloferax gibbonsii* strains LR2-5 (CP063205) and ARA6 (CP011947) are near-identical at the DNA and protein level but only strain LR2-5 has a start codon (GTG) assignment consistent with *Hfx. volcanii* HVO_1510 while in ARA6 the corresponding codon is GTA (Figure S2). This makes it very unlikely that the currently assigned GTG is truly the start codon of HVO_1510. The genome of *Haloferax mediterranei* is near-identical at the DNA level starting from base 23 but lacks similarity to bases 1-22. The region from pos 24 to 91 has a near-identical DNA sequence but contains two in-frame stop codons in *Hfx. mediterranei* (Figure S2). Thus, the ortholog HFX_1573 is annotated as a disrupted gene, starting with Leu-27 (having a CTG, not a TTG codon) but being N-terminally truncated. Further examples are provided in Figure S2, where the *Hfx. volcanii* DNA sequence (codons 3 to 58, centered at codon 27) is compared to the genomes from 20 other strains from the genus *Haloferax*. *Hfx. mediterranei* is not exceptional in this respect, but the described pattern is also found in several other strains from the genus *Haloferax*. In Figure S2, deviating bases are highlighted by coloring, and it is evident that the region upstream of codon 27 is much more diverse than the region thereafter. The conserved sequence starts with a strictly retained CTG codon, which would code for Leu if an internal codon and for Met if used as an atypical start codon. We also note a strictly retained out-of-frame ATG close to the start of the conserved region. Its usage as translation start would necessitate ribosomal slippage, which might be employed for translational regulation. A potential transcription start was mapped ca 10 nt after codon 27, but the applied method is prone to limited positional uncertainty. The comparison to genera other than *Haloferax* is possible at the protein sequence level due to strong sequence conservation while there is a higher diversity at the DNA sequence level. Sequence similarity to orthologs from *Haloquadratum*, *Haloarcula* and *Natrialba* starts with Cys-30 of HVO_1510, which corresponds to Cys-10 of HQ277A/Hqrw_3051, Cys-8 of rrnAC0329/HAH_1067, and Cys-18 of Nmag_0906. Similarity starts even later for orthologs from *Halobacterium*. Glu-45 of HVO_1510 corresponds to Glu-7 of Hhub_3057 from *Hbt. hubeiense* and Phe-46 corresponds to Phe-6 of HBSAL_05970 from strain 91-R6$^T$ of *Hbt. salinarum*. The orthologs from *Natronomonas* are closely related from Arg-36 of HVO_1510 (Arg-4 of NP_2206A and Nmlp_3164) but lack an upstream start codon (the next alternative being the codon for Met-60). The ortholog from *Halohasta litchfieldiae* starts with Cys-30, also lacks an upstream start codon, and the next alternative start codon is that for Met-63. It can be considered highly unlikely that this gene is disrupted in such a large number of haloarchaea. The other genes of the leucine specific part of the biosynthetic pathway are encoded in those genomes, including the gene for the first such enzyme that would have independently become nonfunctional. Thus, it is tempting to speculate that this gene is functional and that it is translated from an atypical start codon. This would allow for a regulatory mechanism where translation occurs only under conditions of an insufficient leucine supply. Conserved in-frame CTG and out-of-frame ATG codons were observed by DNA sequence comparison to other genomes from *Haloferax* (Figure S2) but experimental analysis will be needed to clarify the true start codon used.

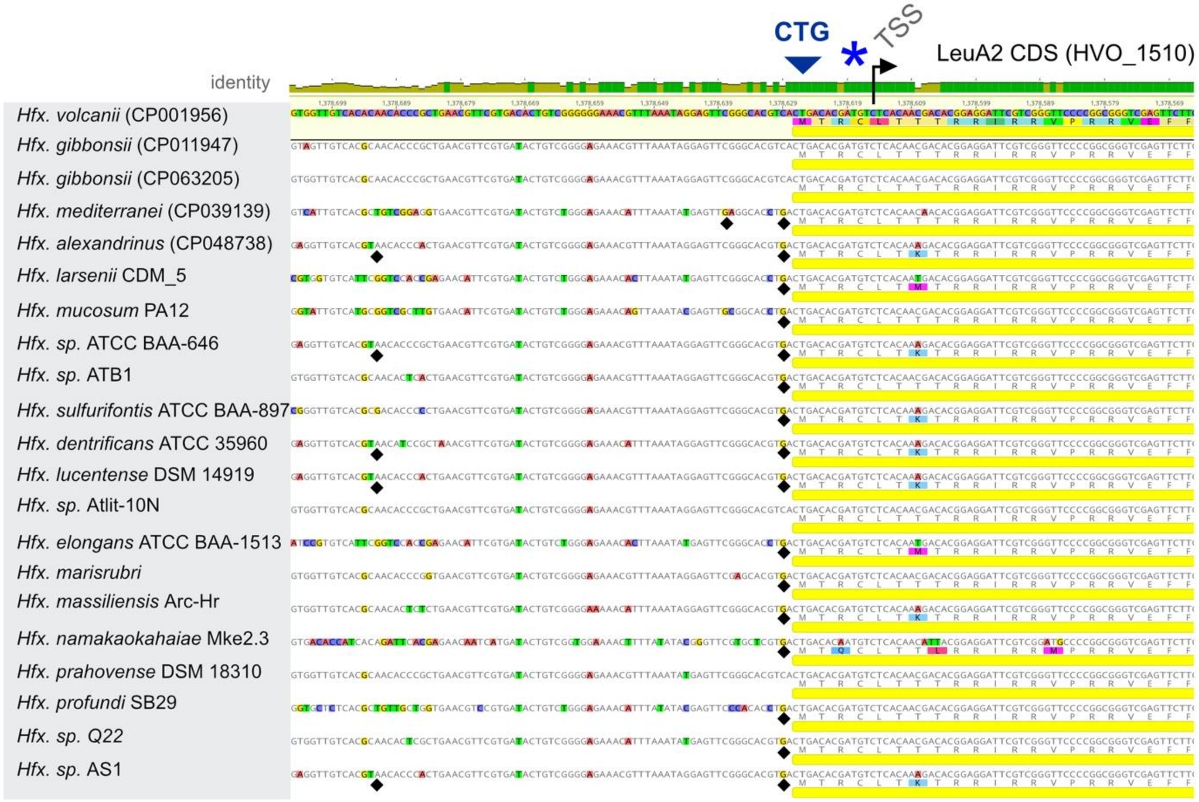


**Figure S2. Alignment of the 5’ end of *leuA2* (HVO_1510) from *Hfx. volcanii* DS2 with homologs from other species of *Haloferax*.** Translation given in single letter code. In the comparison sequences below *Hfx. volcanii*, colored bases (and amino acids) indicate where differences occur. It is evident that colored bases are more frequent in the left half of the alignment. The dark blue arrow marks a conserved CTG codon which might serve as a start codon (tentatively translated to Met, corresponds to Leu-27 of HVO_1510 as currently annotated; the currently assigned start codon is the GTG at the beginning of the sequence). Black diamonds show positions of in-frame stop codons, which are found exclusively upstream of the CTG. The light blue asterisk marks a conserved, out of frame ATG. Its usage as a start codon would necessitate ribosomal slippage, which might be used for translation regulation. The arrow marks the transcription start site (TSS) of *leuA2* as determined by (Laass et al., 2019). Even though the CTG is 10-12 bp upstream and the ATG is 2-4 bp upstream of the TSS, either of these codons may still be potential start codons given that the method for TSS determination carries some positional uncertainty.

#### Section S3: Coenzymes I: cobalamin and heme

Halophilic archaea use the "alternative heme biosynthesis pathway" (Bali et al., 2011, Kuhner et al., 2014). This pathway has been reconstructed (Siddaramappa et al., 2012), with few issues remaining open (see below).

De novo cobalamin biosynthesis is a multistep process, which occurs in two variants for the conversion of precorrin-2 to the corrin ring intermediate cobyrinate a,c diamide (Blanche et al., 1993). The cobalt-late (aerobic) pathway uses cobalt-free substrates, while the cobalt-early pathway uses cobalt-containing substrates. Many of the pathway intermediates are equivalent except for the absence/presence of the central cobalt atom, and the enzymatic reactions thus correspond closely to each other and are typically catalyzed by homologous enzymes. An exception is a key reaction which is chemically distinct, being catalyzed by CbiG (cobalt-early) or CobG (cobalt-late). The central cobalt atom plays an important role in the reaction catalyzed by CbiG (Moore et al., 2013). In the cobalt-late (aerobic) pathway, the reaction is catalyzed by CobG and involves oxidation by molecular oxygen. The presence of a transition metal (cobalt) may not be suitable for this type of reaction.

The reactions beyond ligand cobyrinate a,c diamide are not primarily related to the corrin ring system but with the generation and attachment of the axial ligands. Some of these reactions are also involved in corrin ring salvage.

Both, cobalamin and heme biosynthesis pathways have been reconstructed during manual curation of haloarchaeal genomes (Pfeiffer and Oesterhelt, 2015). Cobalamin biosynthesis has only few pathway gaps remaining. A proteomic study of *Hfx. volcanii* found a lower level of de novo cobalamin biosynthesis enzymes at increased temperature (Jevtic et al., 2019). For heme biosynthesis, some pathway details are yet unresolved.

**S3.a Currently, all *cbiX* gene paralogs are annotated as cobalt chelatase for cobalamin biosynthesis, but one may be an iron chelatase for heme biosynthesis**

Haloarchaeal genomes code for homologs of characterized early cobaltochelatases (CbiX-type, e.g. from *Bacillus megaterium* (Raux et al., 2003), *Methanothermobacter thermoautotrophicus* (Brindley et al., 2003), or *Archaeoglobus fulgidus* (Yin et al., 2006)). CbiX-type chelatases may be composed of a single or a pair of CbiX domains. Most haloarchaeal genomes code for two paralogous CbiX-type chelatases. Some species (e.g. *Nmn. pharaonis*, *Har. marismortui* and *Hbt. salinarum*) code for an additional CbiX domain protein.

*Nab. magadii* lacks genes for de novo cobalamin biosynthesis, so it was unexpected to find an early cobaltochelatase in this species (Siddaramappa et al., 2012). The *Natrialba* protein (Nmag_3212) is a close ortholog of HVO_1128. Thus, it can be speculated that HVO_1128 and its close orthologs from other halophilic archaea do not function in cobalamin biosynthesis. It can be predicted that they function in siroheme biosynthesis as sirohydrochlorin ferrochelatase (EC 4.99.1.4). This prediction awaits experimental confirmation.

It should be noted that the products of the *cbiK* gene in *Salmonella* (Raux et al., 1997) and *Desulfovibrio* (Lobo et al., 2008) were found to function as both cobaltochelatase in cobalamin biosynthesis and ferrochelatase in siroheme biosynthesis.

An alternative enzyme for sirohydrochlorin ferrochelatase activity is HVO_2312, which functions as precorrin-2 dehydrogenase but additionally may have sirohydrochlorin ferrochelatase activity. *E. coli* and *Salmonella* CysG is a trifunctional enzyme consisting of two domains. The N-terminal domain is bifunctional having precorrin-2 dehydrogenase as well as sirohydrochlorin ferrochelatase activities (Spencer et al., 1993, Stroupe et al., 2003, Pennington et al., 2020). The homologous *B. megaterium* SirC is, however, a monofunctional precorrin-2 dehydrogenase devoid of chelatase activity, the latter having been attributed to SirB (Raux et al., 2003). In the case of *Methanosarcina barkeri* (Mbar_A1461), precorrin-2 dehydrogenase was confirmed but sirohydrochlorin ferrochelatase activity was not analyzed (Storbeck et al., 2010). Only experimental analysis will allow to resolve whether HVO_2312 has sirohydrochlorin ferrochelatase activity.

**S3.b Two consecutive pathway gaps in cobalamin biosynthesis and four genes with unassigned function in the cobalamin gene cluster**

The cobalamin biosynthesis pathway has been reconstructed during manual curation of haloarchaeal genomes (Pfeiffer and Oesterhelt, 2015). As this reconstruction has not been described elsewhere, we list the associated genes and the underlying GSPs in Table 3, genes being ordered along the biosynthetic pathway. It is evident that most genes coding for the enzymes of the first part of the de novo cobalamin biosynthesis (upstream of cobyrinate a,c diamide) are clustered in the genome. In *Hfx. volcanii*, they are encoded on megaplasmid pHV3.

After detailed pathway reconstruction, two gaps remained open (EC 2.1.1.195 and 1.3.1.106). These enzymes catalyze consecutive steps which together result in conversion of cobalt-precorrin-5B to cobalt-precorrin-6B with cobalt-precorrin-6A being the intermediate. The first is a methyltransferase step, acting on C-1, while the second is a reduction, affecting the double bond between C18 and C19. It is quite possible that the order is inverted in haloarchaea, which would result in a novel intermediate. The cobalamin biosynthesis cluster contains four genes with yet unassigned function: HVO_B0052 (PQQ repeat protein), HVO_B0053 (DUF3209 domain protein), HVO_B0055 (conserved hypothetical) and HVO_B0056 (probable 4Fe-4S ferredoxin). A proteomic study found a decreased level of HVO_B0052 and HVO_B0055 at increased temperature, as found for the other enzymes of cobalamin biosynthesis (Jevtic et al., 2019).

SynTax analysis was performed in order to analyze if these genes are typical members of the cobalamin biosynthesis cluster. This cluster is plasmid-encoded (pHV3) in *Hfx. volcanii* and SyntTax only considers the main chromosome. Nevertheless, this analysis was possible as *Haloferax alexandrinus* strain wsp1 codes for an equivalent cluster which is chromosomally encoded, codes for highly conserved proteins (90-99% protein sequence identity) and gene order is identical (analyzed from *cbiG* (HVO_B0059, G3A49_06890) to *cbiE* (HVO_B0048, G3A49_06945). This region covers the four genes with yet unassigned function. SyntTax analysis uncovered that the four genes are strictly coupled to the cbiTLFGH1H2 operon in the majority of haloarchaeal genomes.

As this strengthens their status as candidates for closing the pathway gaps, we analyzed if their characteristics (e.g. domain architecture) would be compatible with the enzymatic conversions of the two pathway gaps. One missing enzyme is an oxidoreductase. It is obvious that a 4Fe-4S ferredoxin (HVO_B0056) may be involved in this reaction. HVO_B0052 is annotated as PQQ repeat protein. Such proteins probably use pyrroloquinoline quinone (PQQ) as prosthetic group, as e.g. found in quinoprotein alcohol dehydrogenases, making HVO_B0052 another top candidate for the missing reaction. The other two proteins (HVO_B0053, having a domain of uncharacterized function, DUF3209 and HVO_B0055, uncharacterized and devoid of any domain assignment), did not reveal any functional insights.

The other pathway gap is one of the many methyltransferases which are involved in cobalamin biosynthesis. In this context it should be noted that currently two paralogous enzymes are annotated to be responsible for C17 methylation: HVO_B0058, encoded by *cbiH1*, and HVO_B0057, encoded by *cbiH2*. These paralogs have 38% protein sequence identity to each other, with CbiH2 being much closer to characterized homologs. It may well be that CbiH1 is not responsible for methylation of C17 but of C1.

**S3.c Cobalamin biosynthesis and salvage reactions beyond cobyrinate a,c diamide**

Beyond the ligand cobyrinate a,c diamide are reactions which decorate the corrin ring (adenosylation, aminopropanol arm attachment, assembly of the nucleotide loop). The involved enzymes with their GSPs are listed in Table 3 (along the biosynthetic pathway with enzymes that prepare the ligands for attachment following after those directly acting on the corrin ring). For several of the enzymes, only a distant homolog has been characterized. Four enzymatic reactions need to be considered in more detail.

Two of these reactions are involved in the assembly of the nucleotide loop. In *Salmonella*, nicotinate-nucleotide-dimethylbenzimidazole phosphoribosyltransferase (encoded by *cobT*, EC 2.4.2.21) has been characterized (Trzebiatowski et al., 1994). Upon detailed bioinformatic analysis, this function has been assigned in *Halobacterium* (Rodionov et al., 2003). Based on the adjacent genes shown in Table 1 of that paper, this assignment is for VNG_1572C, which shows 57% protein sequence identity to its *Haloferax* ortholog HVO_0590. This function assignment is circumstantial and has not yet been confirmed experimentally. It should be noted that *Hht. litchfieldiae* has a much closer homolog to *Salmonella* CobT with 39% protein sequence identity (halTADL_3045, now known as SAMN05444271_10280). *Halohasta* lacks an ortholog of HVO_0590.

The subsequent reaction is catalyzed by adenosylcobalamin/alpha-ribazole phosphatase (encoded by CobC, EC 3.1.3.73), as shown in *Salmonella* (O'Toole et al., 1994). No homolog had been identified in *Halobacterium* (Rodionov et al., 2003). This was attributed to non-orthologous displacement and a candidate gene from the *Halobacterium* cobalamin cluster was proposed for this function (labelled HSL01294, Table 5 of that paper) (Rodionov et al., 2003). Based on the adjacent genes shown in Table 1 of that paper, this assignment is for VNG_1577C, which shows 56% protein sequence identity to its *Haloferax* ortholog HVO_0586. This function assignment has to be considered tentative and awaits experimental confirmation. It should be noted that *Salmonella* CobC and CobS show reduced substrate specificity which allows them to function in arbitrary order. If CobC acts first, alpha-ribazole phosphatase is hydrolyzed to alpha-ribazole and then attached by CobS to the corrin ring. If CobS acts first, alpha-ribazole phosphate is attached to the corrin ring, resulting in adenosylcobalamin phosphate which is then hydrolyzed to adenosylcobalamin by CobC.

The other two reactions that need to be considered in more detail are involved in attachment of the aminopropanol arm. In *Salmonella*, L-threonine-O-3-phosphate decarboxylase (encoded by *cobD*, EC 4.1.1.81) generates the intermediate 1-aminopropan-2-yl phosphate, a substrate for CbiB (Brushaber et al., 1998). For this enzyme, two paralogs with 51% protein sequence identity are encoded in close genomic vicinity in *Hfx. volcanii* (HVO_0591, cobD1, HVO_0593, cobD2). HVO_0591 shows 31% protein sequence identity to the characterized homolog from *Salmonella*. For HVO_0593, the assignment has to be based on its closely related paralog. A direct assignment based on a GSP is not possible, because BLASTp against SwissProt (accessed March 2021) returns histidinol-phosphate aminotransferases as top matches, but with less than 30% protein sequence identity. CobD decarboxylates threonine-O-3-phosphate, so that a threonine kinase is required. However, there is yet no candidate for this reaction (EC 2.7.1.177). Thus, either this reaction is catalyzed by a novel protein family or the corresponding pathway does not involve L-threonine-O-3-phosphate and CobD1 and CobD2 catalyze a distinct (but probably related) reaction. Closing of this pathway gap has to await experimental analysis.

**S3.d A potential late cobaltochelatase of the heterotrimeric type**

Cobalt is inserted late in the aerobic biosynthesis pathway. A late cobaltochelatase has been characterized (Debussche et al., 1992). It is heterotrimeric (with genes assigned as *cobNST*) and was stated to originate from *Pseudomonas denitrificans* (reassigned to *Sinorhizobium sp.* by UniProt as of April 2021). Haloarchaea code for a homolog of CobN (e.g. *Hfx. volcanii* HVO_B0050). No homologs to the other two subunits (UniProt:P29929, subunit CobS, UniProt:P29934, subunit CobT) could be identified in haloarchaea.

*Pseudomonas* CobN is distantly related to subunit ChlH from the characterized heterotrimeric magnesium chelatase from *Synechocystis sp.* PCC 6803 (Jensen et al., 1996, Jensen et al., 1998). The other subunits of the magnesium chelatase (ChlD and ChlI) are related to HVO_B0051. The N-terminal part of HVO_B0051 shows 46% protein sequence identity to ChlI and ca 30% protein sequence identity to the N-terminal part of ChlD. The remainder of HVO_B0051 shows ca 30% protein sequence identity to the remainder of ChlD. Comparative genomics analyses led to the observation that in several phylogenetic branches of bacteria the late cobaltochelatase has ChlID subunits instead of CobST subunits (Rodionov et al., 2003).

In summary, HVO_B0050 and HVO_B0051, which are encoded by adjacent genes within the major cobalamin cluster, might represent a late cobalt chelatase.

It may be unexpected to find a late cobalt chelatase in an organism that uses the cobalt-early biosynthetic pathway. However, a late cobalt chelatase may have a function in a corrin ring salvage pathway, a speculation not yet substantiated by experiment.

**S3.e Two siroheme decarboxylase paralogs, each being a two-domain protein**

The enzymes of the alternative heme biosynthesis pathway and their associated GSPs are listed in Table 3 (starting from siroheme and along the biosynthetic pathway; enzymes between precorrin-2 and siroheme are already covered above, see S3.a, *cbiX* gene paralogs). With respect to the conversion of siroheme to 12,18-didecarboxysiroheme, this function has been attributed to two haloarchaeal proteins (encoded by *ahbA* and *ahbB*). Both are two-domain proteins, while several characterized homologs are single-domain proteins forming heterodimers. These catalyze both decarboxylations  (Palmer et al., 2014). *Hydrogenobacter thermophilus* contains a single two-domain protein which has been characterized (Haufschildt et al., 2014). It is unclear if haloarchaeal AhbA and AhbB act independently (each forming a domain dimer) or as heterodimer (forming a domain tetramer).

#### Section S4: Coenzymes II: coenzyme F420

##### S4.a Coenzyme F420 biosynthesis in haloarchaea

Coenzyme F420 was first detected in *Methanobacterium* (Cheeseman et al., 1972) and is predominant in methanogenic archaea (Eirich et al., 1979, Jaenchen et al., 1984) but occurs also in certain bacteria and in small amounts in haloarchaea (Lin and White, 1986, Greening et al., 2016).

The pathway which creates the carbon backbone of coenzyme F420 has been reconstructed, based on the pathway characterized in *M. jannaschii*, and we list the enzymes with their associated GSPs in Table 4. Coenzyme F420 contains a phospholactate moiety which was reported to originate from 2-phospho-lactate (Grochowski et al., 2008). This compound is not well connected to the remainder of metabolism, and the assumed enzymes for its production could never be identified. The archaeal enzyme CofC (e.g. MJ0887) is involved in activation of 2-phospho-lactate and the enzyme CofD (e.g. MJ1256) in its attachment to the preassembled intermediate $F_0$ (Grochowski et al., 2008, Graupner et al., 2002). *Mycobacterium tuberculosis*, which also contains coenzyme F420, was assumed to also use this pathway. It turned out that the precursor for the phospholactate moiety was phosphoenolpyruvate, a common metabolic intermediate, rather than the atypical 2-phospho-lactate (Bashiri et al., 2019). Phosphoenolpyruvate was activated (by FbiD, the homolog of CofC) and coupled (by FbiA, the homolog of CofD), leading to a new intermediate, dehydro-F420. The authors confirmed usage of phosphoenolpyruvate and generation of the novel intermediate also for the enzymes from *M. jannaschii* (Bashiri et al., 2019). As a consequence, an additional enzyme activity is required, which reduces dehydro-F420 to F420. The authors found this activity in FbiB, a two-domain protein. The N-terminal part is homologous of CofE (MJ0768), the enzyme which attaches the polyglutamate tail to F420. The bacterial FbiB has a C-terminal extra domain which was known to contain a bound FMN for enigmatic reasons. It could be shown that this C-terminal domain reduces dehydro-F420 to F420, and that this reduction is a prerequisite for the N-terminal domain to attach the polyglutamate tail. Because the *Methanocaldococcus* CofE lacks such a domain, while CofC and CofD generate dehydro-F420, the authors tried to identify a genomically coupled dehydrogenase, but without success. The next twist to this story came with the analysis of coenzyme F420 biosynthesis in *Paraburkholderia rhizoxinica* (Braga et al., 2019). This species uses 3-phospho-D-glycerate instead of phosphoenolpyruvate or 2-phospho-lactate and generates a F420 derivative which has a phosphoglyceryl moiety instead of a phospholactyl moiety (Braga et al., 2019). These authors also determined the substrate specificity of the *M. jannaschii* enzymes CofC and CofD. In their hands, the enzymes work with the originally proposed precursor 2-phospho-lactate rather than the subsequently proposed precursor phosphoenolpyruvate. That would, again, resolve the problem with the missing dehydrogenase which converts dehydro-F420 to F420, but the problem that no biosynthetic route is known which leads to 2-phospho-lactate remains. It might thus be informative to subject the *Haloferax* coenzyme F420 biosynthetic pathway to experimental analysis.

**S4.b Putative coenzyme F420-dependent oxidoreductases**

To the best of our knowledge, only a single coenzyme F420 dependent enzymatic reaction has yet been reported for halophilic archaea (de Wit and Eker, 1987). No data were provided which would allow a gene assignment for this function. Thus, the importance of this coenzyme in haloarchaeal biology is highly enigmatic.

However, based on detailed bioinformatic analyses using a strategy called SIMBAL, coenzyme F420 dependent reactions have been predicted for bacteria (Selengut and Haft, 2010). Based on corresponding patterns and domains, F420 dependent enzymes were also predicted for haloarchaea.

HVO_0433 is a F420H2:NADP oxidoreductase, based on a characterized homolog from *A. fulgidus* (Kunow et al., 1993).

HVO_B0113 is annotated as a probable F420-dependent oxidoreductase, which is supported by the assignment of a corresponding InterPro domain (IPR019952). Proteins with this domain assignment belong to a coenzyme F420-specific subset of proteins with a luciferase-like monooxygenase domain. HVO_B0113 is located within the rhamnose cluster but is not required for growth on rhamnose (Reinhardt et al., 2019).

HVO_B0342 shows 29% protein sequence identity to a F420-dependent alcohol dehydrogenase from *Methanoculleus thermophilicus* which has been characterized, including a 3D structure with coenzyme F420 included as ligand (Klein et al., 1996, Aufhammer et al., 2004).

HVO_1937 is a probable 5,10-methylenetetrahydrofolate reductase, an enzyme of one-carbon metabolism (see below, one-carbon metabolism). It is briefly listed here because it has coenzyme F420-dependent methanogenic homologs. It should be noted that HVO_1937 is encoded adjacent to the gene coding for one of the coenzyme biosynthesis proteins (CofE).

The *Nmn. pharaonis* protein NP_1902A is name-giving to the InterPro version of TIGR04024 (F420_NP1902A, IPR023909). TIGRFAM entries for coenzyme F420-dependent enzymes were generated during a bioinformatic study of F420-dependent enzymes in *M. tuberculosis*, applying the SIMBAL algorithm (Selengut and Haft, 2010). NP_1902A or any of its close homologs has not been experimentally characterized, so that usage of F420 has not yet been confirmed. There is a paralog in *Nmn. pharaonis* (NP_4374A, 44% protein sequence identity) and there are more closely related orthologs (at least 60% protein sequence identity) in several haloarchaea (e.g. *Har. marismortui*, *Hbt. salinarum*, *Nab. magadii*). A close ortholog is absent from *Hfx. volcanii*, which only has distant homologs (e.g. HVO_A0572, 33% protein sequence identity, and no assignment of the F420_NP1902A domain by InterPro).

The *Nmn. pharaonis* protein NP_4006A is distantly related to Coenzyme F420-dependent sulfite reductase from *M. jannaschii* (Johnson and Mukhopadhyay, 2005) and has the corresponding InterPro domain assigned (IPR007516).

In addition, there are proteins annotated as F420-dependent NADP oxidoreductase family protein (e.g. HVO_A0605). They belong to the same protein family as F420H2:NADP oxidoreductase, but only a subset uses coenzyme F420 while the others use NAD. Only experimental analysis can clarify the coenzyme specificity.

##### S4.c The coenzyme F420 precursor may be involved in DNA repair

UV light induces two types of DNA lesions in halophilic archaea: cyclobutane pyrimidine dimers and pyrimidine (6-4) pyrimidone photoproducts (for reviews on DNA repair see (Jones and Baxter, 2017, Perez-Arnaiz et al., 2020)). Photoreactivation of both lesion types has been detected, in addition to dark repair (McCready, 1996). The action spectrum of photoreactivation has been analyzed in vivo and in cell extracts of *Hbt. salinarum*. The process depends on 8-hydroxy-5-deazaflavin (F0, the precursor of F420 lacking the poly-Glu tail) (Eker et al., 1991, Iwasa et al., 1988). This is, however, an assignment to an overall process and not to individual proteins.

The enzyme which repairs cyclobutane pyrimidine dimers has been characterized in *Hbt. salinarum* (VNG_1335G, encoded by *phr2*) (Takao et al., 1989, McCready and Marcello, 2003). However, the chromophore has not been analyzed. Phr2 is one key player for the remarkable UV-radiation resistance of *Hbt. salinarum* (Baliga et al., 2004). The ortholog from *Hfx. volcanii* is HVO_2911.

*Hfx. volcanii* codes for two other potential photo-lyases (HVO_2843, encoded by *phr1*, HVO_1234, encoded by *phr3*). HVO_2843 belongs to the cryptochrome DASH class which was long considered not to be related to DNA repair (Brudler et al., 2003, Kleine et al., 2003). However, it turned out that this class of proteins mediates photorepair of cyclobutane pyrimidine dimers in single-stranded DNA (Selby and Sancar, 2006). HVO_1234 may function as pyrimidine (6-4) pyrimidone photolyase (Zhang et al., 2013).

#### Section S5: Coenzymes III: coenzymes of C1 metabolism: tetrahydrofolate in haloarchaea, methanopterin in methanogens

Several coenzymes are prominently associated with methanogenic organisms (methanopterin, coenzyme F420, coenzyme M, coenzyme B, cofactor F430). For two of these (methanopterin, coenzyme F420), corresponding annotations are found for halophilic archaea in public databases. As detailed above, this is valid for coenzyme F420. However, methanopterin, which is the one-carbon carrier in methanogenic archaea, does not exist in halophilic archaea, which use tetrahydrofolate as one-carbon carrier (White, 1988). The distinct one-carbon carriers share a near-identical core structure (a pterin heterocyclic ring is linked via a methylene bridge to a phenyl ring) and also share a polyglutamate tail. The sequence similarity between methanopterin-specific methanogenic enzymes and tetrahydrofolate-specific haloarchaeal enzymes indicates divergence from a common ancestor. Experimentally characterized methanogenic enzymes thus may trigger annotation of haloarchaeal enzymes as being methanopterin-specific, which as a consequence implies the presence of methanopterin in haloarchaea. Such annotations are found in UniProt and KEGG (accessed April 2021). However, when haloarchaea switched from methanopterin to tetrahydrofolate as a C1 carrier, the haloarchaeal enzymes had to adapt to tetrahydrofolate, which shows structural similarity to methanopterin. Additionally, some details of haloarchaeal tetrahydrofolate biosynthesis are yet unresolved. A detailed review on the many variants of the tetrahydrofolate biosynthetic pathway is available (de Crecy-Lagard, 2014). Also, tetrahydrofolate and tetrahydromethanopterin have been compared (Maden, 2000).

**S5.a A proposed pathway for aminobenzoate biosynthesis in haloarchaea**

Tetrahydrofolate biosynthesis requires aminobenzoate. No haloarchaeal enzymes for the biosynthesis of aminobenzoate are assigned in KEGG (accessed April 2021). We had proposed candidates for this pathway (Falb et al., 2008, Pfeiffer et al., 2008) which would convert chorismate to para-aminobenzoate. Our proposal is based on clustering of three genes, the products of which are homologous to enyzmes involved in amino acid biosynthesis. The components of a proposed aminodeoxychorismate synthase (HVO_0709 and HVO_0710, encoded by *pabAB*) are homologous to those of anthranilate synthase (encoded by *trpEG*). A proposed aminodeoxychorismate lyase (encoded by *pabC*) is distantly related to the product of *ilvE* (branched-chain amino acid aminotransferase). According to SyntTax analysis, *pabAB* are strictly clustered and *pabC* is genomically coupled in the majority of haloarchaeal genomes. This prediction is not supported by sufficient evidence for databases to adopt this. Also, the detailed review on tetrahydrofolate biosynthesis states that “in Archaea, the pABA synthesis pathway remains a mystery” (de Crecy-Lagard, 2014).Thus, this prediction is awaiting experimental support or disproval.

**S5.b GTP cyclohydrolase MptA is probably involved in biosynthesis of tetrahydrofolate rather than methanopterin**

The annotation of HVO_2348 (GTP cyclohydrolase MptA, encoded by *mptA*, with gene synonym *folE2*) is based on the characterized homolog from *M. jannaschii* (Grochowski et al., 2007). In UniProt, the protein from this methanogenic archaeon, is stated to "convert GTP to 7,8-dihydro-D-neopterin 2',3'-cyclic phosphate, the first intermediate in the biosynthesis of coenzyme methanopterin". Unfortunately, this statement is copied to HVO_2348 (as of March 2021), which is invalid, given that haloarchaea do not generate/contain methanopterin. The initial reactions of methanopterin and of tetrahydrofolate biosynthesis are common (up to the intermediate (7,8-dihydropterin-6-yl)methyl diphosphate). However, the enzymes specific for methanopterin biosynthesis are absent from haloarchaea. It should be noted that HVO_2348 has been partially characterized (El Yacoubi et al., 2009). Deletion mutants of HVO_2348 are auxotrophic for thymidine and hypoxanthine. The conversion of dUMP to dTMP is dependent on dihydrofolate. Two enzymatic reactions of de novo purine biosynthesis (EC 2.1.2.2 and 2.1.2.3) are also dependent on tetrahydrofolate, and this block can be overcome by purine salvage, using hypoxanthine.

The product of HVO_2348 is the substrate for EC 3.1.4.56. This enzyme has been characterized from *M. jannaschii* (MJ0837) but remains a pathway gap in halophilic archaea. MJ0837 is very distantly related to HVO_A0533 with only 27% protein sequence identity. Nevertheless, HVO_A0533 might be a promising candidate and a deletion strain could be checked for thymidine and hypoxanthine auxotrophy.

HVO_2628 is a member of the GHMP kinase family (kinases for galactose, homoserine, mevalonate and phosphomevalonate). There is about 30% protein sequence identity to beta-ribofuranosylaminobenzene 5'-phosphate synthase (EC 2.4.2.54) and this function is also assigned to HVO_2628 by UniProt and by KEGG (as of March 2021). This is based on the enzymatically characterized and distantly related enzymes from *A. fulgidus* (Scott and Rasche, 2002) and *M. jannaschii* (Dumitru and Ragsdale, 2004). The reaction of EC 2.4.2.54 is not a simple kinase reaction, and it represents the first committed step to methanopterin biosynthesis.

As this coenzyme is absent from halophilic archaea, a distinct function is likely. However, no corresponding reaction is required in the tetrahydrofolate biosynthesis pathway. The true biological function of HVO_2628 remains unresolved.

**S5.c Enzymes which alter the oxidation level of the coenzyme-attached one-carbon compound likely operate on tetrahydrofolate rather than methanopterin**

HVO_2573 (*mch*, probable methenyltetrahydrofolate cyclohydrolase; EC 3.5.4.9) is annotated in UniProt and KEGG as "methenyltetrahydromethanopterin cyclohydrolase (EC 3.5.4.27)" (as of March 2021). The characterized homolog is MK0625 from the methanogenic archaeon *M. kandleri* (Vaupel et al., 1998). These proteins show 45% protein sequence identity. The C1 compound is attached to the substructure shared between tetrahydrofolate and methanopterin, and it is quite possible that these homologous proteins catalyze an equivalent reaction, but using distinct coenzymes. An experimental characterization of the haloarchaeal protein would unequivocally resolve this annotation ambiguity.

HVO_1937 (*mer*, probable 5,10-methylenetetrahydrofolate reductase) is annotated in UniProt and KEGG as "5,10-methylenetetrahydromethanopterin reductase (EC 1.5.98.2)" (as of March 2021). This annotation is based on the characterized *mer* from the methanogenic archaeon *M. thermoautotrophicus* (Shima et al., 2000, Vaupel and Thauer, 1995, te Brömmelstroet et al., 1990). These proteins show 38% protein sequence identity. As stated for HVO_2573, the usage of distinct but structurally related coenzymes is well possible at this level of sequence conservation. As mentioned above (see section S4.b), the methanogenic enzyme functions with coenzyme F420 and this is also likely for the halophilic protein. Again, experimental characterization would resolve these issues.

#### Section S6: Coenzymes IV: NAD and FAD (riboflavin)

**S6.a Two NAD kinase paralogs may function with ATP or polyphosphate**

The paralogs *nadK1* (HVO_2363) and *nadK2* (HVO_0837) are only distantly related to each other. HVO_2363 is more closely related to NAD kinase from *M. tuberculosis* than to NAD kinase from *A. fulgidus*. The *Mycobacterium* enzyme can use both, ATP and pyrophosphate as energy source (Kawai et al., 2000). This information seems unavailable for the enzyme from *A.*

*fulgidus*, even though the protein has been crystallized together with NAD, NADP, and ATP (Liu et al., 2005). After soaking with ATP, the structure contains two ATP molecules. One is well structured but occupies part of the NAD binding site, which thus may represent fortuitous binding in the absence of NAD. For the other, which is considered to represent the phosphate donor, only the beta/gamma pyrophosphate is ordered enough to be visible. This might imply that this NAD kinase may function with both, ATP and polyphosphate. HVO_0837 is even more distantly related to characterized homologs. Thus, the assignment of the energy source of the *Haloferax* enzymes is not possible in the absence of experimental data. It should, however, be noted that polyphosphate could not be identified in exponentially growing *Hfx. volcanii* cells by DAPI staining (Zerulla et al., 2014), so that ATP is the more likely energy source.

**S6.b HVO_0781 may be involved in NAD biosynthesis due to a highly conserved gene neighbourhood with HVO_0782 (*nadM*)**

HVO_0782 is annotated as nicotinamide-nucleotide adenylyltransferase (*nadM*, EC 2.7.7.1), a step in NAD biosynthesis, based on the characterized homolog from *M. jannaschii* (Raffaelli et al., 1997, Raffaelli et al., 1999). This enzyme is present in all Halobacteria (163 species) and even in 399 of 405 archaea (according to OrthoDB analysis, March 2021). The gene neighbourhood to its adjacent gene, HVO_0781, is conserved in most haloarchaea and beyond according to SyntTax analysis. HVO_0781 occurs in nearly all halophilic archaea (158 of 166) and in 296 out of 405 archaea, indicating that it must have an important function. It is related to the characterized S-adenosyl-L-methionine hydrolase (adenosine-forming) from *Salinispora arenicola* and from *Pyrococcus horikoshii* (Eustaquio et al., 2008, Deng et al., 2009). S-adenosyl-L-methionine hydrolase has the function to destroy S-adenosylmethionine, which seems a wasteful reaction, and not expected to have been very well conserved over evolutionary time. Currently, HVO_0781 is annotated as "S-adenosylmethionine hydroxide adenosyltransferase family protein" to avoid the firm assignment of a function that is considered metabolically useless. The very strongly conserved gene neighbourhood with the NAD biosynthesis enzyme NadM may hint to an involvement in NAD biosynthesis, which could be investigated by generating mutants and checking for potential phenotypes.

**S6.c Candidate genes which fill various gaps in riboflavin biosynthesis**

Archaeal organisms were included in a 2017 study that provided an extensive bioinformatic analysis of riboflavin biosynthesis (Rodionova et al., 2017). Reconstruction of this pathway for haloarchaea, including enzymes and their GSPs, is provided in Table 6. In riboflavin biosynthesis, D-ribulose-5-phosphate is converted by RibB (HVO_0327) to a compound termed "VI" (Rodionova et al., 2017). This is condensed with another compound (termed "V'") by RibH (HVO_0974, 6,7-dimethyl-8-ribityllumazine synthase, EC 2.5.1.78). Compound "V'" is generated by the successive actions of ArfA, ArfB, ArfC, a gene product referred to as PyrD, and a putative phosphatase (Rodionova et al., 2017). Only two of these enzymes have sufficient support to be annotated (ArfA, HVO_1284, EC 3.5.4.29, ArfC, HVO_1341, EC 1.1.1.302). For ArfB (EC 3.5.1.102), HVO_1235 has been proposed as a candidate (Rodionova et al., 2017). For the so-called PyrD (RIB2 in KEGG as of April 2021, similar to but distinct from EC 3.5.4.26), HVO_2483 was proposed as a candidate (Rodionova et al., 2017). If confirmed, a distinct gene name would have to be assigned (*pyrD* is already assigned to dihydroorotate dehydrogenase). No proposal has been made for the phosphatase which has now been assigned in various species (according to the information associated with EC 3.1.3.104) (Haase et al., 2013, London et al., 2015, Sarge et al., 2015) but without enabling a prediction for haloarchaea.

The conversion of riboflavin to FMN is catalyzed by a bifunctional CTP-dependent riboflavin kinase / transcription regulator (HVO_0326, *rbkR*) which contains a N-terminal HTH domain and a C-terminal catalytic domain. FMN is then converted to FAD by HVO_1015 (encoded by *ribL*). The enzymes proposed to close the pathway gaps (ArfB and pyrimidine deaminase) require experimental confirmation.

#### Section S7: Biosynthesis of membrane lipids, bacterioruberin and menaquinone

##### S7.a Geranylgeranyl diphosphate synthase and geranylfarnesyl diphosphate synthase

The main core lipid of haloarchaea is archaeol, which consists of glycerol with ether-linked C20 isoprenoid units, which are derived from geranylgeranyl diphosphate. In *Hfx. volcanii*, HVO_2725 (*idsA1*) and HVO_0303 (*idsA2*) are annotated as geranylgeranyl diphosphate synthase.

Besides the main core lipid archaeol, there are variants, some of which have a C25 isoprenoid rather than a C20 isoprenoids (see e.g. (De and Gambacorta, 1988, Dawson et al., 2012). These longer isoprenoids commonly occur in alkaliphilic organisms like *Nmn. pharaonis* and *Nab. magadii* (De Rosa et al., 1982). An enzyme involved in C25 isoprenoid biosynthesis (geranylfarnesyl diphosphate synthase) has been purified and characterized on the enzymatic level in *Nmn. pharaonis* (Tachibana, 1994). However, data allowing gene assignment were not provided in that study, nor has it been cited in any later publication reporting gene assignment. Three candidate genes exist in *Nmn. pharaonis*, (NP_0604A, NP_3696A, and NP_4556A). As NP_0604A, NP_3696A have orthologs in *Hfx. volcanii*, an organism devoid of C25 lipids, and as orthologs of these are found in nearly all genomes from Halobacteria according to OrthoDB analysis (159/162 of 166), we assign geranylfarnesyl diphosphate synthase activity to the third paralog, NP_4556A. This protein has a close homolog in comparably few halophilic archaea according to OrthoDB (64 of 166), among them *Nab. magadii*, which is also alkaliphilic. Uniprot added the Tachibana 1994 reference (Tachibana, 1994) to the entry for NP_3696A and assigned geranylfarnesyl diphosphate synthase activity to this paralog (as of April 2021) but without providing any evidence for this assignment. Geranylfarnesyl diphosphate synthase has been characterized in *A. pernix* (*fgs*, APE_1764) (Tachibana et al., 2000), but for this class of enzymes general sequence similarity is insufficient to predict substrate specificity. Our assignments are supported by analysis of key residues which determine the length of the isoprenoid chain (Bale et al., 2019). These authors label the cluster containing NP_3696A (WP011323557.1) as "C15/C20" and the cluster containing NP_4556A (WP011323984.1) as "C20->C25->C30?". Unambiguous specificity assignments have to await experimental characterization.

**S7.b Biosynthesis of the phospholipid headgroups is only partially resolved**

The haloarchaeal core lipid, archaeol, is activated to CDP-archaeol by the product of the *carS* gene (HVO_0332). Distinct headgroups are then attached by CDP-archaeol 1-archaetidyltransferases which share a common domain (represented e.g. by InterPro:IPR000462). Four members of this protein family are encoded in the *Hfx. volcanii genome* (HVO_1136, HVO_1143, HVO_1297, HVO_1971). Even though these proteins share the same domain, they are so distant that they do not detect each other upon BLASTp analysis (stringency of $10^{-5}$).

HVO_1143 is distantly related (32% protein sequence identity) to a functionally characterized archaetidylserine synthase from *M. thermoautotrophicus* (MTH_1027) (Morii and Koga, 2003). The *Hbt. salinarum* ortholog (VNG_0784G) has been assigned the same function in a bioinformatic analysis (Daiyasu et al., 2005). HVO_1143 has recently been shown to be involved in ArtA-dependent protein maturation, a process which includes C-terminal proteolytic cleavage and lipid attachment (Abdul-Halim et al., 2020). It is referred to as PssA, based on the gene assigned to the bacterial phosphatidylserine synthase. HVO_0146, which probably decarboxylates archaetidylserine to archaetidylethanolamine, was also found to be involved in ArtA-dependent protein maturation (referred to as PssD) (Abdul-Halim et al., 2020). The same function was also proposed for the *Hbt. salinarum* ortholog (VNG_2255C) (Daiyasu et al., 2005).

The substrate assignment to HVO_1297 is uncertain. It shows distant similarity (25% protein sequence identity) to the characterized archaetidylinositol phosphate synthase from *M. thermoautotrophicus* (MTH_1691, EC 2.7.8.39) (Morii et al., 2009). The ortholog from *Nmn. pharaonis* (NP_2144A) is somewhat less distant, having 32% protein sequence identity. KEGG also assigns EC 2.7.8.39 (via KO group K17884, accessed April 2021). It should be noted that MTH_1691 does not retrieve any of the other CDP-archaeol 1-archaetidyltransferases from *Hfx. volcanii* upon BLASTp analysis (stringency of $10^{-5}$). Upon bioinformatic analysis, the *Hbt. salinarum* ortholog (VNG_1030G) had been assigned to function as archaetidylglycerol synthase (Daiyasu et al., 2005). This assignment is exclusively based on the reported absence of archaetidylinositol from haloarchaea (Daiyasu et al., 2005, Kates, 1977). In a subsequent review, the corresponding cluster is stated to be "related to myo-inositol-phosphate transfer" (Lombard et al., 2012) and the authors add "(and maybe also glycerol in archaea, according to classification in [70]", which refers to (Daiyasu et al., 2005).

Various issues need to be taken into account for the evaluation of this specific open issue. (a) Is archaetidylinositol really absent from haloarchaea, even as a pathway intermediate? In this context it should be noted that archaetidylethanolamine has long been considered absent from haloarchaea, but recently has been identified (see above). (b) It should be taken into account which fraction of the haloarchaeal genomes codes for an ortholog of the *Haloferax* proteins. According to OrthoDB, orthologs of HVO_1143 and HVO_1297 are encoded in nearly all of the genomes from *Halobacteria* (159/161 of 166), of which only about half (80 out of 166) code for an ortholog of HVO_1136 and only one-fifth (26 out of 166) code for an ortholog of HVO_1971. As PGP and PGP-Me are prominent phospholipids in most haloarchaea, this might imply that HVO_1297 is more likely involved in attachment of glycerol phosphate than myo-inositol phosphate. (c) HVO_1297 is part of a three-gene operon, the synteny of which is highly conserved according to SyntTax analysis. We thus evaluated if the adjacent genes might shed light on the function prediction for HVO_1297.

One gene of the cluster is annotated as histidinol-phosphate aminotransferase (HVO_1295, encoded by *hisC*). This function assignment is based on the capability of this gene to overcome a histidine auxotrophy mutation (Conover and Doolittle, 1990). The complementing gene was then found to be distantly related (31% protein sequence identity) to *E. coli* HisC. Substrate assignments are highly uncertain at this level of sequence identity in this protein family (see above, Section 2). It should be kept in mind that the complementation activity observed for HVO_1295 might be due to a side reaction of an enzyme with relaxed substrate specificity and that the major function of this pyridoxal-phosphate dependent aminotransferase might be involved in a polar lipid biosynthesis pathway.

The other gene of this highly conserved three-gene operon is annotated as adenylate kinase (HVO_1296, encoded by *adk2*). There is comparably low sequence identity to proteins with confirmed adenylate kinase activity. HVO_1296 shows 34% protein sequence identity to a *Pyrococcus* homolog, which seems to have its primary function in ribosome biogenesis (Loc'h et al., 2014). HVO_1296 shows 32% protein sequence identity to human AK6 which has broad substrate specificity and is more a nucleotide kinase rather than an adenylate kinase (Ren et al., 2005). It should be noted that the *adk1* gene product of *Haloferax* (HVO_2496) has a much stronger support as being adenylate kinase (45% protein sequence identity to a characterized homolog from *B. subtilis*). Thus, there is only weak support for HVO_1296 being an adenylate kinase and the protein might, instead, function as inositol kinase. In that case, the reaction product, myo-inositol-1-phosphate, would be a potential substrate for HVO_1297 if that enzyme would generate archaetidylinositol phosphate.

*Haloferax* codes for an additional gene that is annotated as myo-inositol-1-phosphate synthase, HVO_B0213. Assignment of this specificity is based on a characterized homolog from *A. fulgidus* (AF_1794). Only few haloarchaea code for a close homolog (32 of 166). If a majority of haloarchaea would produce archaetidylinositol, at least as a pathway intermediate, a more widely distributed enzyme would be required. In that case, HVO_1296 (as inositol kinase) and HVO_1297 (as archaetidylinositol phosphate synthase) might be responsible for these activities. In summary, the substrate of HVO_1297 remains uncertain and will stay an open issue until being resolved by experimental analysis.

According to OrthoDB analysis, the two residual members of the CDP-archaeol 1-archaetidyltransferase family are encoded in about half (80 out of 166 for HVO_1136) and about one-fifth (26 out of 166 for HVO_1971) of the haloarchaea. HVO_1136 does not show full-length similarity to a GSP, so that this approach does not allow a specific function assignment. Speculations regarding the function of HVO_1136 depend on the tentative function assignment to HVO_1297. If HVO_1297 were an archaetidylinositol phosphate synthase, then HVO_1136, having a wider representation in haloarchaea, would be a candidate for an archaetidylglycerophosphate synthase. The product of this enzyme is PGP, which is probably methylated to PGP-Me, a polar lipid found in a wide variety of haloarchaea. On the other hand, PGP is dephosphorylated to PG, which also is found in many haloarchaea. Support for this assignment might be the adjacent gene (HVO_1135), which is encoded on the opposite strand. Based on an InterPro domain assignment, it is probably a S-adenosylmethionine-dependent methyltransferase. HVO_1971, being comparably rare in haloarchaea, shows 26% protein sequence identity to the characterized archaetidylserine synthase from *M. thermoautotrophicus* (MTH_1027) which, however, is more closely related to HVO_1143. Thus, we think that no specific function can be assigned for HVO_1971. All proposed function assignments have to be considered uncertain until their experimental confirmation or rejection.

Haloarchaea are also known to contain cardiolipin and glycolipids (e.g. S-DGD-1, sulfated diglycosyldiether) (Sprott et al., 2003). The biosynthetic pathways leading to these polar lipids are currently enigmatic. Also enigmatic are the enzyme responsible for sulfation of the glycolipid, and the enzyme responsible for the sulfur transfer reaction that leads to PGS.

**S7.c Conversion of geranylgeranyl diphosphate to the carotenoid bacterioruberin**

The C20 compound geranylgeranyl diphosphate is converted to the C40 carotenoid phytoene by head-to-head condensation. This reaction is catalyzed by phytoene synthase (encoded by *crtB*, e.g. HVO_2524). Only distantly related homologs have been enzymatically characterized (Chamovitz et al., 1992). However, *crtB* mutants are colorless (Kiljunen et al., 2014, Maurer et al., 2018, Liu et al., 2011), which supports the involvement of this gene in carotenoid biosynthesis.

Next, phytoene is reduced to lycopene by phytoene desaturase. The bioinformatic assignment of phytoene desaturase activity to a gene proved to be difficult and will be covered after bacterioruberin biosynthesis.

Lycopene is converted to bacterioruberin by three enzymes which have been characterized experimentally in *Haloarcula japonica* (Yang et al., 2015). The genes for these enzymes form a three gene cluster which is syntenically highly conserved according to SyntTax analysis. Based on the strict syntenic conservation of this three gene cluster in haloarchaea, we hypothesize that all orthologs are isofunctional to the enzymes from *Har. japonica*. Lycopene elongase (encoded by *lyeJ*, c0506 in *Har. japonica*, HVO_2527 in *Hfx. volcanii*, NP_4766A in *Nmn. pharaonis*) adds a C5 isoprenoid unit to one end of lycopene, generating a C45 intermediate (Yang et al., 2015). LyeJ is bifunctional and also acts as a 1,2-hydratase. Next, carotenoid 3,4 desaturase (encoded by *crtD*, c0506, HVO_2528, NP_4764A) introduces a double bond in the newly attached isoprenoid unit. This pair of reactions is repeated on the other side of the molecule, thus resulting in a C50 compound. Addition of C5 isoprenoid units to one end of lycopene was first observed in *Hbt. salinarum* (Dummer et al., 2011, Yang et al., 2015). As last step in biosynthesis, a C50 carotenoid 2”-3”-hydratase (encoded by *cruF*, c0505, HVO_2526, NP_4768A) generates the final product bacterioruberin.

This leaves one pathway gap, phytoene desaturase. For *Nmn. pharaonis*, this reaction had initially been attributed to NP_4764A (encoded by *crtI1*, now known as *crtD*, see above) and NP_0204A, encoded by *crtI2* (Falb et al., 2008). It is uncertain and may even be considered unlikely that the *crtD* product (c0506, HVO_2528, NP_4764A) has broad substrate specificity and also has phytoene desaturase activity, thus acting on a C40 compound and on bonds which occur in the center of the molecule. Deletion of the *crtD* gene in *Har. japonica* (c0507) did not interfere with lycopene biosynthesis (Yang et al., 2015). In *Hfx. volcanii*, a deletion mutant of *crtD* was white in colour (Maurer et al., 2018), which might indicate that lycopene, a red-colored compound, cannot be produced. The *Nmn. pharaonis* paralog of *crtD*, NP_0204A, which has 60% protein sequence identity to NP_4764A, has comparably few orthologs in haloarchaea and thus may not represent a key enzyme of lycopene biosynthesis. Most haloarchaea have additional very distant paralogs of *crtD* (in *Nmn. pharaonis*/*Hfx. volcanii* NP_1630A/HVO_0817 and NP_4520A/HVO_2340). Despite being very distantly related, these are obvious candidates for being a phytoene desaturase. The *Hfx. mediterranei* ortholog of HVO_0817 (HFX_0786, HFX_RS03805) is depicted as a candidate for phytoene desaturase activity in the detailed bioinformatic reconstruction of haloarchaeal carotenogenesis (see Fig.3 of (Giani et al., 2020)), even though *Hfx. volcanii* HVO_0817, HVO_RS08620, is not depicted in that figure. It should be noted that phytoene desaturase catalyzes an oxidoreductase reaction, and the next gene upstream of the *crtD*-*lyeJ*-*cruF* three gene cluster codes for an oxidoreductase of the short-chain dehydrogenase family (HVO_2529, an ortholog of NP_4762A).

**S7.d Reconstruction of menaquinone biosynthesis**

Halophilic archaea contain high levels of menaquinone (mainly MK8:8) (Elling et al., 2016, Kellermann et al., 2016). The pathway for menaquinone biosynthesis has been reconstructed, but two pathway gaps remain open. The enzymes and their associated GSPs are listed in Table 7 (starting from chorismate and along the biosynthetic pathway). Many functions can be attributed by clear, though distant, sequence similarity (enzymes encoded by *menF*, *menD*, *menE, menB*, *menA*, *menG*). In one case, sequence similarity is very low but the assignment is supported by clustering of many genes coding for menaquinone biosynthesis enzymes (*menC*, HVO_1461). This assignment, and also that of menG has not been made by KEGG (accessed March 2021). Two pathway gaps remain (MenH, EC 4.2.99.20 and MenI, EC 3.1.2.28). There are five genes with unassigned function in the gene cluster coding for enzymes of the menaquinone biosynthesis pathway (HVO_1463,1464,1466,1467,1468), but we could not detect any obvious relationship between the domains assigned to their products and the missing enzymatic reactions.

#### Section S8: Issues concerning RNA polymerase, protein translation components and signal peptide degradation

##### S8.a RNA polymerase subunit epsilon (*rpoeps*)

Beyond the canonical haloarchaeal RNA polymerase subunits (encoded by *rpoA1A2B1B2DEFHKLNP*), *Hbt. salinarum* was found to contain an additional subunit called epsilon (Leffers et al., 1989, Madon and Zillig, 1983). This was required for efficient translation of native templates (Madon and Zillig, 1983). Even though the subunit is encoded in the genome of *Halococcus morrhuae* (C448_10607), it was not identified in the purified RNA polymerase from that species (Madon et al., 1983). While canonical RNA polymerase subunits are found in nearly all haloarchaea (e.g. 159 of 166 when OrthoDB is queried for subunit H), the epsilon subunit is only found in a subset of the Halobacteria (73 of 166). It occurs in most *Natrialbales* (47 of 49) and about half of the *Halobacteriales* (24 of 46) but is rare in *Haloferacales* (3 of 71). We consider it astounding that the core component of genetic information processing shows this level of variability within the haloarchaea. The biological relevance of this subunit seems largely unresolved.

##### S8.b Ribosomal proteins having two distant paralogs

Two distant paralogs are found for haloarchaeal ribosomal protein S10 in nearly all haloarchaeal genomes (in *Hfx. volcanii* HVO_0360, encoded by *rps10a*, and HVO_1392, encoded by *rps10b*, with 24% protein sequence identity). It is uncertain if both occur in the ribosome, together or alternatively, the latter would result in heterogeneity of the ribosomes. Alternatively, one of the paralogs may exclusively have a non-ribosomal function. According to SyntTax analysis, *rps10a* and *tef1a1* form a gene pair that is highly conserved and is found in nearly all haloarchaea. The gene *tef1a1* codes for translation elongation factor aEF-1 alpha / peptide chain release factor aRF-3. This gene clustering supports a ribosomal function of the *rps10a* gene product. The gene *rps10b* is monocistronic but the divergently transcribed upstream gene (HVO_1391, NUDIX family hydrolase) is adjacent in most haloarchaeal genomes.

In a subset of archaea, two distant paralogs are found for haloarchaeal ribosomal protein S14 (in *Nmn. pharaonis* NP_4882A, encoded by *rps14a*, and NP_1768A, encoded by *rps14b*). The C-terminal ca 30 aa show 82% protein sequence identity while the N-terminal ca 20 aa are unrelated. The gene *rps14a* is encoded within a huge operon encoding 22 ribosomal proteins. The gene *rps14b* is monocistronic and its gene neighbourhood is not conserved.

##### S8.c A highly conserved C2-C2 type zinc finger is present in ribosomal protein L43e from *Halobacteriales* but absent in those from *Haloferacales* and *Natrialbales*

A subset of archaeal ribosomal proteins has homologs in eukaryotes but not in bacteria. Among these is ribosomal protein L43e, the homolog of the eukaryotic protein initially named L37a (rat) and later L43 (yeast). This protein is known to contain a zinc finger, representing a rubredoxin-like metal-binding fold according to InterPro (IPR011331) (CxxCx$_n$CxxC). For the yeast protein it was found that each of the four cysteine residues is required for zinc binding but only one of them is essential for the function of the protein (Rivlin et al., 1999). Between the CxxC patterns is a conserved “Trp motif” (Rxxx[G/A/S][I/V/L]W) and mutation of the strictly conserved tryptophan residue is lethal (Dresios et al., 2002). Ribosomal protein L43e (L37a3) from *Har. marismortui* had not been sequenced prior to determination of the X-ray structure of the large ribosomal subunit at 2.4 A resolution (Ban et al., 2000) but the protein could be built into the structure, based on the homologous sequences from rat and yeast, which was facilitated by the bound zinc atom.

The strictly conserved residues of the zinc finger motif (CPxC and CxxCG) are reminiscent of those found in the so-called "small CPxCG-related zinc finger proteins" (Klein et al., 2007, Tarasov et al., 2008, Nagel et al., 2019).

While the zinc finger and the central "Trp motif" are conserved in most homologs from euryarchaeota, there are exceptions. Both motifs are conserved in all euryarchaeal proteins from taxonomic orders outside Halobacteria, while within Halobacteria this is only the case for proteins of the order *Halobacteriales*. None of the proteins from the taxonomic order *Haloferacales* and a minority of the proteins from the taxonomic order *Natrialbales* contain the zinc finger, while the Trp-motif is conserved. This surprising observation was first made by UniProt staff (and communicated by e-mail) when their stringent consistency checking routines, applied to members of HAMAP MF_00327, failed to detect the expected cysteine residues (Pedruzzi et al., 2015).

**Section S8.d Diphthamide, a posttranslational modification of translation elongation factor EF-2**

Three enzymatic reactions convert a histidine residue in translation elongation factor EF-2 into diphthamide. We list these three enzymes in Table 8. Two of these (encoded by *dph2* and *dph5*) are annotated according to characterized homologs from *P. horikoshii* (PH1105, PH0725) (Zhu et al., 2011, Zhu et al., 2010). The last enzymatic step (encoded by *dph6*) was assigned in a detailed bioinformatic analysis (de Crecy-Lagard et al., 2012). This assignment is consistent with distant similarity to the N-terminal region of a characterized homolog from yeast (YLR143W encoded by *DPH6*) (Su et al., 2012, Uthman et al., 2013).

**S8.e Signal peptide peptidase**

N-terminal signal sequences target proteins to the secretion machinery. Subsequent to membrane insertion or transmembrane transfer, the signal sequence is cleaved off by signal peptidase I (HVO_0002, HVO_2603) or signal peptidase II (still awaiting its identification in haloarchaea). After cleavage, the signal peptide must be degraded to avoid clogging of the membrane. Degradation is catalyzed by signal peptide peptidase. Candidates for this activity have been predicted from two protein families: the paralogs *sppA1* (HVO_0881) and *sppA2* (HVO_1987) are members of MEROPS family S49. A DUF1119 family protein (HVO_1107), which is a member of MEROPS family A22, has been predicted to function as signal peptide peptidase (Ng et al., 2007, Raut et al., 2021).

#### Section S9: Miscellaneous metabolic enzymes and proteins with other functions

##### S9.a A candidate gene for ketohexokinase

Ketohexokinase has been characterized in *Haloarcula vallismortis* (Rangaswamy and Altekar, 1994). Beyond enzymatic properties, only the amino acid composition has been reported. A speculative gene assignment was subsequently made (Anderson et al., 2011) in *Halomicrobium mukohataei* (Hmuk_2662, the ortholog of HVO_1812). This protein belongs to a sugar kinase family and is in genomic vicinity with fructose 1-phosphate kinase (Hmuk_2661) and fructose bisphosphate aldolase (Hmuk_2663). Further bioinformatic analyses supported this assignment (Williams et al., 2019). Homologs of Hmuk_2662 were found in six genomes, including species that are stated to have confirmed ketohexokinase activity (*Har. vallismortis, Har. marismortui, Hfx. mediterranei*). The amino acid composition of the homolog from *Har. vallismortis* (C437_04081) shows a close correlation with that of the characterized enzyme. Overall, bioinformatic analyses strongly support the assignment of ketohexokinase activity to Hmuk_2662 and C437_04081, with experimental confirmation still pending.

##### S9.b A candidate gene for fructokinase

The same paper which strongly supported the gene assignment to ketohexokinase (Williams et al., 2019) also assigned a gene to fructokinase. Differential proteomics in *Hht. litchfieldiae* grown in the presence and absence of sucrose showed an up-regulation of halTADL_1913 which, according to bioinformatic analyses, is a candidate for having fructokinase activity. A phylogenetic tree places halTADL_1913 into a cluster of 17 haloarchaeal genes. Among the most closely related sequences is a *Salmonella* gene product with 37% protein sequence identity that complements a fructose kinase deficiency (UniProt:P26984) (Aulkemeyer et al., 1991) and OCC_03567 from *Thermococcus litoralis* (31% protein sequence identity) which has been experimentally analyzed (Qu et al., 2004). Again, experimental confirmation may be possible by heterologous expression in *Hfx. volcanii* of halTADL_1913, OCC_03567, or another member of this sequence cluster.

**S9.c A candidate gene for glucoamylase**

There is a candidate ortholog set for having glucoamylase activity (*Hfx. volcanii* HVO_1711, *Haloquadratum walsbyi* HQ2539A, *Halorubrum sodomense* SAMN04487937_2677 [UniProt:A0A1I6HD35]). Glucoamylase from *Hrr. sodomense* (referred to in the older literature as *Halobacterium sodomense*) has been characterized (Chaga et al., 1993). Beyond enzymatic characterization, only the amino acid composition has been reported. The correlation between the reported amino acid composition and the theoretical amino acid composition of UniProt:A0A1I6HD35 is only moderate ($R^2$ = 0.72). UniProt:A0A1I6HD35 shows 51% protein sequence identity to HVO_1711. The C-terminal half of HVO_1711 shows 33% protein sequence identity to glucoamylase from *Clostridium* strain G0005 which has been characterized (Ohnishi et al., 1992). Taken together, there is clear bioinformatic support for assignment of that function but due to the high level of uncertainty experimental validation seems necessary.

**S9.d A candidate gene for glucose-6-phosphate isomerase**

There is a strong candidate for having glucose-6-phosphate isomerase activity is *Hfx. volcanii* (HVO_1967, *pgi*). It is a homolog to *M. jannaschii* MJ1605 (36% protein sequence identity) which has been characterized (Rudolph et al., 2004).

**S9.e A candidate gene for 2-dehydro-3-deoxy-(phospho)gluconate aldolase**

A candidate gene product for 2-dehydro-3-deoxy-(phospho)gluconate aldolase activity is *Hbt. salinarum* KdgA (OE_1665R, *kdgA*). It shows 36% protein sequence identity to *Hfx. volcanii* HVO_1101 (encoded by *dapA*), which is involved in lysine biosynthesis. However, there are no other lysine biosynthesis genes in *Hbt. salinarum*. Instead, OE_1665R may be isofunctional with much more distantly related proteins, KdgA from *Saccharolobus (Sulfolobus) solfataricus* (SSO3197) and *Thermoproteus tenax* (TTX_1156.1) which have been characterized (Ahmed et al., 2005). The function assignment to OE_1665R is shared by KEGG but not by UniProt (accessed March 2021). The assignment is supported by the adjacently encoded protein (OE_1664R) which catalyzes the preceding step in the pathway and another enzyme encoded in the vicinity (OE_1669F) which also codes for a step in the same pathway. Only few haloarchaea seem to have a close homolog of OE_1665R, including *Hbt. hubeiense*. This species, which codes for the enzymes of the lysine biosynthesis pathway, has two paralogs, one being close to OE_1665R (Hhub_1547, *kdgA*), the other close to HVO_1101 (Hhub_1591, *dapA*). OrthoDB places the paralogs from *Hbt. hubeiense* into the same group, so that the number of haloarchaeal genomes coding for an ortholog of OE_1665R cannot be easily retrieved. The function assignment to OE_1665R and its close orthologs has to be considered uncertain until its experimental confirmation (or rejection).

**S9.f A putative NAD-independent L-lactate dehydrogenase**

LudBC (e.g. HVO_1692 and HVO_1693) is a probable NAD-independent L-lactate dehydrogenase which converts lactate into pyruvate. Lactate is one of the products of rhamnose catabolism, and in some haloarchaea *ludBC* gene pairs are found within the rhamnose catabolic cluster (Reinhardt et al., 2019). In *Hfx. volcanii*, the *ludBC* gene pair is not part of the rhamnose catabolic cluster, but deletion of *ludBC* (HVO_1692 and HVO_1693) impairs growth on rhamnose (Reinhardt et al., 2019). LudBC orthologs are found in many haloarchaea, with gene clustering being highly conserved. Both, *ludC* and *ludB* contain an assigned InterPro domain (LUD, previously DUF162, IPR003741).

A NAD-independent L-lactate dehydrogenase (*lutABC* in *B. subtilis* (Chai et al., 2009); *lldABC* in *Pseudomonas stutzeri* (Gao et al., 2015)) has been characterized. LutC/LldC is very distantly related (up to 33% protein sequence identity, BLASTp E-value $10^{-8}$) to homologs from *Hfx. volcanii* (HVO_1693, encoded by *ludC*) and *Nmn. pharaonis* (NP_1724A), which show 44% protein sequence identity to each other. LutB/LldB is related (up to 38% protein sequence identity) to the N-terminal part (up to ca pos 490 of 733) of the homologs from *Hfx. volcanii* (HVO_1692, encoded by *ludB*) and *Nmn. pharaonis* (NP_1722A) which show 60% protein sequence identity to each other.

Upon superficial analysis, it seems that *Hfx. volcanii* and *Nmn. pharaonis* lack homologs to *B. subtilis* LutA and *Pseudomonas* LldA. LutA and LldA show 42% protein sequence identity to each other and have a pair of IPR004017 domains assigned. LutA and LldA are distantly related to the C-terminal part of GlpC (e.g. HVO_1540, glycerol-3-phosphate dehydrogenase subunit C, 25% protein sequence identity, E-value $10^{-9}$) and other FAD-dependent oxidoreductases. LldA but not LutA is very distantly related to the C-terminal part of LudB (27% protein sequence identity, E-value $10^{-5}$). Neither *Hfx. volcanii* nor *Nmn. pharaonis* LudB have an IPR004017 domain assigned but have a C-terminal part with no other InterPro domain assignment. Thus, , haloarchaeal *ludB* may well represent a domain fusion (*lutBA*, *lldBA*). In that case, HVO_1692 and HVO_1693 would correspond to a complete heterotrimeric NAD-independent L-lactate dehydrogenase and would be promising candidates for having NAD-independent L-lactate dehydrogenase activity.

In *Pseudomonas*, *lldB* (PST_3338) and *lldC* (PST_3339) are clustered with genes for D-lactate dehydrogenase (PST_3340) and for a lactate permease (PST_3336). In the genomic vicinity of *Haloferax* ludBC is HVO_1696, a LctP family transport protein with 44% protein sequence identity to the *Pseudomonas* lactate permease. HVO_1697 shows 24% protein sequence identity to PST_3340. It cannot be excluded that those genes are also involved in lactate metabolism in *Haloferax*.

**S9.g A gene cluster which may be responsible for conversion of urate to allantoin and may be part of a complete purine degradation pathway**

We reconstructed a four-step pathway for conversion of urate to allantoate. According to KEGG (accessed April 2021), purines are degraded with intermediates being xanthine, urate and allantoate, the latter being further decomposed to $CO_2$ and glyoxylate. The four-step pathway is annotated according to a gene cluster from *B. subtilis* which codes for enzymes with the corresponding activities. The enzymes for the first and third pathway step are encoded by a single fused gene in *B. subtilis* (*pucL*) (Pfrimer et al., 2010, Kim et al., 2007). The enzymes are encoded by independent genes in haloarchaea (e.g. HVO_B0301, *pucL2*, corresponding to the N-terminus of *B. subtilis* PucL, HVO_B0300, *pucL1*, corresponding to the C-terminus of *B. subtilis* PucL). The second pathway step is catalyzed by the product of *pucM* (Lee et al., 2005). The above assignments have also been made by KEGG (as of April 2021). The fourth step is catalyzed by allantoinase, a member of the cyclic amidohydrolase family. HVO_B0303 has only 30% protein sequence identity to characterized proteins with this activity (from *B. subtilis* and from *Salmonella* (Schultz et al., 2001, Ho et al., 2013), but is slightly closer (33% protein sequence identity) to *Ralstonia* (*Burkholdia*) D-hydantoinase, which is involved in pyrimidine catabolism (Xu et al., 2002). Purine and pyrimidine degradation pathways contain successive reactions which display similar chemistry and are catalyzed by homologous proteins (Barba et al., 2013). KEGG (accessed April 2021) annotates HVO_B0302 as dihydropyrimidinase (D-hydantoinase), thus being involved in pyrimidine degradation, while we predict it as allantoinase, involved in purine degradation, taking the function of products from clustered genes into account. There is a paralog with 61% protein sequence identity (HVO_A0303) which does not show function-related gene clustering. The pathway step subsequent to D-hydantoinase in pyrimidine catabolism is catalyzed by EC 3.5.1.6 (beta-ureidopropionase) and KEGG assigns this function to three *Haloferax* paralogs (as of April 2021), one being HVO_B0306 (encoded by *amaB4*). The chemically similar reaction in the purine degradation pathway is EC 3.5.1.116 (ureidoglycolate hydrolase). Given the close proximity of the genes coding for other purine degradation pathway enzymes, we have assigned this function to HVO_B0306. Support comes from a characterized *Arabidopsis* homolog (34% protein sequence identity) (Werner et al., 2010, Werner et al., 2013) but a more closely related protein from *Geobacillus* (39% protein sequence identity) has a distinct function (Martinez-Rodriguez et al., 2012). Two paralogs of HVO_B0306 in *Haloferax*, HVO_A0341 and HVO_2128, show more than 50% protein sequence identity but do not show function-related gene clustering and are currently not annotated as catalyzing EC 3.5.1.116. With these assignments, only two pathway gaps remain for xanthine degradation. The conversion of allantoate to ureido-glycolate, the substrate of EC 3.5.1.116, is yet unassigned. The other gap is the conversion of xanthine to urate. The genome region codes in a three-gene operon for the subunits of a molybdopterin-containing oxidoreductase (HVO_B0308, small subunit, HVO_B0309, large subunit, HVO_B0310, medium subunit). These are distantly related to enzymes acting on various substrates (e.g. carbon monoxide). Among those is a probable xanthine dehydrogenase from *E. coli* which has been partially characterized (Xi et al., 2000). It should be noted that xanthine oxidase activity has not been detected in *Hfx. volcanii* and that xanthine was not metabolized (Stuerlauridsen and Nygaard, 1998). The proposed purine degradation pathway, if it exists, may not be expressed under optimal growth conditions. Finally, one gene in this cluster (HVO_B0303) codes for a transport protein for which xanthine and uracil are currently assigned as potential substrates. It shows 38% protein sequence identity to the product of *E. coli xanP*, a characterized xanthine permease (Karatza and Frillingos, 2005).

Many of the assignments are insecure due to the low sequence similarity of functionally characterized homologs, and because homologs with distinct functions are equidistant for several of the enzymes. However, some enzymes are clearly related to purine catabolism and the clustering of these genes makes it likely that the complete region from HVO_B0299 to HVO_B0310 codes for proteins involved in purine catabolism. These assignments will remain speculative until their experimental confirmation (or rejection).

**S9.h *Hfx. volcanii* may have an enzyme with a nickel-pincer cofactor, generated by *larBCE***

HVO_2381 is annotated as LarC family protein, based on the product of the *Lactobacillus plantarum larC* gene which has been characterized (Desguin et al., 2016). The *Lactobacillus* LarC forms a complex with LarB and LarE, which generates the "nickel-pincer cofactor" of lactate racemase (LarA). The corresponding genes are part of an operon (*larA-larB-larC1-larC2-larD-larE*) and a regulatory gene (*larR*) is found nearby and transcribed in the opposite direction.

However, in the case of *Hfx. volcanii* DS2, the LarC family protein gene occurs without other *lar* homologs nearby, and not within an operon. In this and other species of haloarchaea, HVO_2381 is regularly found nearby an oppositely facing gene encoding AAA-type ATPase (CDC48 subfamily).

Later studies (Desguin et al., 2017, Hausinger et al., 2018) identified many prokaryotic genomes with *larBCE* homologs that lack *larA*, and suggested from those findings that LarB, C and E can be used by enzymes other than LarA, and so for other functions. In *Hfx. volcanii* DS2 there are distant homologs of *larB* (HVO_0197) and of *larE* (HVO_0190) elsewhere in the chromosome but neither are part of an operon. In summary, the function of HVO_2381 is not clear even though a standard BLASTp search at GenBank (nr) returns a conserved protein domain match to PRK04194, “nickel pincer cofactor biosynthesis protein LarC” with high probability ($3.98x10^{-138}$) (accessed January, 2021).

##### S9.i Candidate genes which may catalyze cyclic di-AMP hydrolysis

Cyclic di-AMP (c-di-AMP) is generated from two molecules of ATP by diadenylate cyclase (encoded by *dacZ*) (Corrigan and Grundling, 2013). *Hfx. volcanii* DacZ (HVO_1660) has been characterized and it was shown that c-di-AMP levels must be tightly regulated (Braun et al., 2019). c-di-AMP is degraded to pApA by phosphodiesterases (Corrigan and Grundling, 2013). As the level of this signalling molecule needs to be strictly controlled (Gundlach et al., 2015) (Commichau et al., 2019), the degrading enzyme is also an important player. In *Hfx. volcanii*, this enzyme has yet not been identified but candidates have been proposed (Corrigan and Grundling, 2013, Yin et al., 2020). These are proteins with a DHH and DHHA1 domain that also have an N-terminal trkA_N domain. *Haloferax* candidates HVO_0990 and HVO_1690 were proposed recently, based on a phylogenetic profile (He et al., 2020). Also proposed was HVO_0756, which, however, lacks the N-terminal trkA_N domain. Gene neighbourhood analysis did not provide additional clues to support or challenge the involvement of one or several of these candidates.

##### S9.j A homolog to RNaseZ

HVO_2763 shows 27% protein sequence identity to RNase Z (HVO_0144, *rnz*). The experimental characterization of HVO_2763 (Fischer et al., 2012) specifically excluded its activity as a tRNA or 5S RNA processing enzyme and as an exonuclease generally. However, it did not reveal the physiological function of HVO_2763. The authors point out that this protein belongs to the metallo-beta-lactamase family, which is known to fulfil a diversity of functions. The authors deleted the gene, performed a transcriptome analysis, and found several genes to be downregulated in the deletion mutant. Among these were several transporter genes, and the authors note that the gene for HVO_2763 is adjacent to a pair of permease genes. However, SyntTax analysis shows that the permease genes are a strain-specific insertion in *Hfx. volcanii* DS2 when compared to closely related genomes. In other genomes, the adjacent gene corresponds to *lhr1* (HVO_2766) which codes for a helicase. The HVO_2763/2766 clustering is not only found in closely related strains, but in a considerable number of more distantly related haloarchaeal genomes. This should be considered when searching for a physiological function of

HVO_2763. Additionally, three regulated genes are clustered and have since then been characterized (HVO_2055, *agl15*, HVO_2060, *agl8*, HVO_2061, *agl6*) (Kaminski et al., 2013). They participate in a N-glycosylation pathway that was initially observed mainly under low-salt conditions. With respect to the downregulated transporter genes no annotation updates have been made since publication of the results.

##### S9.k A potential CO2 transporter (*dabAB*)

HVO_2410 and HVO_2411 probably function as carbon dioxide transporter, based on the identification of such transporters in *Halothiobacillus neapolitanus* (Desmarais et al., 2019). The authors assume that CO2 is directly transported and not a hydrated form (e.g. hydrogen carbonate), because transport activity is pH independent. This transporter is one of the $CO_2$ concentrating mechanisms. Orthologs are found in about two-thirds of the haloarchaeal genomes (108 of 166) according to OrthoDB. HVO_2411 belongs to the proton-conducting membrane transporter family (PFAM PF00361, InterPro IPR001750), which also contains subunits of the *nuo* complex and of the Mrp-type sodium/proton antiporter. In KEGG, HVO_2411 is annotated as NAD(P)H-quinone oxidoreductase subunit 5 (as of April 2021).
