## Supplementary material for "Open issues for protein function assignment in *Haloferax volcanii* and other halophilic archaea": Suppl.Tables S10 and S11

| **Section** | **Code** (*Hfx. volcanii*) | **Gene** | **Code** (other species or paralog) | **Gene** | **%seq_id** | **Comment** |
|---|---|---|---|---|---|---|
| 1a | **HVO_1305** | *porA* | OE2623R | *porA* | 67% | |
| 1a | **HVO_1304** | *porB* | OE2622R | *porB* | 80% | |
| 1a | **HVO_0888** | *korA* | OE1711R | *korA* | 77% | |
| 1a | **HVO_0887** | *korB* | OE1710R | *korB* | 77% | |
| 1a/1b | HVO_2995 | *fdx* | OE4217R | *fdx* | 88% | |
| 1c | **HVO_0978** | *nuoA* | | | | |
| 1c | **HVO_0979** | *nuoB* | | | | |
| 1c | **HVO_0980** | *nuoCD1* | | | | gene fusion |
| 1c | **HVO_0981** | *nuoH* | | | | |
| 1c | **HVO_0982** | *nuoI* | | | | |
| 1c | **HVO_0983** | *nuoJ1* | | | | N-term of NuoJ |
| 1c | **HVO_0984** | *nuoJ2* | | | | C-term of NuoJ |
| 1c | **HVO_0985** | *nuoK* | | | | |
| 1c | **HVO_0986** | *nuoL* | | | | |
| 1c | **HVO_0987** | *nuoM* | | | | |
| 1c | **HVO_0988** | *nuoN* | | | | |
| 1c | HVO_0968 | *nuoCD2* | HVO_0980 | *nuoCD1* | 75% | |
| 1d | HVO_1578 | *ndh1* | NP_3508A | *ndh* | | |
| 1d | HVO_1413 | *ndh2* | HVO_1578 | *ndh1* | 75% | |
| 1e | **HVO_2620** | *petA* | OE_1876R | *petA* | 54% | |
| 1e | **HVO_0842** | *petB* | OE_1874R | *petB* | 81% | |
| 1e | **HVO_0841** | *petD* | OE_1872R | *petD* | 77% | |
| 1f | **HVO_2808** | *sdhA* | NP_4264A | *sdhA* | 76% | |
| 1f | **HVO_2809** | *sdhB* | NP_4266A | *sdhB* | 76% | |
| 1f | **HVO_2810** | *sdhD* | NP_4268A | *sdhD* | 66% | |
| 1f | **HVO_2811** | *sdhC* | NP_4270A | *sdhC* | 81% | |
| 1g/1h | HVO_0943 | *cbaD* | NP_2966A | *cbaD* | 57% | |
| 1g/1h | HVO_0943 | *cbaD* | OE_4073R | *hcpB* | 63% | matches to C-term |
| 1g | HVO_2150 | *hcpG* | OE_4073R | *hcpB* | 44% | matches to N-term |
| 1h | **HVO_0945** | *cbaA* | NP_2962A | *cbaA* | 64% | |
| 1h | **HVO_0944** | *cbaB* | NP_2964A | *cbaB* | 60% | |
| 1g/1h | **HVO_0943** | *cbaD* | NP_2966A | *cbaD* | 57% | |

| | | | | | | |
|---|---|---|---|---|---|---|
| 1h | **HVO_0942** | *cbaE* | NP_2968A | *cbaE* | 45% | |
| 1h | HVO_0945 | *cbaA* | OE_4070R | *cbaA* | 65% | |
| 1h | HVO_0944 | *cbaB* | OE_4071R | *cbaB* | 56% | |
| 1g/1h | HVO_0943 | *cbaD* | OE_4073R | *hcpB* | 63% | gene function in *Hbt. salinarum*; matches to C-term |
| 1h | HVO_0942 | *cbaE* | - | - | | no ortholog in *Hbt. salinarum* |
| 1h | **HVO_0907** | *coxA1* | NP_2456A | *coxA1* | 68% | |
| 1h | **HVO_1014** | *coxB1* | NP_2448A | *coxB1* | 50% | |
| 1h | **HVO_1138** | *coxC1* | NP_2450A | *coxC1* | 59% | |
| 1h | HVO_0907 | *coxA1* | OE_1979R | *coxA1* | 70% | |
| 1h | HVO_1014 | *coxB1* | OE_1988R | *coxB1* | 43% | |
| 1h | HVO_1138 | *coxC1* | OE_1984F | *coxC1* | 65% | |
| 1h | **HVO_1645** | *coxAC2* | | | | |
| 1h | **HVO_1646** | *coxB2* | | | | |
| 1h | **HVO_1644** | *coxD* | | | | |
| 1h | HVO_1645 | *coxAC2* | HVO_0907 | *coxA1* | 39% | matches to N-term |
| 1h | HVO_1645 | *coxA23* | HVO_1138 | *coxC1* | 28% | matches to C-term |
| 1h | **HVO_0462** | *cydA* | OE_6185F OE_7065F | *cydA* | 56% | encoded on a plasmid pHS1/pHS2 duplication |
| 1h | **HVO_0461** | *cydB* | OE_6186F OE_7066F | *cydB* | 45% | encoded on a plasmid pHS1/pHS2 duplication |
| 1h | **NP_4296A** | *coxA3* | Hhub_2087 | | 60% | |
| 1h | **NP_4294A** | *coxB3* | Hhub_2086 | | 57% | |
| 1h | NP_4296A | *coxA3* | NP_2962A | *cbaA* | 28% | |
| 1h | NP_4294A | *coxB3* | NP_2964A | *cbaB* | - | 33% seq_id in the C-term ca 80 aa |
| 1i | **HVO_2958** | *oadhA1* | | | | component E1 alpha subunit |
| 1i | **HVO_2959** | *oadhB1* | | | | component E1 beta subunit |
| 1i | **HVO_2960** | *dsa1* | | | | component E2 |
| 1i | **HVO_2961** | *lpdA* | | | | component E3 |
| 1i | HVO_2209 | *oadhA4* | | | | component E1 alpha subunit; no gene cluster |
| 1i | **HVO_0669** | *oadhA3* | | | | component E1 alpha subunit |
| 1i | **HVO_0668** | *oadhB3* | | | | component E1 beta subunit |
| 1i | **HVO_0667** | - | | | | homolog to NAD kinase |
| 1i | **HVO_0666** | *dsa2* | | | | component E2 |
| 1i | **HVO_2595** | *oadhA2* | | | | component E1 alpha subunit |
| 1i | **HVO_2596** | *oadhB2* | | | | component E1 beta subunit |
| 1i | **HVO_2597** | *oadhL* | | | | a short protein consisting of a lipoyl domain |

| | | | | | | |
|---|---|---|---|---|---|---|
| 1i | HVO_2958 | *oadhA1* | HVO_2209 | *oadhA4* | 59% | |
| 1i | HVO_2958 | *oadhA1* | HVO_0669 | *oadhA3* | 33% | |
| 1i | HVO_2958 | *oadhA1* | HVO_2595 | *oadhA2* | 33% | |
| 1i | HVO_0669 | *oadhA3* | HVO_2595 | *oadhA2* | 41% | |
| 1i | HVO_0669 | *oadhA3* | HVO_2209 | *oadhA4* | 30% | |
| 1i | HVO_2595 | *oadhA2* | HVO_2209 | *oadhA4* | 32% | |
| 1i | HVO_2959 | *oadhB1* | HVO_0668 | *oadhB3* | 43% | |
| 1i | HVO_2959 | *oadhB1* | HVO_2596 | *oadhB2* | 41% | |
| 1i | HVO_0668 | *oadhB3* | HVO_2596 | *oadhB2* | 42% | |
| 1i | HVO_2960 | *dsa1* | HVO_0666 | *dsa2* | 31% | |
| 1i | HVO_2597 | *oadhL* | HVO_2960 | *dsa1* | - | 38% seq_id in N-term ca 80 aa of *dsa1* |
| 1i | HVO_2597 | *oadhL* | HVO_0666 | *dsa2* | - | 35% seq_id in N-term ca 80 aa of *dsa2* |
| 1i | HVO_2597 | *oadhL* | HVO_2486 | *pccA* | | 35% seq_id in N-term ca 80 aa of *pccA* |
| 2a | HVO_0041 | *argF* | | | | Arg biosynthesis |
| 2a | HVO_0042 | *argE* | | | | Arg biosynthesis |
| 2a | HVO_0043 | *argD* | | | | Arg biosynthesis |
| 2a | HVO_0044 | *argB* | | | | Arg biosynthesis |
| 2a | HVO_0045 | *argC* | | | | Arg biosynthesis |
| 2a | HVO_0046 | *argX* | | | | Arg biosynthesis |
| 2a | HVO_0047 | *argW* | | | | Arg biosynthesis |
| 2a | HVO_0048 | *argH* | | | | Arg biosynthesis |
| 2a | HVO_0049 | *argG* | | | | Arg biosynthesis |
| 2a | HVO_0008 | *lysC* | | | | Lys biosynthesis |
| 2a | HVO_1096 | *dapE* | | | | Lys biosynthesis |
| 2a | HVO_1097 | *dapF* | | | | Lys biosynthesis |
| 2a | HVO_1098 | *lysA* | | | | Lys biosynthesis |
| 2a | HVO_1099 | *dapD* | | | | Lys biosynthesis |
| 2a | HVO_1100 | *dapB* | | | | Lys biosynthesis |
| 2a | HVO_1101 | *dapA* | | | | Lys biosynthesis |
| 2a | HVO_2487 | *asd* | | | | Lys biosynthesis |
| 2a | HVO_A0634 | - | HVO_1096 | *dapE* | 25% | KEGG annotates HVO_A0634 also as DapE |
| 2b | HVO_0790 | *fba2* | | | | |
| 2b | HVO_0792 | *aroB* | | | | |
| 2b | HVO_0602 | *aroD1* | HVO_0603 | *aroD2* | 45% | |
| 2b | HVO_0603 | *aroD2* | | | | |
| 2c | HVO_0009 | *tnaA* | | | | |
| 2d | HVO_A0559 | *hutH* | | | | |

| | | | | | | |
|---|---|---|---|---|---|---|
| 2d | HVO_A0562 | *hutU* | | | | |
| 2d | HVO_A0560 | *hutI* | | | | |
| 2d | HVO_A0561 | *hutG* | | | | |
| 2e | HVO_0431 | - | | | | |
| 2e/7b | HVO_1295 | *hisC* | | | | |
| 2e | HVO_1153 | - | | | | |
| 2e | HVO_0644 | *leuA1* | HVO_1510 | *leuA2* | 39% | |
| 2e | HVO_0644 | *leuA1* | HVO_A0489 | - | 37% | |
| 2e/2f | HVO_1510 | *leuA2* | HfgLR_07660 | *leuA2* | 99% | Met-1/Met-1 |
| 2e/2f | HVO_1510 | *leuA2* | HFX_1573 | *leuA2* | 95% | Leu-27/Leu-1; no start codon |
| 2e/2f | HVO_1510 | *leuA2* | HQ2700A | *leuA1* | 76% | Cys-30/Cys-10 |
| 2e/2f | HVO_1510 | *leuA2* | Hqrw_3051 | *leuA1* | 76% | Cys-30/Cys-10 |
| 2e/2f | HVO_1510 | *leuA2* | rrnAC0329 | *leuA2* | 78% | Cys-30/Cys-8 |
| 2e/2f | HVO_1510 | *leuA2* | HAH_1067 | *leuA1* | 77% | Cys-30/Cys-8 |
| 2e/2f | HVO_1510 | *leuA2* | Nmag_0906 | *leuA1* | 73% | Cys-30/Cys-18 |
| 2e/2f | HVO_1510 | *leuA2* | HBSAL_05970 | *leuA* | 68% | Phe-46/Phe-6; no start codon |
| 2e/2f | HVO_1510 | *leuA2* | Hhub_3057 | *leuA1* | 70% | Glu-45/Glu-7 |
| 2e/2f | HVO_1510 | *leuA2* | NP_2206A | *leuA1* | 79% | Arg-36/Arg-4; no start codon |
| 2e/2f | HVO_1510 | *leuA2* | Nmlp_3164 | *leuA1* | 76% | Arg-36/Arg-4; no start codon |
| 2e/2f | HVO_1510 | *leuA2* | halTADL_0359 | *leuA1* | 78% | Cys-30/Cys-1; no start codon |
| 2e | HVO_A0489 | - | | | | |
| 2e | HVO_1153 | - | | | | |
| 2e | HVO_1154 | - | | | | |
| 2e | HVO_1155 | - | | | | |
| 3a | HVO_B0054 | *cbiX1* | NP1108A | *cbiX1* | 71% | |
| 3a | HVO_1128 | *cbiX2* | NP1588A | *cbiX2* | 79% | |
| 3a | HVO_1128 | *cbiX2* | NP0734A | *cbiX3* | 26% | |
| 3a | HVO_1128 | *cbiX2* | Nmag_3212 | *cbiX* | 79% | no de novo cobalamin biosynthesis genes in *Nab. magadii* |
| 3a | - | | NP_0734A | *cbiX3* | | no ortholog in *Hfx.volcanii* |
| 3a | HVO_2312 | *sirC* | | | | |
| 3b | HVO_B0061 | *cbiL* | | | | |
| 3b | HVO_B0057 | *cbiH2* | HVO_B0058 | *cbiH1* | 38% | |
| 3b | HVO_B0058 | *cbiH1* | | | | |
| 3b | HVO_B0060 | *cbiF* | | | | |
| 3b | HVO_B0059 | *cbiG* | | | | |
| 3b | HVO_B0062 | *cbiT* | | | | |

| | | | | | | |
|---|---|---|---|---|---|---|
| 3b | HVO_B0048 | *cbiE* | | | | |
| 3b | HVO_B0049 | *cbiC* | | | | |
| 3b | HVO_A0487 | *cbiA* | | | | |
| 3b | HVO_B0059 | *cbiG* | G3A49_06890 | | 98% | |
| 3b | HVO_B0058 | *cbiH1* | G3A49_06895 | | 94% | |
| 3b | HVO_B0057 | *cbiH2* | G3A49_06900 | | 99% | |
| 3b | HVO_B0056 | | G3A49_06905 | | 97% | |
| 3b | HVO_B0055 | | G3A49_06910 | | 90% | |
| 3b | HVO_B0054 | *cbiX1* | G3A49_06915 | | 98% | |
| 3b | HVO_B0053 | | G3A49_06920 | | 99% | |
| 3b | HVO_B0052 | | G3A49_06925 | | 99% | |
| 3b | HVO_B0051 | *chlID* | G3A49_06930 | | 97% | |
| 3b | HVO_B0050 | *cobN* | G3A49_06935 | | 98% | |
| 3b | HVO_B0049 | *cbiC* | G3A49_06940 | | 99% | |
| 3b | HVO_B0048 | *cbiE* | G3A49_06945 | | 94% | |
| 3c | HVO_A0488 | *cobA* | | | | |
| 3c | HVO_2395 | *pduO* | | | | |
| 3c | HVO_A0553 | *cbiP* | | | | |
| 3c | HVO_0587 | *cbiB* | | | | |
| 3c | HVO_0592 | *cbiZ* | | | | |
| 3c | HVO_0589 | *cobY* | | | | |
| 3c | HVO_0588 | *cobS* | | | | |
| 3c | HVO_0586 | - | VNG_1577C<br>OE_3249F | - | 56% | predicted to be isofunctional to *cobC* |
| 3c | HVO_0591 | *cobD1* | | | | |
| 3c | HVO_0593 | *cobD2* | HVO_0591 | *cobD1* | 51% | |
| 3c | HVO_0590 | *cobT* | VNG_1572C<br>OE_3242F | *cobT* | 57% | |
| 3c | - | | halTADL_3045 | *cobT* | | |
| 3d | HVO_B0051 | *cobN* | | | | |
| 3d | HVO_B0050 | *chlID* | | | | |
| 3e | HVO_1121 | *ahbC* | | | | |
| 3e | HVO_2144 | *ahbD* | | | | |
| 3e | HVO_2227 | *ahbA* | | | | |
| 3e | HVO_2313 | *ahbB* | HVO_2227 | *ahbA* | 39% | |
| 4a | HVO_2198 | *cofH* | | | | |
| 4a | HVO_2201 | *cofG* | | | | |

| | | | | | | |
|---|---|---|---|---|---|---|
| 4a | HVO_2202 | *cofC* | | | | |
| 4a | HVO_2479 | *cofD* | | | | |
| 4a | HVO_1936 | *cofE* | | | | |
| 4b | HVO_0433 | *npdG* | | | | |
| 4b | HVO_B0113 | - | | | | |
| 4b | HVO_B0342 | - | | | | |
| 4b | HVO_A0572 | - | NP_1902A | - | 44% | |
| 4b | NP_1902A | - | NP_4374A | - | 33% | |
| 4b | - | | NP_4006A | - | | |
| 4b | HVO_A0605 | - | | | | |
| 4c/5c | HVO_1937 | *mer* | | | | |
| 4d | HVO_2911 | *phr2* | HVO_2843 | *phr1* | 31% | |
| 4d | HVO_2843 | *phr1* | | | | |
| 4d | HVO_1234 | *phr3* | | | | |
| 5a | HVO_0709 | *pabA* | HVO_2453 | *trpG* | 44% | |
| 5a | HVO_0710 | *pabB* | HVO_2454 | *trpE* | 37% | |
| 5a | HVO_0708 | *pabC* | HVO_0329 | *ilvE* | 32% | |
| 5b | HVO_2348 | *mptA* | | | | |
| 5b | HVO_A0533 | - | | | | |
| 5b | HVO_2628 | - | | | | |
| 5c | HVO_2573 | *mch* | | | | |
| 4c/5c | HVO_1937 | *mer* | | | | |
| 6a | HVO_2363 | *nadK1* | HVO_0837 | *nadK2* | 25% | |
| 6a | HVO_0837 | *nadK2* | | | | |
| 6b | HVO_0782 | *nadM* | | | | |
| 6b | HVO_0781 | - | | | | |
| 6b | HVO_0326 | *rbkR* | | | | |
| 6b | HVO_0327 | *ribB* | | | | |
| 6b | HVO_0974 | *ribH* | | | | |
| 6b | HVO_1015 | *ribL* | | | | |
| 6b | HVO_1284 | *arfA* | | | | |
| 6b | HVO_1235 | - | | | | *arfB* candidate |
| 6b | HVO_1341 | *arfC* | | | | |
| 6b | HVO_2483 | - | | | | *RIB2* candidate |
| 6b | HVO_0326 | *rbkR* | | | | |
| 6b | HVO_1015 | *ribL* | | | | |
| 7a | HVO_0303 | *idsA2* | NP_0604A | *idsA2* | 66% | |

| | | | | | | |
|---|---|---|---|---|---|---|
| 7a | HVO_2725 | *idsA1* | NP_3996A | *idsA3* | 67% | |
| 7a | - | | NP_4556A | *idsA1* | | |
| 7b | HVO_0332 | *carS* | | | | |
| 7b | HVO_1136 | *pgsA1* | | | | |
| 7b | HVO_1143 | *assA* | VNG_0784G<br>OE_2155R | *assA* | 56% | |
| 7b | HVO_1297 | *aisA* | VNG_1030G<br>OE_2503R | *agsA* | 64% | |
| 7b | HVO_1971 | *pgsA4* | | | | |
| 7b | HVO_1297 | *aisA* | NP2144A | *aisA* | 66% | |
| 7b | HVO_1135 | - | | | | |
| 7b | HVO_0146 | *asd* | VNG_2255C<br>OE_4164F | *asd* | 61% | |
| 2e/7b | HVO_1295 | *hisC* | | | | |
| 7b | HVO_1296 | *adk2* | | | | |
| 7b | HVO_2496 | *adk1* | | | | |
| 7b | HVO_B0213 | - | | | | |
| 7c | HVO_2524 | *crtB* | NP_4770A | *crtB* | 57% | |
| 7c | HVO_2527 | *lyeJ* | NP_4766A | *lyeJ* | 61% | |
| 7c | HVO_2528 | *crtD* | NP_4764A | *crtD* | 72% | |
| 7c | HVO_2528 | *crtD* | NP_0204A | *crtI2* | 57% | |
| 7c | HVO_2526 | *cruF* | NP_4768A | *cruF* | 60% | |
| 7c | HVO_0817 | - | NP_1630A | - | 54% | |
| 7c | HVO_0817 | - | HFX_0786 | - | 88% | |
| 7c | HVO_2340 | - | NP_4520A | - | 69% | |
| 7c | NP_4764A | *crtD* | NP_0204A | *crtI2* | 60% | |
| 7c | NP_0204A | *crtI2* | NP_4520A | - | 26% | |
| 7c | NP_0204A | *crtI2* | NP_1630A | - | 26% | |
| 7c | HVO_2529 | - | NP_4762A | - | 60% | |
| 7d | HVO_1470 | *menF* | | | | |
| 7d | HVO_1469 | *menD* | | | | |
| 7d | HVO_1461 | *menC* | | | | |
| 7d | HVO_1375 | *menE* | | | | |
| 7d | HVO_1465 | *menB* | | | | |
| 7d | HVO_1462 | *menA* | | | | |
| 7d | HVO_0309 | *menG* | | | | |
| 7d | HVO_1463 | *trxB4* | | | | in *men* gene cluster |

| | | | | | | |
|---|---|---|---|---|---|---|
| 7d | HVO_1464 | - | | | | in *men* gene cluster |
| 7d | HVO_1466 | - | | | | in *men* gene cluster |
| 7d | HVO_1467 | - | | | | in *men* gene cluster |
| 7d | HVO_1468 | - | | | | in *men* gene cluster |
| 8a | - | | OE_1279R | *rpoeps* | | |
| 8b | HVO_0360 | *rps10a* | | | | |
| 8b | HVO_1392 | *rps10b* | HVO_0360 | *rps10a* | 24% | |
| 8b | HVO_0361 | *tef1a1* | | | | |
| 8b | HVO_1391 | - | | | | |
| 8b | HVO_2550 | *rps14* | NP_4882A | *rps14a* | 75% | full-length |
| 8b | HVO_2550 | *rps14* | NP_1768A | *rps14b* | 80% | partial |
| 8c | HVO_0654 | *rpl43e* | OE_1373R | *rpl43e* | 53% | zinc finger only in OE_1373R |
| 8d | HVO_1631 | *dph2* | | | | |
| 8d | HVO_0916 | *dph5* | | | | |
| 8d | HVO_1077 | *dph6* | | | | |
| 8e | HVO_0881 | *sppA1* | | | | |
| 8e | HVO_1987 | *sppA2* | HVO_0881 | *sppA1* | 28% | |
| 8e | HVO_1107 | - | | | | |
| 8e | HVO_0002 | *sec11b* | | | | |
| 8e | HVO_2603 | *sec11a* | HVO_0002 | *sec11b* | 54% | |
| 9a | HVO_1812 | - | Hmuk_2662 | - | 50% | |
| 9b | - | | halTADL_1913 | - | | |
| 9c | HVO_1711 | - | | | | |
| 9d | HVO_1967 | *pgi* | | | | |
| 2a/9e | HVO_1101 | *dapA* | OE_1665R | *kdgA* | 36% | |
| 2a/9e | HVO_1101 | *dapA* | Hhub_1591 | *dapA* | 69% | |
| 2a/9e | HVO_1101 | *dapA* | Hhub_1547 | *kdgA* | 32% | |
| 9e | OE_1665R | *kdgA* | Hhub_1547 | *kdgA* | 77% | |
| 9e | OE_1665R | *kdgA* | Hhub_1591 | *dapA* | 37% | |
| 9e | HVO_1488 | *gnaD* | OE_1664R | *gnaD* | 88% | |
| 9e | HVO_1083 | *gdh* | OE_1669F | *gdh* | 62% | |
| 9f | HVO_1692 | *ludB* | NP_1724A | *ludB* | 60% | |
| 9f | HVO_1693 | *ludC* | NP_1724A | *ludC* | 44% | |
| 9f | HVO_1697 | - | | | | |
| 9f | HVO_1696 | *lctP* | | | | |
| 9g | HVO_B0300 | *pucL1* | | | | |
| 9g | HVO_B0299 | *pucM* | | | | |

| | | | | | | |
|---|---|---|---|---|---|---|
| 9g | HVO_B0301 | *pucL2* | | | | |
| 9g | HVO_B0302 | *pucH1* | HVO_A0303 | *pucH2* | 61% | |
| 9g | HVO_B0306 | *amaB4* | HVO_2128 | *amaB* | 55% | |
| 9g | HVO_B0306 | *amaB4* | HVO_A0341 | *amaB* | 54% | |
| 9g | HVO_B0308 | *coxS* | | | | |
| 9g | HVO_B0309 | *coxL* | | | | |
| 9g | HVO_B0310 | *coxM* | | | | |
| 9g | HVO_B0303 | *uraA4* | | | | |
| 9h | HVO_0197 | - | | | | |
| 9h | HVO_2381 | - | | | | |
| 9h | HVO_0197 | - | | | | |
| 9i | HVO_1660 | *dacZ* | | | | |
| 9i | HVO_0990 | - | HVO_1690 | - | 39% | full-length similarity |
| 9i | HVO_0990 | - | HVO_0756 | - | 38% | similarity starts at pos 140/55 |
| 9i | HVO_1690 | - | HVO_0756 | - | 32% | similarity starts at pos 129/32 |
| 9i | HVO_0756 | - | | | | |
| 9j | HVO_2763 | - | HVO_0144 | *rnz* | 27% | |
| 9k | HVO_2410 | *dabA* | | | | |
| 9k | HVO_2411 | *dabB* | HVO_0986 | *nuoL* | 29% | up to pos 328 of 494; up to pos 372 of 678 |
| 9k | HVO_2411 | *dabB* | HVO_0987 | *nuoM* | 24% | up to pos 325 of 494; up to pos 362 of 509 |
| 9k | HVO_2411 | *dabB* | HVO_0988 | *nuoN* | 25% | up to pos 322 of 494; up to pos 368 of 506 |
| 9k | HVO_2411 | *dabB* | HVO_1069 | *mrpA* | 26% | up to pos 421 of 494; up to pos 454 of 801 |
| 9k | HVO_2411 | *dabB* | HVO_1066 | *mrpD1* | 24% | up to pos 359 of 494; up to pos 409 of 565 |

**Table S10: Listing of all proteins mentioned in the text and in Suppl. Text S1.** This table lists all haloarchaeal proteins which are mentioned in the manuscript. Proteins may be listed more than once. (1) The column Section refers to the section in the Results and in Suppl. Text S1. As an example, 2c covers topic (c) from the decimal-numbered Results subsection 3.2 Amino Acid Biosynthesis. In Suppl. Text S1, this is covered under Section S2 subsection S2.c. Table 2 lists proteins from this section. For a few proteins, two sections are indicated (e.g. 1a/1b). (2) The two columns headed Code refer to proteins by their locus tag. Nearly all codes in the first Code column are from *Haloferax volcanii* (HVO). The second Code column "(other species or paralog)" contains information only when a paralog or a haloarchaeal homolog is indicated. The correlation between genome and locus tag prefix is available from Table S3. Protein complexes are represented in this table. Subunits that belong to the same complex are listed in consecutive rows which are highlighted in grey. If adjacent rows contain subunits of distinct protein complexes, the highlighting toggles between light grey and dark grey. If subunits of complexes are listed more than once, then the complex is only indicated at first listing. (3) The two columns headed Gene list the assigned gene for the corresponding locus_tag in the left adjacent Code column, or a dash if no gene has been assigned. (4) The column headed %seq_id indicates the protein sequence identity in case of paralogs or haloarchaeal homologs. (5) The column headed Comment provides various types of additional information.

| Species | Strain | Collection | Accessions | Locus tag prefix | Reference | PMID | Contribution |
|---|---|---|---|---|---|---|---|
| *Halobacterium salinarum* | R1 | DSM 671 | AM774415-AM774419 | OE | (Pfeiffer et al., 2008) | 18313895 | fpf@oe: seq,anno |
| *Halobacterium salinarum* | NRC-1 | ATCC 700922 | AE004437<br>AE004438<br>AF016485 | VNG | (Ng et al., 2000)<br>(Ng et al., 1998) | 11016950<br>9847077 | fpf: 3rd party anno BK010829-BK010831 |
| *Halobacterium salinarum* | 91-R6 | DSM 3754 | CP038631-CP038633 | HBSAL | (Pfeiffer et al., 2019)<br>(Pfeiffer et al., 2020) | 31296677<br>31797576 | fpf: seq,anno |
| *Halobacterium hubeiense* | JI20 | HAMBI 3616 | LN831302-LN831305 | Hhub | (Jaakkola et al., 2016) | 26628271 | fpf: anno |
| *Haloferax volcanii* | DS2 | ATCC 29605 | CP001953-CP001957 | HVO | (Hartman et al., 2010) | 20333302 | fpf: anno |
| *Haloferax gibbonsii* | LR2-5 | - | CP063205-CP063208 | HfgLR | (Tittes et al., 2021) | 33664716 | fpf: anno |
| *Haloferax mediterranei* | R-4 | ATCC 33500 | CP001868-CP001871 | HFX | (Han et al., 2012) | 22843593 | none |
| *Natronomonas pharaonis* | Gabara | DSM 2160 | CR936257-CR936259 | NP | (Falb et al., 2005) | 16169924 | fpf@oe: seq,anno |
| *Natronomonas moolapensis* | 8.8.11 | DSM 18674 | HF582854 | Nmlp | (Dyall-Smith et al., 2013) | 23516216 | fpf@oe: seq,anno |
| *Haloquadratum walsbyi* | HBSQ001 | DSM 16790 | AM180088-AM180089 | HQ | (Bolhuis et al., 2006) | 16820047 | fpf@oe: seq,anno |
| *Haloquadratum walsbyi* | C23 | DSM 16854 | FR746099-FR746102 | Hqrw | (Dyall-Smith et al., 2011) | 21701686 | fpf@oe: seq,anno |
| *Natrialba magadii* | MS3 | ATCC 43099 | CP001932-CP001935 | Nmag | (Siddaramappa et al., 2012) | 22559199 | fpf: anno |
| *Haloarcula marismortui* | - | ATCC 43049 | AY596290-AY596298 | rrnAC<br>rrnB<br>pNG | (Baliga et al., 2004) | 15520287 | none |
| *Haloarcula hispanica* | Y-27 | ATCC 33960 | CP002921-CP002923 | HAH | (Baliga et al., 2004) | 21994921 | none |
| *Halorubrum lacusprofundi* | ACAM 34 | ATCC 49239 | CP001365-CP001367 | Hlac | (Baliga et al., 2004) | 27617060 | none |
| *Halohasta litchfieldiae* | tADL | DSM 22187 | - | halTADL | (Demaere et al., 2013) | 24082106 | none |

**Table S11: Listing of the genomes which are manually curated and kept up to date.** Each row lists one genome. (1) Three columns provide information about the biological source of the genome. These are the column Species, the column Strain, and the column Collection which reports a culture collection and the number in that collection. (2) The column Accessions lists the accession numbers in the GenBank and EMBL nucleotide sequence databases. (3) The column Locus tag prefix provides the term which precedes the underscore and serial number of the locus tag for that genome. This information can be used to back-translate from a locus tag to a genome and the strain and species from which the genome originates. (4) The reference describing genome sequencing is provided in two columns. (4A) The column Reference links to the reference list. (4B) The column PMID lists the PubMed ID of the publication. (5) The column Contribution provides information about the contribution of one of us (F.P., tagged fpf) to genome sequencing and genome annotation. The term “fpf@oe” is used for genomes which have been sequenced in department Oesterhelt at the Max-Planck-Institute of Biochemistry. The term “seq_anno” indicates participation in genome sequencing and genome annotation, “anno” indicates participation in genome annotation. “none” indicates that the genome has been sequenced and annotated independently and has been adopted for manual curation independent from the original sequencing and annotation consortium. A species case is the genome of *Halobacterium salinarum* strain NRC-1, which has been sequenced and annotated independently, but our reannotation has been submitted to GenBank as a third party annotation.
